# A spatial single-cell atlas of the claustro-insular region uncovers key regulators of neuronal identity and excitability

**DOI:** 10.1101/2024.11.05.622016

**Authors:** Leon Fodoulian, Madlaina Boillat, Marie Moulinier, Alan Carleton, Ivan Rodriguez

## Abstract

The claustro-insular region is an evolutionarily conserved and extensively interconnected brain area, critical for functions such as attention, cognitive flexibility, interoception, and affective processing. Despite its importance, its cellular composition and organization remain poorly characterized, hindering a comprehensive understanding of the mechanisms underlying its diverse functions. By combining single-cell RNA sequencing and spatial transcriptomics, we created a high-resolution atlas of this region in mice, uncovering distinct neuronal subtypes and unexpected complexity. Leveraging this atlas, we investigated the role of NR4A2, a neuropsychiatric risk factor expressed in several claustro-insular neuronal subtypes. In an *Nr4a2* haploinsufficiency model, we found that only claustrum neurons exhibited shifts in molecular identity. This identity shift, which involved the activation of a transcription factor cascade, was associated with alterations in neuronal firing activity. Our findings provide new insights into the cellular architecture of the claustro-insular region and highlights *Nr4a2* as a master regulator of its component’s identities.

## Introduction

The claustro-insular region of the mammalian brain is an evolutionarily conserved bilateral complex composed of multiple structures. The insula, located laterally, is a neocortical area involved in diverse functions, such as interoception, salience detection, pain processing, and regulation of affective states^1–3^. The claustrum (CLA), situated medially adjacent to the external capsule, is an intricate subcortical nucleus characterized by dense, reciprocal connections with nearly all cortical areas^4–6^. This peculiar feature, unique to the CLA, suggests a key role in various higher-order cognitive functions^7^, and a growing body of research has already implicated the CLA in processes such as attention modulation, cognitive flexibility, and sleep regulation, among others (reviewed in^8–11)^. Despite increasing interest in the CLA, progress in understanding this structure has been hindered by anatomical challenges, particularly in rodent species where the CLA boundaries with the adjacent insula remain poorly outlined.

Visual examination of spatial gene expression patterns in the mouse claustro-insular area reveals an unusually complex landscape of cellular composition and organization^12^. Several studies have, sometimes indirectly, profiled the gene expression of CLA projection neurons at single-cell resolution^13–24^. Most identified a single transcriptomically defined population; however, in rare instances, two separate populations were distinguished^19,24^, which were suggested to correspond to a proposed subdivision of the CLA into a core and a shell^19^. The observation of a relatively homogenous CLA cell population at the transcriptomic level is counterintuitive, especially considering that numerous CLA subpopulations have been defined based on their electrophysiological properties^25–28^ or on their projection targets in the brain^6,29,19,30–32^, and that CLA neurons have been shown to have both inhibitory and excitatory effects in the cortex^17,33^. The discrepancy between transcriptomic homogeneity and functional diversity underscores the need for deeper investigation into the cellular architecture of the CLA. The thorough characterization and spatial mapping of claustro-insular cell types are essential for further elucidating the complex roles of this brain region.

CLA and dorsal endopiriform (EP) neurons, broadly defined, are characterized by their early and sustained transcription of *Nr4a2* (also called *Nurr1*), a canonical gene marker of the CLA, coding for the NR4A2 transcription factor^34–39^. In the dopaminergic system, NR4A2 plays a critical role in determining neuronal identity^40–43^ and has been linked to various neurodevelopmental diseases^44–51^. In the CLA, the role played by *Nr4a2* remains elusive.

To fill the gap in our understanding of the identity and organization of cells in the claustro-insular region, we sought to generate a high-resolution map of its various cell types by combining single-cell RNA sequencing (scRNAseq) with spatial transcriptomics. Furthermore, given the early and persistent expression of *Nr4a2* in the CLA and surrounding cell types, and considering its crucial role in establishing and maintaining cellular phenotypes in other brain regions, we investigated its function in defining the cellular identities of claustro-insular neurons.

## Results

### Transcriptomic characterization of cell types in the claustro-insular region

To characterize the cellular composition and organization of the claustro-insular region, we performed a dual analysis by combining scRNAseq and multiplexed error-robust fluorescence in situ hybridization (MERFISH) spatial transcriptomics (Fig. 1a). First, we performed scRNAseq on isolated cells from claustro-insular microdissections, covering a substantial portion of the CLA along the anteroposterior axis (bregma +1.545 to −1.055). In total, we recovered 67’226 cells encompassing various neuronal and non-neuronal cell types (Supplementary Fig. 1a-c). To identify the neuronal cell types populating the claustro-insular region, we filtered for high-quality neurons, clustered them, and computed their marker genes. This analysis revealed the presence of CLA projection neurons and known adjacent cortical and subcortical cell types, including deep-layer L5b, L6a and L6b neurons, upper-layer L2/3 neurons, piriform cortex neurons (pir), a heterogeneous group of interneurons (IN) and several groups of medium spiny neurons (MSN) of the striatum (Fig. 1b, Supplementary Fig. 1d-f).

**Fig. 1.**
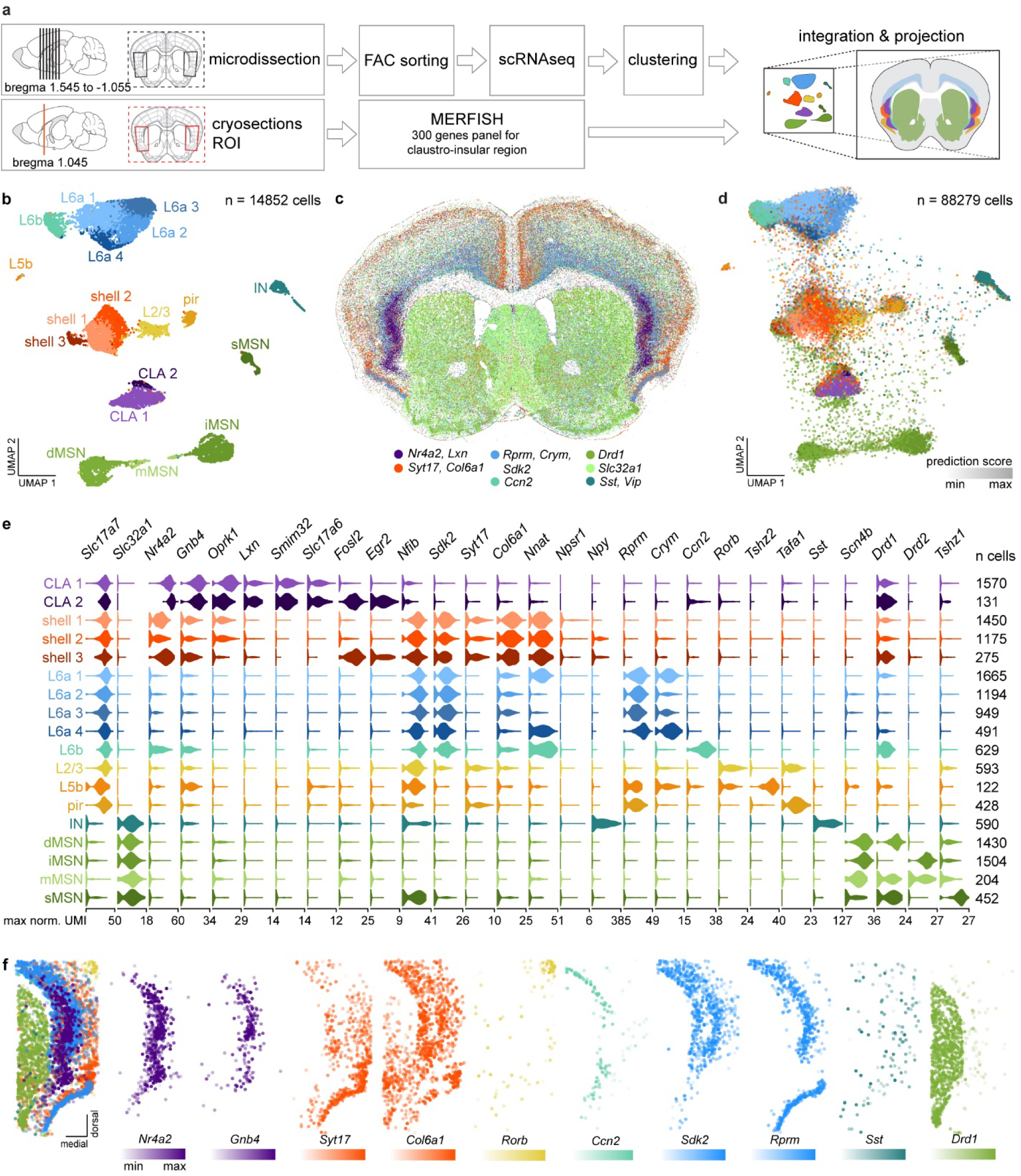
High-resolution characterization of cell types in the claustro-insular region. **a** Schematic of the experimental design for high-resolution characterization of the claustro-insular region. **b** Visualization of neuronal cell types from the scRNAseq dataset on a Uniform Manifold Approximation and Projection (UMAP) plot. CLA, claustrum projection neuron; dMSN, striatal direct pathway medium spiny neuron; iMSN, striatal indirect pathway medium spiny neuron; IN, interneuron; L2/3, layer 2/3 glutamatergic neuron; L5b, layer 5b glutamatergic neuron; L6a, layer 6a glutamatergic neuron; L6b, layer 6b glutamatergic neuron; mMSN, mixed striatal medium spiny neuron; pir, piriform glutamatergic neuron; shell, shell projection neuron; sMSN, striosome medium spiny neuron. n = 13 mice. **c** Visualization of expression of cell type-specific marker genes on a representative coronal section of a mouse brain after MERFISH labeling and imaging. **d** Visualization of neuronal cell types from the MERFISH dataset on a UMAP plot. Cell type attributes of cells were obtained after mapping the MERFISH data onto the scRNAseq dataset. n = 7 mice. **e** Violin plots showing cell type-specific distribution of expression of cell type-specific marker genes in the scRNAseq dataset (normalized data are shown in log_10_ scale). **f** Visualization of cell type-specific expression of cell type-specific marker genes on a representative coronal section of the claustro-insular region from the MERFISH dataset. Cells expressing at least 5 mRNA molecules of the corresponding gene are plotted. Colors indicate the cell type to which the genes are specific.

We identified two subtypes of CLA excitatory neurons, namely CLA 1 and CLA 2. Both subtypes exhibit high transcription levels of *Nr4a2* and other canonical CLA markers, including *Gnb4*, *Oprk1*, *Lxn*, *Slc17a6* and *Smim32*, a recently identified marker of the CLA^52^ (Fig. 1e). The CLA 2 subtype is further distinguished by the expression of immediate early genes such as *Fosl2* and *Egr2*. Additionally, we identified four subtypes of L6a neurons, all characterized by high expression levels of *Nfib*, *Sdk2* and *Rprm*, with more variable expression levels of *Nnat* and *Crym*, which define the distinct subtypes (Fig. 1b,e).

In addition to these cell types, we identified a large cluster of cells with a unique transcriptomic identity, intermediate between a CLA-like and a cortical identity, which we refer to as shell projection neurons (or “shell” for short) (Fig. 1b,e, Supplementary Fig. 1f). We identified three subpopulations of shell neurons, all characterized by low expression levels of *Nr4a2*, *Gnb4* and *Oprk1*, high expression levels of *Nfib*, *Col6a1* and *Nnat*, low but relatively specific expression of *Syt17*, sparse expression of *Npsr1*, and no expression of *Slc17a6*, *Smim32* or L6a markers such as *Rprm* and *Crym* (Fig. 1e). Two of these subpopulations, shell 1 and shell 2, differ in their *Nr4a2* expression levels, with the shell 2 subtype further distinguished by low levels of *Npy* expression. The third subtype, shell 3, exhibits expression of immediate early genes, similar to CLA 2. To localize the identified cell types and subtypes within the claustro-insular region, we designed a custom panel of 300 genes enriched for markers specific to the various populations described above and performed MERFISH spatial transcriptomics (Fig. 1a,c, Supplementary Tables, MERFISH gene panel). Imaging multiple coronal sections at bregma +1.045 revealed robust expression of these cell type markers (Fig. 1c). To identify cell populations in the MERFISH data, we performed a reference-based mapping of the data onto the scRNAseq dataset^53^, successfully detecting most of the previously identified cell types and subtypes (Fig. 1d). Many of these populations consisted of cells with high prediction and mapping scores, indicating high confidence in cell type assignment and accurate representation by the scRNAseq reference dataset, respectively (Supplementary Fig. 2). Visualization of the expression levels of cell type marker genes confirmed their spatial specificity within the claustro-insular region, highlighting the complex organization and intermingling of cell types in this brain region (Fig. 1f).

### Generation of a map of the claustro-insular region

The complexity of gene expression patterns observed in MERFISH sections suggests that drawing discrete, non-overlapping boundaries to map the claustro-insular region is not an ideal approach to represent its brain structures. Instead, we aimed to create a map that captures this complexity while providing insights into the relative localization of each component of the claustro-insular region. To achieve this, we employed a data-driven approach by staining sections for gene expression, identifying cell types based on their gene expression profiles, and delineating outlines that reflect the 2-dimensional density distribution of these cell types across the sections. We labeled multiple coronal sections spanning four anteroposterior levels of the CLA for the expression of five key gene markers using single-molecule fluorescence in situ hybridization (smFISH). These markers enabled us to distinguish four major components of the claustro-insular region: *Nr4a2* for CLA, shell and L6b neurons, *Slc17a6* for CLA neurons, *Ccn2* for L6b neurons, *Rprm* for L6a neurons, and *Syt17* for shell neurons (Fig. 2a-c).

**Fig. 2.**
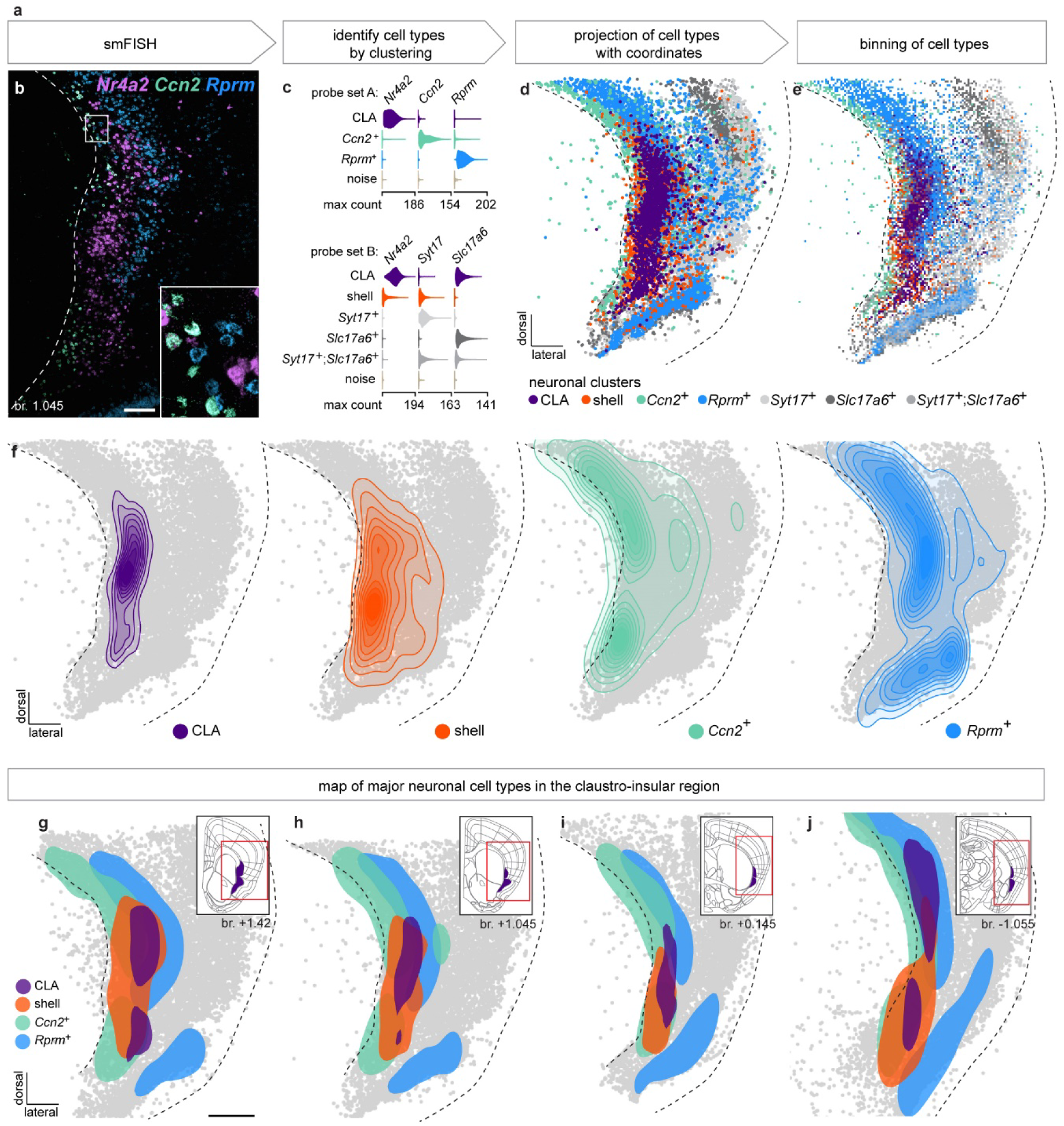
Map of the claustro-insular region. **a** Schematic of the data processing approach. **b** Representative image of a coronal section (bregma +1.045) of the claustro-insular region labeled by smFISH with probes for *Nr4a2* (purple), *Ccn2* (cyan) and *Rprm* (blue). The white square highlights the region magnified in the inset. The dotted line highlights the border of the external capsule. Scale bar: 200 μm. **c** Violin plots showing the distribution of expression of marker genes across identified cell types in the smFISH data (raw data are shown in linear scale). Probe set A: n = 2 mice, 15 sections, 91’634 clustered cells; Probe set B: n = 2 mice, 15 sections, 80’627 clustered cells. **d** Spatial organization of neuronal cell populations identified in the smFISH dataset. Multiple images are aligned and cells are projected on the same framework. n = 2 mice, 8 sections. **e** Proportional visualization of the claustro-insular region, binned into 20 μm bins. Bin colors are weighted by the relative proportion of neuronal cell types within that bin. n = 2 mice, 8 sections. **f** 2D density maps showing the spatial distribution of neuronal cell populations within the claustro-insular region. The total area comprises 100% of the 2D kernel density estimation of the distribution of all cells within the respective population. n = 2 mice, 8 sections. **g-j** Maps of the claustro-insular region at four different antero-posterior levels. Colored areas represent the highest density region of cell types at a 50% probability level of the two-dimensional kernel density estimate. Inset shows the representation of the CLA and endopiriform nucleus (EP) in the Allen Mouse Brain Atlas (mouse.brain-map.org), at the corresponding antero-posterior position. Red squares highlight the region imaged after smFISH labeling for map generation. Scale bar: 500 μm. n = 2 mice, 6-8 sections/bregma level.

We chose smFISH for its high quantitative resolution, enabling us to capture the large variability in expression levels of these marker genes (Supplementary Fig. 3). After staining, we clustered the cells, identified the cell types, aligned images from the same anteroposterior level, and projected the cells’ coordinates onto a reference image to create 2D maps of cell type localization at each brain level (Fig. 2d, Supplementary Fig. 4a). To highlight the diversity of cell types in the claustro-insular region, we created proportional visualizations of the maps (Fig. 2e and Supplementary Fig. 4b). We binned the 2D spatial maps into 20 μm bins, and the proportion of each cell type in each bin was used to assign a color weighted by these proportions. To determine the spatial distribution of these cell types within the claustro-insular region, we generated 2D density maps for CLA, shell, *Ccn2*+ neurons (labeling mainly L6b), and *Rprm*+ neurons (labeling mainly L6a, L5b and pir) at each brain level (Fig. 2f, Supplementary Fig. 4c). The final maps display the highest density regions of CLA, shell, *Ccn2*+, and *Rprm*+ neurons at the 50% probability level of their respective two-dimensional kernel density estimates (Fig. 2g-j). These maps were more versatile than conventional brain atlases, allowing for overlapping delineations of brain structures. Notably, the boundaries of CLA and shell overlapped substantially with each other and with cortical L6a and L6b (Fig. 2g-j). This approach proved essential for accurately characterizing the highly complex claustro-insular region, where cell types are closely juxtaposed and intermingled.

We leveraged this map to localize each cell type and subtype identified by scRNAseq within the brain. As expected, well-defined cell types from the MERFISH dataset, located within or near the boundaries of the claustro-insular region, were accurately mapped (Supplementary Fig. 5). We then focused on characterizing the spatial localization of CLA, shell and L6a subtypes (Fig. 3a). Since MERFISH may not be accessible to all researchers, we compared these results with a simpler approach for targeting cell subtypes using smFISH, which employs combinations of only 2-3 gene to identify cell subpopulations (Figure 3b-i). By assessing the presence or absence of specific marker genes, we were able to match subpopulations from the smFISH data to those identified in the scRNAseq dataset. A strong correspondence was observed between the localization of cell types and subtypes identified by smFISH and MERFISH (Fig. 3b-i, Supplementary Fig. 6), particularly for cells with high prediction score in the MERFISH dataset. While shell and L6a subpopulations displayed distinct localization patterns, CLA 1 and CLA 2 subtypes were intermingled. Moreover, shell subtypes were consistently identified across sections from different animals and exhibited stereotypical spatial localizations across mice (Supplementary Fig. 6), similar to CLA and L6a subpopulations. Together, these results demonstrate the utility of the claustro-insular map we generated and showcase the power of combining spatial transcriptomics with scRNAseq for the high-resolution characterization of this complex brain region.

**Fig. 3.**
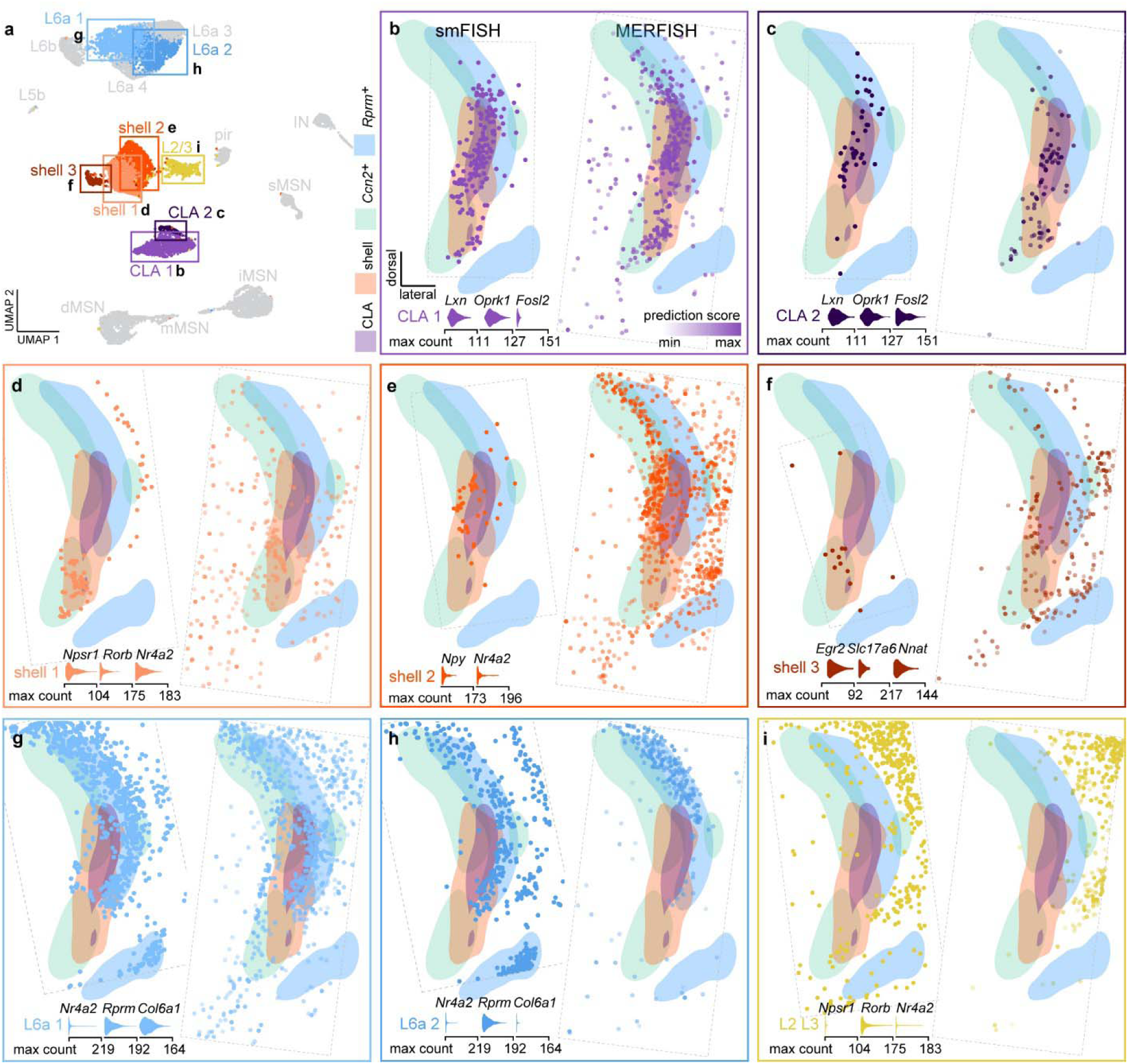
Spatial organization of cell types and subtypes in the claustro-insular region. **a** UMAP plot highlighting the cell types or subtypes whose localization was determined by smFISH and MERFISH spatial transcriptomics. **b-i** Spatial organization of different neuronal types and subtypes projected on the map of the claustro-insular region at bregma +1.045. *Left*: cells from smFISH datasets. Violin plots below the maps show the distribution of expression of marker genes used for cell type and subtype identification in the smFISH data (raw data are shown in linear scale). *Right*: cells from MERFISH dataset. Prediction scores highlight the confidence of cell type attribution in the MERFISH data after reference-based mapping onto the scRNAseq dataset. Dotted lines highlight the border of the image or of the ROI used for cell selection. Cells from one representative section are shown for each cell type and subtype.

### *Nr4a2* haploinsufficiency induces transcriptomic alterations in CLA projection neurons

We next aimed to investigate how the transcriptomic identity of CLA neurons is established. Noting the variable levels of *Nr4a2* expression across different subpopulations of the claustro-insular region, particularly in CLA and shell subtypes (Fig. 1e), and given the early developmental onset of *Nr4a2* expression in CLA neurons^34,35^, we hypothesized that this transcription factor may regulate gene expression within these subpopulations. To test this, we used *Nr4a2* heterozygous mice (*Nr4a2^del/wt^*) carrying one non-functional copy of *Nr4a2*, and compared them to their wild-type littermates (*Nr4a2^wt/wt^*). smFISH quantification of *Nr4a2* mRNA transcripts in the claustro-insular region showed a 2.6-fold reduction of *Nr4a2* expression in heterozygous mice (Fig. 4a-c). To assess the impact of *Nr4a2* heterozygosity on gene expression in the claustro-insular region, we compared the transcriptomes of the various cell types and subtypes in the scRNAseq dataset from *Nr4a2^wt/wt^* mice to those from *Nr4a2^del/wt^* mice. CLA projection neurons, particularly the CLA 1 subtype, exhibited a substantial number of differentially expressed genes, while other cell types appeared to be less affected (Fig. 4d, Supplementary Fig. 7). The CLA 2 subtype had fewer cells represented in the dataset, likely contributing to the low number of significantly modulated genes. Among the differentially expressed genes, we identified key marker genes (e.g. *Cdh13*, *Rxfp1*, *Nr2f2*, *Syt17* and *Ntm*) as well as genes coding for ion channels that could influence neuronal function (e.g. *Scn1b* and *Ryr2*) (Fig. 4e,f, Supplementary Fig. 8). This strong modulation of gene expression in *Nr4a2*-expressing cells was further confirmed by smFISH quantification of selected up- and downregulated genes in haploinsufficient mice (Fig. 4g,h, Supplementary Fig. 9).

**Fig. 4.**
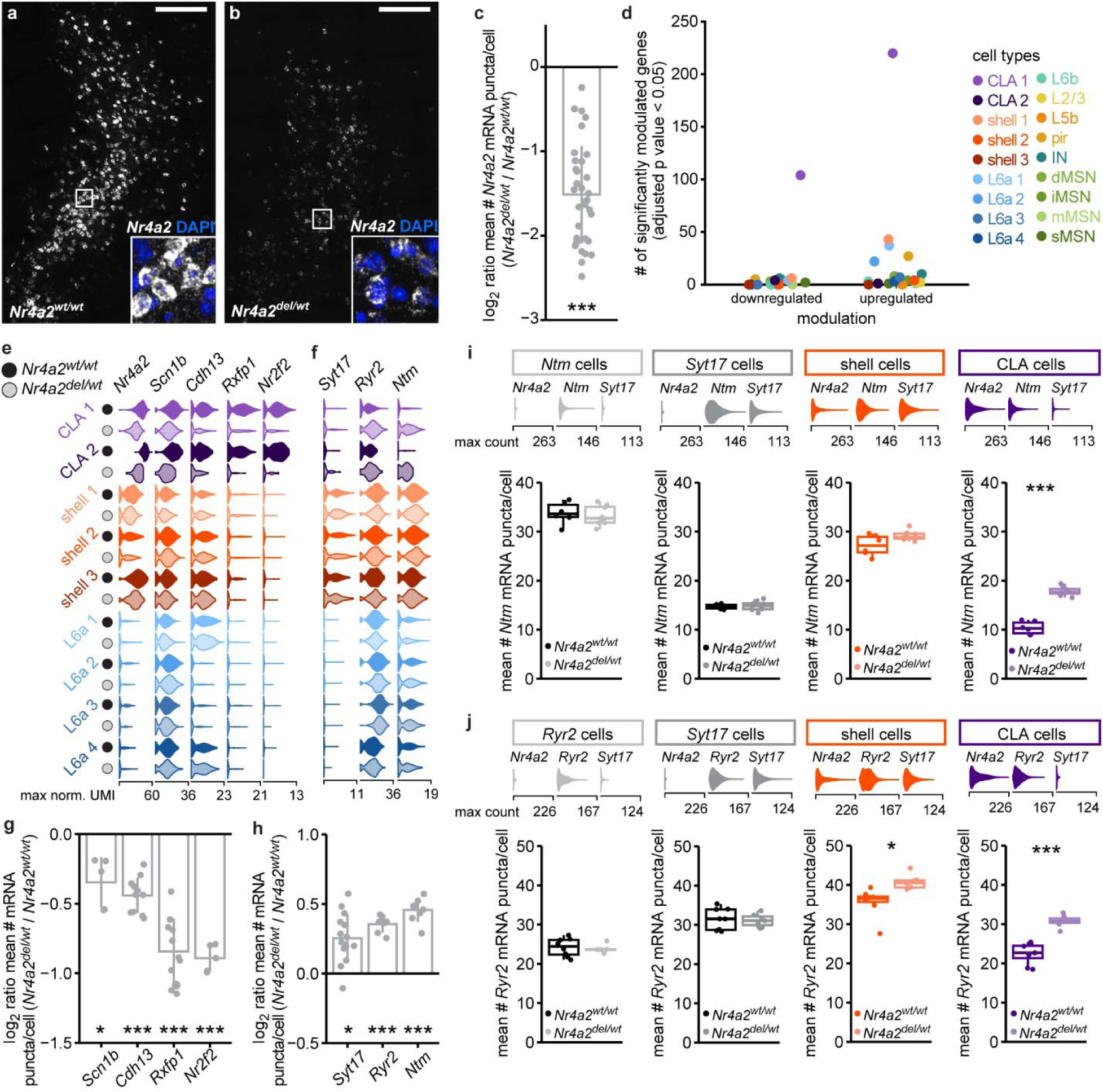
Transcriptomic perturbations in *Nr4a2* haploinsufficient mice. **a,b** Images of coronal sections of the CLA from an *Nr4a2^wt/wt^* mouse (**a**) and an *Nr4a2^del/wt^* mouse (**b**) with *Nr4a2* mRNA transcripts labeled by smFISH (white). The white square highlights the region magnified in the inset. DAPI staining of the nuclei (blue). Scale bar: 200 μm. **c** Mean number of *Nr4a2* mRNA puncta detected per cell in the claustro-insular area. Each dot represents the mean of all cells from one section of an *Nr4a2^del/wt^* mouse relative to the mean of all sections of *Nr4a2^wt/wt^* mice processed in the same batch, and the bar represents the mean ± SD. n = 4 *Nr4a2^wt/wt^* mice, 26 sections, 55’697 cells; n = 4 *Nr4a2^del/wt^* mice, 34 sections, 71’389 cells. *** p<0.001; chi-square-based LRT applied to a negative binomial GLMM with quadratic parametrization. **d** Total number of significantly modulated genes in *Nr4a2^del/wt^*mice in each cell type from the scRNAseq dataset. n = 5 *Nr4a2^wt/wt^* mice, n = 8 *Nr4a2^del/wt^* mice. **e,f** Violin plots illustrating the distribution of expression for downregulated (**e**) and upregulated (**f**) genes across claustro-insular neuronal populations from *Nr4a2^wt/wt^* and *Nr4a2^del/wt^* mice in the scRNAseq dataset (normalized data are shown in log_10_ scale). **g,h** Mean number of mRNA puncta of downregulated (**g**) and upregulated (**h**) genes detected per cell in *Nr4a2^+^* and/or *Oprk1^+^* cells from the claustro-insular region of *Nr4a2^wt/wt^*and *Nr4a2^del/wt^* mice. Each dot represents the mean of cells from one section of an *Nr4a2^del/wt^* mouse relative to the mean of all sections of *Nr4a2^wt/wt^*mice processed in the same batch, and bars represent the mean ± SD. n = 2-3 *Nr4a2^wt/wt^* mice, 4-14 sections; n = 2-3 *Nr4a2^del/wt^* mice, 5-15 sections. * p<0.05, *** p<0.001; chi-square-based LRT applied to a negative binomial GLMM with quadratic parametrization; *p*-values were adjusted for multiple comparisons using the Holm method. **i,j** Cell type-specific modulation of *Ntm* (i) and *Ryr2* (j) gene expression. *Top*: Violin plots at the top of each panel show the distribution of expression of cell marker genes used for cell type identification in the smFISH data (raw data are shown in linear scale). *Bottom*: Box plots showing the distribution of *Ntm* (i) and *Ryr2* (j) expression levels within the different identified cell populations from *Nr4a2^wt/wt^* and *Nr4a2^del/wt^* mice in the smFISH dataset. Each dot represents the mean expression level of all cells from one section. Box limits represent Q1 to Q3, the line represents the median, error bars represent ±1.5*IQR. n = 2 *Nr4a2^wt/wt^* mice, 6-8 sections, n = 2 *Nr4a2^del/wt^* mice, 6-8 sections. * p<0.05, *** p<0.001; chi-square-based LRT applied to a negative binomial GLMM with quadratic parametrization; *p*-values were adjusted for multiple comparisons using the Holm method.

The scRNAseq analyses also suggested that CLA and shell neurons were not similarly affected by the reduction in *Nr4a2* mRNA levels (Fig. 4e,f). To explore this further, we clustered cells from the smFISH experiments to differentiate CLA from shell neurons based on the expression patterns of *Nr4a2* and *Syt17*, and evaluated the expression of two genes that showed strong upregulation in *Nr4a2^del/wt^*mice (Fig. 4i,j). *Ntm* and *Ryr2* were either not or only slightly modulated in cells lacking CLA identity, indicating that *Nr4a2* heterozygosity primarily affects transcription in CLA neurons, with no or minimal effects on neighboring shell and other cell populations.

### *Nr4a2* haploinsufficiency perturbs CLA projection neuron firing

Since multiple genes encoding ion channels, neurotransmitter receptors, and related signaling proteins were dysregulated by *Nr4a2* haploinsufficiency (Fig. 5a and Supplementary Fig. 10a), we investigated whether the transcriptomic changes led to alterations in spiking behavior. Whole-cell patch-clamp recordings of mPFC-projecting CLA neurons (Fig. 5b) revealed reduced firing frequency in *Nr4a2^del/wt^*cells (Fig. 5c-e), with no change in resting membrane potential (Fig. 5f). This hypoexcitability persisted even in the presence of blockers for fast GABAergic and glutamatergic synaptic transmission (Fig. 5d,e), suggesting intrinsic cellular changes. Action potential (AP) analysis showed that while rise time and amplitude were unaffected, the decay time and half-width were shorter (Fig. 5g,h), which usually is typically indicative of higher excitability. Although several genes encoding sodium and potassium channels were dysregulated in *Nr4a2* haploinsufficient mice, pharmacological targeting of these channels failed to restore WT-like excitability (see Methods and Supplementary Fig. 10b-f). This suggests that the observed changes in firing properties cannot be easily explained by alterations in these channels alone. In contrast, we observed a 25% increase in afterhyperpolarization potential (AHP) amplitude - an important regulator of firing frequency - in *Nr4a2^del/wt^*neurons (Fig. 5g,h). During the action potential, the opening of voltage-gated calcium channels (VGCCs) allows calcium influx into the cytoplasm, activating various calcium-activated potassium channels (such as big and small conductance potassium channels, BK and SK, respectively), which regulate different phases of the AHP^54–56^. BK channels (encoded by *Kcnma1*), responsible for the fast AHP^54^, were highly transcribed in CLA neurons but showed no significant differences in expression in haploinsufficient neurons (Supplementary Fig. 10). SK channels (encoded by *Kcnn1-3*) were expressed at very low levels in both genotypes (Supplementary Fig. 10a). Interestingly, blocking VGCC normalized excitability and AHP amplitude (Fig. 5i–l).

**Fig. 5.**
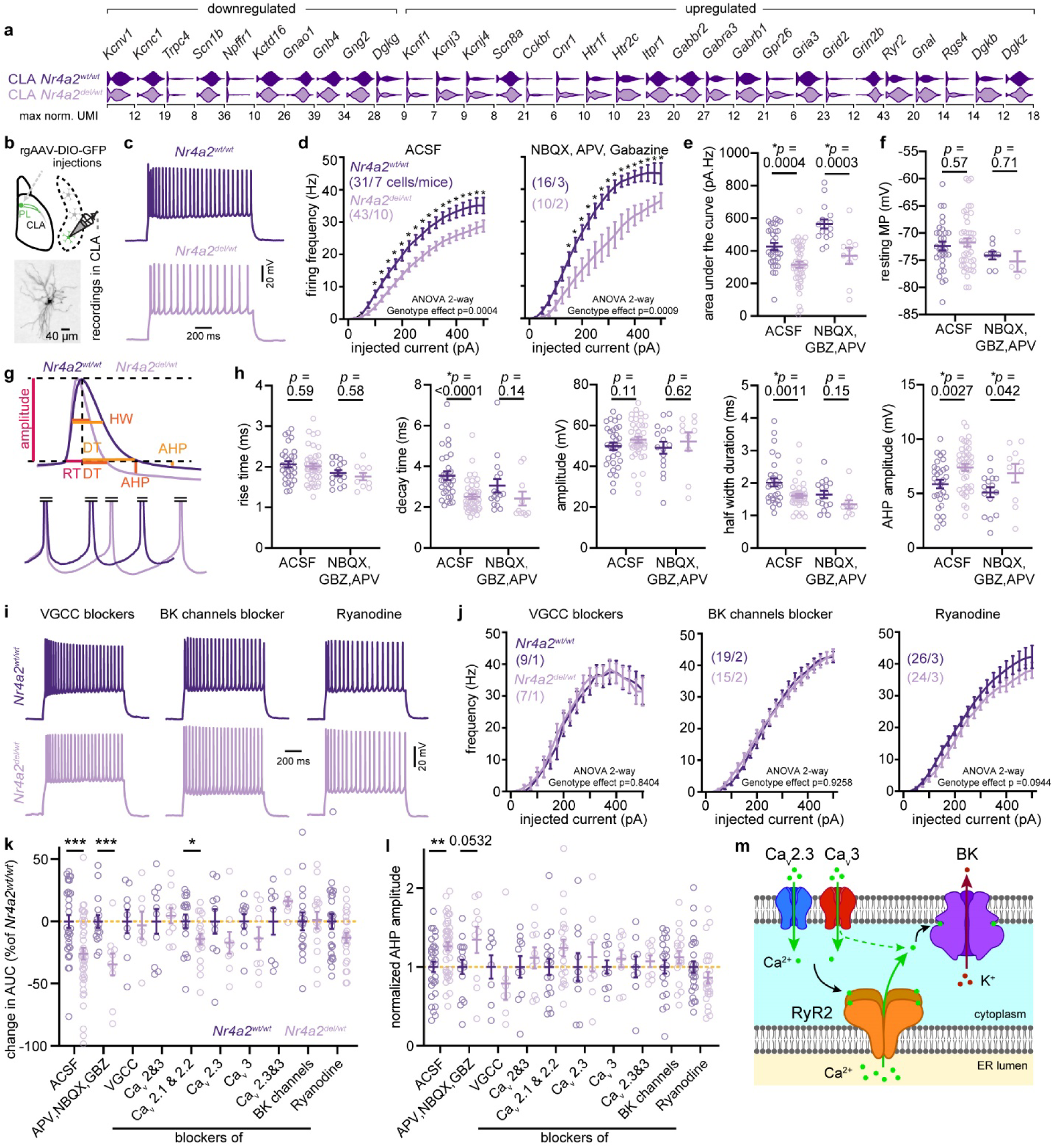
*Nr4a2* haploinsufficiency suppresses CLA neuron firing via enhanced recruitment of Ca^2+^-activated potassium channels. **a** Significantly modulated genes encoding channels, neurotransmitter receptors and related signaling proteins in CLA cells of *Nr4a2^del/wt^*mice when compared to wild-type littermates. **b** Schema of the experimental procedure. Retrogradely labeled CLA neurons projecting to the mPFC were recorded in brain slices. *Bottom*: example of a patched cell filled with biocytin. **c** Example of the spiking activity evoked by a 1s depolarizing step of current (250 pA). **d** Input/Output functions of CLA neurons recorded either in artificial cerebrospinal fluid (ACSF) or in blockers of the fast glutamatergic (NBQX and APV) and GABAergic (Gabazine) synaptic transmission (* indicates significant post-hoc testing using Fisher’s LSD test with a 5% false discovery rate; *p*-values were adjusted for multiple comparisons using the Benjamini-Hochberg false discovery rate method). **e** Area under the curve (AUC) of the input/output function for each *Nr4a2^wt/wt^*and *Nr4a2^del/wt^* cell computed in ACSF and synaptic blockers. **f** Resting membrane potential recorded at the opening of cells of *Nr4a2^wt/wt^*and *Nr4a2^del/wt^* mice in both ACSF and synaptic blockers. **g** Examples of recorded action potentials (AP, note that the AP were truncated for the bottom traces). HW, half width; RT, rise time; DT, decay time; AHP, after hyperpolarization potential. **h** Quantifications of the different AP parameters (each circle represents the mean parameter value computed on all spikes of all steps for a given cell). **I** Example of the firing frequency evoked by a one-second 250 pA depolarizing current step under various pharmacological conditions. **j** Input/Output functions of CLA neurons recorded in the presence of VGCC blockers (10 µM nifedipine, 100 nM ω-agatoxin IVA, 3 µM ω-conotoxin GVIA, 400 nM SNX-482, 2 µM ML218), BK channels blocker (100 nM iberiotoxin) and RyR2 channel blocker (10-20 µM ryanodine). **k** Comparison of the firing functions between *Nr4a2^wt/wt^* and *Nr4a2^del/wt^* cells, expressed as a percentage of the mean wild-type AUC calculated for each pharmacological condition (see Methods). Cav2&3: ω-Agatoxin IVA, ω-Conotoxin GVIA, SNX-482 and ML218. Cav2.1&2.2: ω-Agatoxin IVA and ω-Conotoxin GVIA. Cav2.3: SNX-482. Cav3: ML218. * p<0.05, *** p<0.001; Mann-Whitney U test. **l** Comparison of the relative AHP amplitude between *Nr4a2^wt/wt^*and *Nr4a2^del/wt^* cells under each pharmacological condition. ** p<0.01; Mann-Whitney U test. **m** Summary diagram of the mechanism controlling changes in firing frequency through the activation of BK channels. Data are shown as mean ± SEM. In **e,f,h,k** and **l,** each circle represents one recorded neuron. See Supplementary Tables, Statistics table for more details.

Despite no transcriptomic alterations in their expression, Cav2.3 and Cav3 VGCC subunits were identified as contributors to hypoexcitability (Fig. 5k,l and Supplementary Fig. 10g-k). Furthermore, blocking BK channels with iberiotoxin fully restored excitability (Fig. 5j–l). Potentially explaining the changes in BK-mediated firing, we found that the ryanodine receptor gene *Ryr2*, whose product is a key component of the calcium-induced calcium release (CICR) mechanism (Fig. 5m), was overexpressed in *Nr4a2^del/wt^* neurons (Fig. 4 and Fig. 5a), and that blocking ryanodine receptors significantly improved excitability and reduced AHP in *Nr4a2^del/wt^*neurons (Fig. 5l). Overall, our data suggest that the hypoexcitability observed in *Nr4a2^del/wt^*neurons arises primarily from RyR2 overexpression, which disrupts the CICR pathway in CLA neurons (Fig. 5m).

### The cellular composition of the claustro-insular region is altered in *Nr4a2* haploinsufficient mice

Given the alterations observed in other brain systems^40–43^ and the substantial transcriptomic changes in CLA identity genes in *Nr4a2^del/wt^* mice (Supplementary Fig. 8), we investigated whether these modifications could affect the cellular composition of the claustro-insular area. Our smFISH analyses showed that while the overall number of cells in the claustro-insular region remained unchanged (Fig. 6a), there was a 2.3-fold reduction in the number of cells exhibiting robust *Nr4a2* expression (defined as a minimum of 20 mRNA puncta/cell)(Fig. 6b). The scRNAseq dataset revealed a strong decrease in the proportion of cells classified as CLA 1 and CLA 2, with no significant differences noted in other cell types and subtypes between genotypes (Fig. 6c). This reduction was also evident in the MERFISH dataset (Fig. 6d,e). To further characterize the changes in cellular composition, we performed a similar analysis on smFISH data, confirming cell-type specific alterations (Fig. 6f). We aligned multiple images, overlaid cell identities onto a reference section, and generated a contrast map to highlight the relative proportions of CLA and shell cells within 20 µm bins (Fig. 6g). The resulting map revealed a notable reduction in CLA cell density, particularly in the core of the CLA and in the EP.

**Fig. 6.**
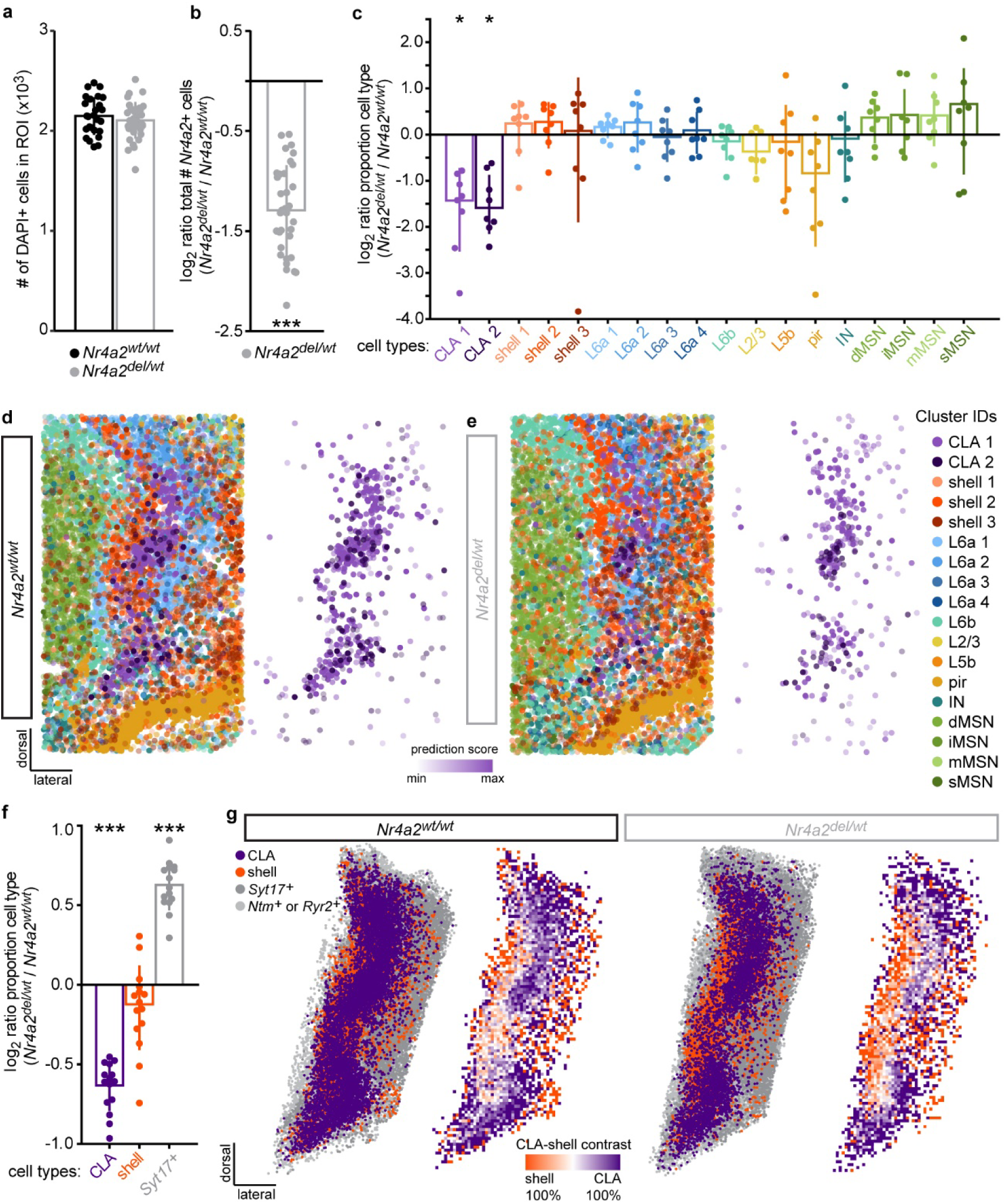
Alteration of cellular composition in the claustro-insular region of *Nr4a2* haploinsufficient mice. **a** Quantification of the total number of cells present in the ROI around the CLA in *Nr4a2^wt/wt^* and *Nr4a2^del/wt^* mice. Each dot represents the mean of one section, and barplots and error bars represent the mean ± SD of all sections. n = 4 *Nr4a2^wt/wt^* mice, 26 sections, 55’697 cells; n = 4 *Nr4a2^del/wt^* mice, 34 sections, 71’389 cells. **b** Quantification of the total number of *Nr4a2-*positive cells (minimum of 20 mRNA puncta/cell) within the ROI. Each dot represents the number of cells from one section of an *Nr4a2^del/wt^* mouse relative to the mean number of cells from all sections of *Nr4a2^wt/wt^* mice processed in the same batch, and the barplot and error bar represent the mean ± SD. *** p<0.001; chi-square-based LRT applied to a negative binomial GLMM with quadratic parametrization. **c** Ratio of the proportion of cell types from the scRNAseq dataset. Each dot represents the cell type proportion of an *Nr4a2^del/wt^* mouse relative to the mean proportion of that cell type of all *Nr4a2^wt/wt^* mice. Each dot represents one mouse, and barplots and error bars represent the mean ± SD. * p<0.05; moderated *t*-test with robust empirical Bayes variance shrinkage; *p*-values were adjusted for multiple comparisons using the Benjamini-Hochberg false discovery rate method. **d,e** Spatial organization of cell types and subtypes on a representative coronal section of the claustro-insular region of an *Nr4a2^wt/wt^* (**d**) and an *Nr4a2^del/wt^* (**e**) mouse from the MERFISH dataset. *Left:* all cell types and subtypes. *Right:* CLA 1 and CLA 2 cells. **f** Ratio of the proportion of cell types from the smFISH dataset. Each dot represents the cell type proportion of an *Nr4a2^del/wt^* mouse relative to the mean proportion of that cell type of all *Nr4a2^wt/wt^* mice. Each dot represents one section, and barplots and error bars represent the mean ± SD. *** p<0.001; moderated *t*-test with robust empirical Bayes variance shrinkage; *p*-values were adjusted for multiple comparisons using the Benjamini-Hochberg false discovery rate method. **g** *Left plots*: Spatial organization of neuronal cell populations identified in the smFISH dataset. Multiple images from *Nr4a2^wt/wt^* and *Nr4a2^del/wt^* mice are aligned and cells are projected on the same framework. *Right plots*: CLA/shell contrast maps of the claustro-insular region, binned into 20 μm bins. Bin colors correspond to the relative proportion of CLA or shell neurons within that bin. n = 2 *Nr4a2^wt/wt^* mice, 14 sections, n = 2 *Nr4a2^del/wt^* mice, 15 sections.

### *Nr4a2* is a master regulator of CLA and shell identities

*Nr4a2* haploinsufficient CLA projection neurons exhibited downregulation of CLA identity genes, including *Cdh13*, *Rxfp1* and *Nr2f2*, alongside upregulation of shell identity genes, such as *Syt17* and *Ntm* (Fig. 4e-j, Supplementary Fig. 8 and 9). To further explore the role of *Nr4a2* in establishing CLA projection neuron identity, we trained a linear classifier using only the modulated genes (Fig. 7a, Supplementary Fig.11). The expression pattern of these genes in wild-type cells allowed for accurate classification of CLA, shell and L6a neurons. Similarly, shell and L6a neurons from *Nr4a2^del/wt^*mice were classified with accuracy scores comparable to that of *Nr4a2^wt/wt^*cells. However, CLA neurons from haploinsufficient mice showed a significant decrease in classification accuracy, with nearly 18% of the cells being misclassified as shell projection neurons. These results suggest that *Nr4a2* differentially influences various transcriptional programs based on its expression levels.

**Fig. 7.**
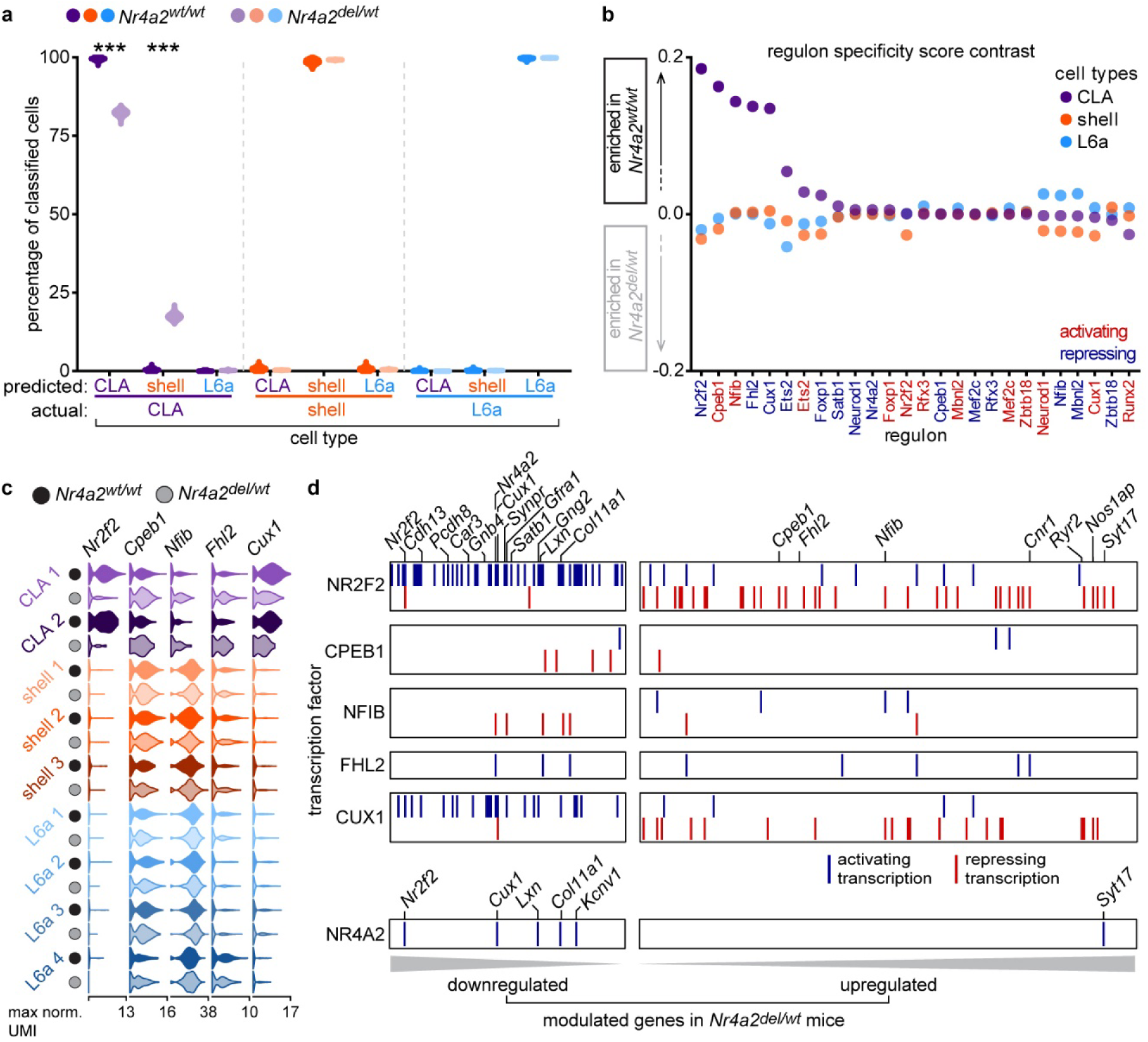
CLA and shell neuronal identities are driven by *Nr4a2*. **a** Cell type classification performance of CLA, shell and L6a neurons of *Nr4a2^wt/wt^*and *Nr4a2^del/wt^* mice from the scRNAseq dataset using a linear support vector classification algorithm. Violin plots show the distribution of classification outcomes across 1’000 iterations. *** p<0.001; chi-square-based LRT applied to a logistic regression model; *p*-values were adjusted for multiple comparisons using the Holm method. **b** Dot plot showing the contrast in regulon specificity scores between *Nr4a2^wt/wt^*and *Nr4a2^del/wt^* mice for CLA, shell and L6a cell types from the scRNAseq dataset. **c** Violin plots showing the distribution of downregulated and upregulated transcription factors expression across claustro-insular neuronal populations from *Nr4a2^wt/wt^* and *Nr4a2^del/wt^* mice in the scRNAseq dataset (normalized data are shown in log_10_ scale). **d** Raster plot showing transcription factors from enriched regulons and their target-modulated genes in *Nr4a2^del/wt^* CLA neurons.

Several transcription factors were modulated due to *Nr4a2* haploinsufficiency. To assess whether the shift in CLA identity was directly mediated by *Nr4a2*, we inferred gene regulatory networks (i.e. regulons) from the list of modulated genes, identifying 26 regulons involving 15 transcription factors (Fig. 7b). We characterized the regulons specific to each cell type and genotype by computing regulon specificity scores^57^ and calculated contrasts in these scores between genotypes. Notably, the Nr4a2 regulon did not show differential enrichement in CLA neurons from *Nr4a2^wt/wt^*and *Nr4a2^del/wt^* mice (Fig. 7b). However, five other regulons, involving *Nr2f2*, *Cpeb1*, *Nfib*, *Fhl2* and *Cux1*, displayed higher enrichment scores in *Nr4a2^wt/wt^* CLA projection neurons (Fig. 7b). Among these, *Nr2f2* and *Cux1* were downregulated due to the *Nr4a2* haploinsufficiency, while *Cpeb1*, *Nfib* and *Fhl2* were upregulated (Fig. 7c). Matching the downstream targets of these transcription factors with the list of modulated genes revealed that *Nr2f2* and *Cux1* play critical roles in activating many downregulated genes and repressing several upregulated genes (Fig. 7d). These transcription factors are direct targets of *Nr4a2*, which promotes their transcription (Fig. 7d). Together, these results indicate that *Nr4a2* exerts a dose-dependent regulation over CLA and shell identities by modulating the expression levels of *Nr2f2* and *Cux1*.

## Discussion

In this study, we first characterized the transcriptomic identity and spatial organization of cell types and subtypes in the mouse claustro-insular region at high-resolution, then created a versatile map of this region using cell type-specific marker genes, and ultimately demonstrated that *Nr4a2*, a gene encoding a nuclear transcription factor, plays a crucial role in the development of this brain area.

By combining single-cell RNA sequencing with MERFISH spatial transcriptomics, we molecularly distinguished and spatially localized several cortical and subcortical neuronal cell types in the mouse claustro-insular region, including L2/3, L5b, L6b and several L6a and striatal populations. Our analysis revealed two subtypes of CLA projection neurons, CLA 1 and CLA 2, both defined by their expression of canonical CLA marker genes such as *Nr4a2*, *Gnb4*, *Smim32,* and *Slc17a6*. While the CLA 2 population can be distinguished by the expression of immediate early genes, both CLA subtypes are genetically and spatially indistinguishable, supporting previous findings of a single transcriptomically defined CLA population. Notably, the CLA 2 subtype comprises only 8% of all CLA neurons, suggesting it may represent recently activated CLA 1 neurons whose transcriptomic profile is transiently modulated, a sparse population missed in previous studies.

Do the CLA 1 and CLA 2 subtypes exclusively correspond to CLA projection neurons? Previous transcriptomic studies primarily focused on CLA neurons, often overlooking EP neurons and the less characterized dorsal and caudal CLA-like neurons located within L6a (e.g., bregma - 1.055)^13–16,18–24,58^. Our MERFISH and smFISH data indicate that EP and CLA-like L6a neurons are molecularly indistinguishable from CLA projection neurons, suggesting that these neurons represent the same cell type but are differentiated by their localization, likely along with their projection targets and functions within the brain.

In addition to CLA neurons, we identified a second neuronal population with a transcriptomic identity that lies intermediate between claustral and cortical neurons. We named these neurons “shell projection neurons”, as they likely correspond to the “claustrum shell” neurons described by Erwin et al.^19^. However, their transcriptomic profiles and spatial organization significantly differ from those of the CLA neurons, leading us to classify them as a separate population. This population is subdivided into three subtypes, each characterized by specific expression of *Syt17* and low levels of *Nr4a2*, and exhibiting distinct and stereotypical spatial organizations within the mouse brain.

Several anatomical definitions of the CLA boundaries have been proposed over the years^59–64,32,65^, including a subdivision of the CLA into a core and a shell^19,29^. Our findings demonstrate that CLA, shell, and cortical neurons exhibit a significant overlap in their spatial localizations, indicating that sharp delineations around claustro-insular structures are inadequate. Additionally, studies by other groups using retrograde tracing have shown a partial overlap in the projection sites of CLA and shell neurons, complicating the distinction between these populations based solely on their projection targets^29,19^. Given the complexity of this brain region, we propose a new map for the claustro-insular area, grounded in its cell types. Using a data-driven approach, involving smFISH labeling of coronal sections, we have constructed a versatile map that accounts for this overlap. Whether or not shell neurons are considered part of the CLA complex, in this map, these cells are appropriately annotated. We demonstrate the utility of this map by projecting cells from the MERFISH dataset onto it. Moreover, this map can also be overlaid over any labeled coronal section of the mouse claustro-insular region, using landmarks such as the tip of the piriform cortex and the lining of the external capsule.

Do CLA and shell neurons share the same developmental origin? Both populations express *Nr4a2*, albeit at different levels, and given the sustained and early expression of this transcription factor in developing CLA neurons^35,37^, we aimed to investigate *Nr4a2’s* role in the development of the claustro-insular region. Using a haploinsufficiency model, we surprisingly found that only neurons with high *Nr4a2* expression (i.e. CLA neurons) exhibited transcriptomic alterations and reduced abundance, while shell neurons, characterized by lower *Nr4a2* expression levels, were minimally affected, if at all. In CLA neurons, many of the modulated genes were associated to cell type identity, with numerous canonical CLA markers showing downregulation. Another research group, whose data was published while preparing this manuscript, also demonstrated a downregulation of CLA identity genes (though at the population level) in *Nr4a2* deficient mice^66^. Our results expand on these observations, at the single-cell level. Building on our high-resolution atlas, we also show that upregulated genes in *Nr4a2^del/wt^*CLA neurons were indicative of shell marker genes. These findings, along with predictions of a linear classifier, suggest that both shell and CLA neuronal populations are developmentally related, with *Nr4a2* transcription levels influencing, if not determining, their cell type identity in a dose-dependent manner. Unexpectedly, our results also indicate that NR4A2 does not directly modulate the expression of these identity genes, but instead orchestrates a cascade of other transcription factors that influence a broad array of target genes. Future studies should investigate whether CLA and shell neurons arise from a common pool of precursor cells and explore the roles of NR2F2 and CUX1 in defining their identities.

Among genes modulated in *Nr4a2* haploinsufficient mice, we identified those coding for channels or channel subunits, such as *Scn1b* and *Ryr2*. *Scn1b* showed higher transcript levels in wild-type CLA neurons, whereas *Ryr2* was more prevalent in shell neurons, suggesting differing firing properties between these neurons. Several studies have reported the existence of distinct groups of CLA neurons characterized by varied electrophysiological profiles^25–28^. Moreover, the projection sites and functional effects of CLA neurons in the cortex remain controversial, with studies reporting disynaptic inhibition of deep-layer cortical neurons and others excitation of upper-layer neurons^33,58^. Whether these functional subtypes of neurons correspond to CLA and shell neurons remains to be determined, but these results constitute an important first step in that direction.

What are the behavioral implications of *Nr4a2* haploinsufficiency-mediated alterations in the claustro-insular region? Several studies have reported that alterations in the excitation of CLA projection neurons lead to behavioral deficits in mice, particularly affecting higher-order cognitive functions^67–69,58,70,71^. We demonstrate that the overexpression of *Ryr2* in *Nr4a2* heterozygous mice is responsible for the hypoexcitability of CLA projection neurons. Given that *Nr4a2* coding variants are associated with neurodevelopmental disorders such as schizophrenia, and considering the electrophysiological alterations we observed, we hypothesize that these functional impairments may contribute to specific higher-order cognitive deficits, rather than to generic deficits linked to alterations in downstream or upstream brain targets of the CLA.

In summary, we developed a high-resolution map of the claustro-insular region, providing a nuanced understanding of the CLA, shell, and cortical populations. Our findings highlight that using transgenic mouse lines is the most effective way to achieve specificity when targeting these cell populations. This work lays the foundation for future studies aimed at exploring the developmental origins of CLA and shell neurons and their roles in higher-order cognitive functions, particularly in the context of neurodevelopmental disorders.

## Supplementary Figures

**Supplementary Fig. 1.**
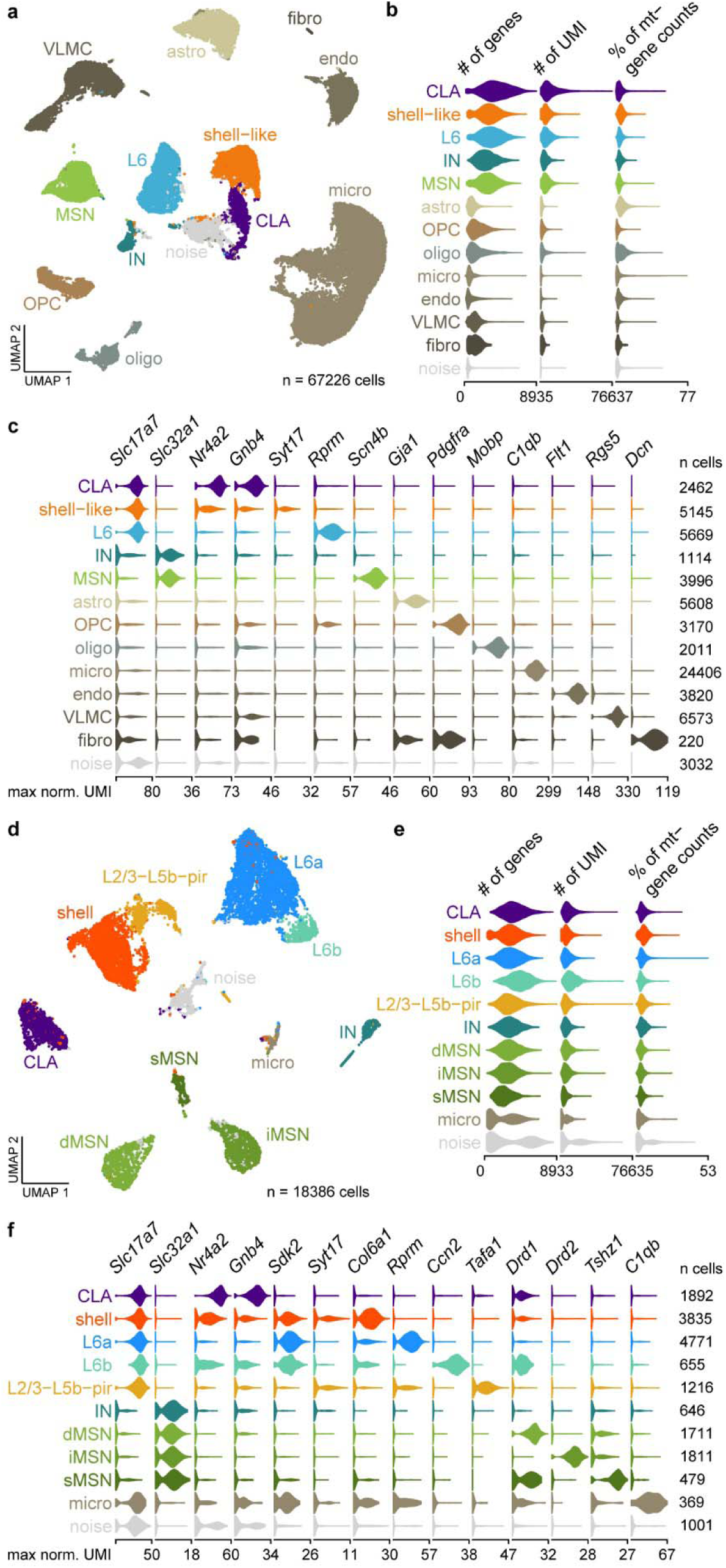
Cell types in the claustro-insular region detected by scRNAseq. **a** Visualization of neuronal and non-neuronal cell types from the scRNAseq dataset before cell filtering on a UMAP plot. **b** Violin plots showing the distribution of the number of genes, number of UMI counts, and percentage of mitochondrial gene counts in neuronal and non-neuronal cell types from the scRNAseq dataset before cell filtering (raw data are shown in linear scale). **c** Violin plots showing neuronal and non-neuronal cell type-specific distribution of expression of marker genes in the scRNAseq dataset before cell filtering (normalized data are shown in log_10_ scale). **d** Visualization of neuronal cell types from the scRNAseq dataset before cell filtering on a UMAP plot. **e** Violin plots showing the distribution of the number of genes, number of UMI counts, and percentage of mitochondrial gene counts in neuronal cell types from the scRNAseq dataset before cell filtering (raw data are shown in linear scale). **f** Violin plots showing neuronal cell type-specific distribution of expression of marker genes marker genes expression in the scRNAseq dataset before cell filtering (normalized data are shown in log_10_ scale).

**Supplementary Fig. 2.**
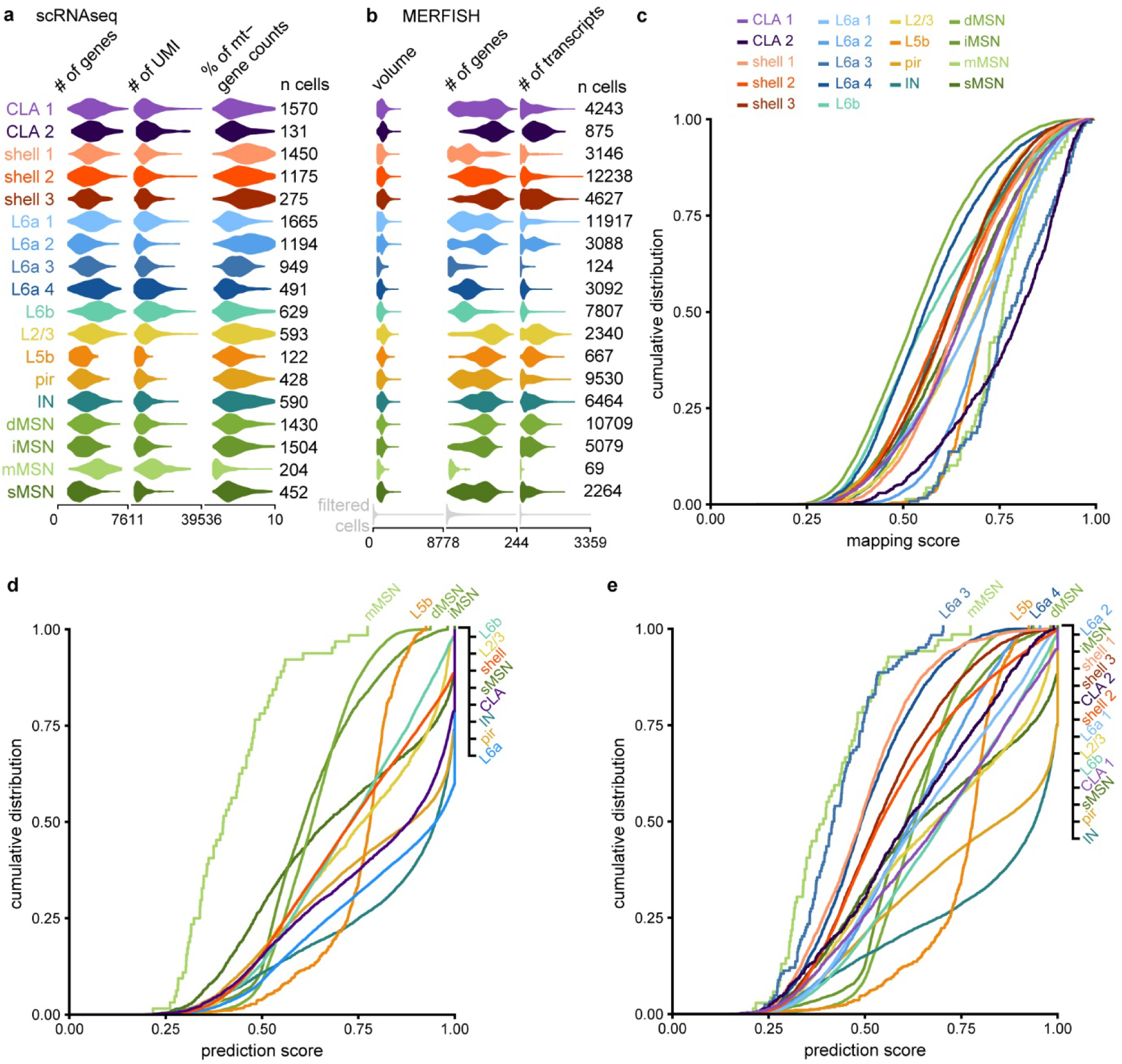
Quality control of scRNAseq and MERFISH data processing. **a** Violin plots showing the distribution of the number of genes, number of UMI counts, and percentage of mitochondrial gene counts in neuronal cell types from the scRNAseq dataset after cell filtering (raw data are shown in linear scale). **b** Violin plots showing the distribution of the cell volume, number of genes, and number of transcripts in neuronal cell types from the MERFISH dataset after reference-based mapping onto the scRNAseq dataset (raw data are shown in linear scale). **c** Empirical cumulative distribution of mapping scores for neuronal cell types and subtypes from the MERFISH dataset after reference-based mapping onto the scRNAseq dataset. Higher scores indicate greater confidence in the representation of MERFISH cells by the scRNAseq reference dataset. **d,e** Empirical cumulative distribution of prediction scores for neuronal cell types (**d**) and subtypes (**e**) from the MERFISH dataset after reference-based mapping onto the scRNAseq dataset. Higher scores indicate greater confidence in the attribution of cell type and subtype identities in the MERFISH dataset.

**Supplementary Fig. 3.**
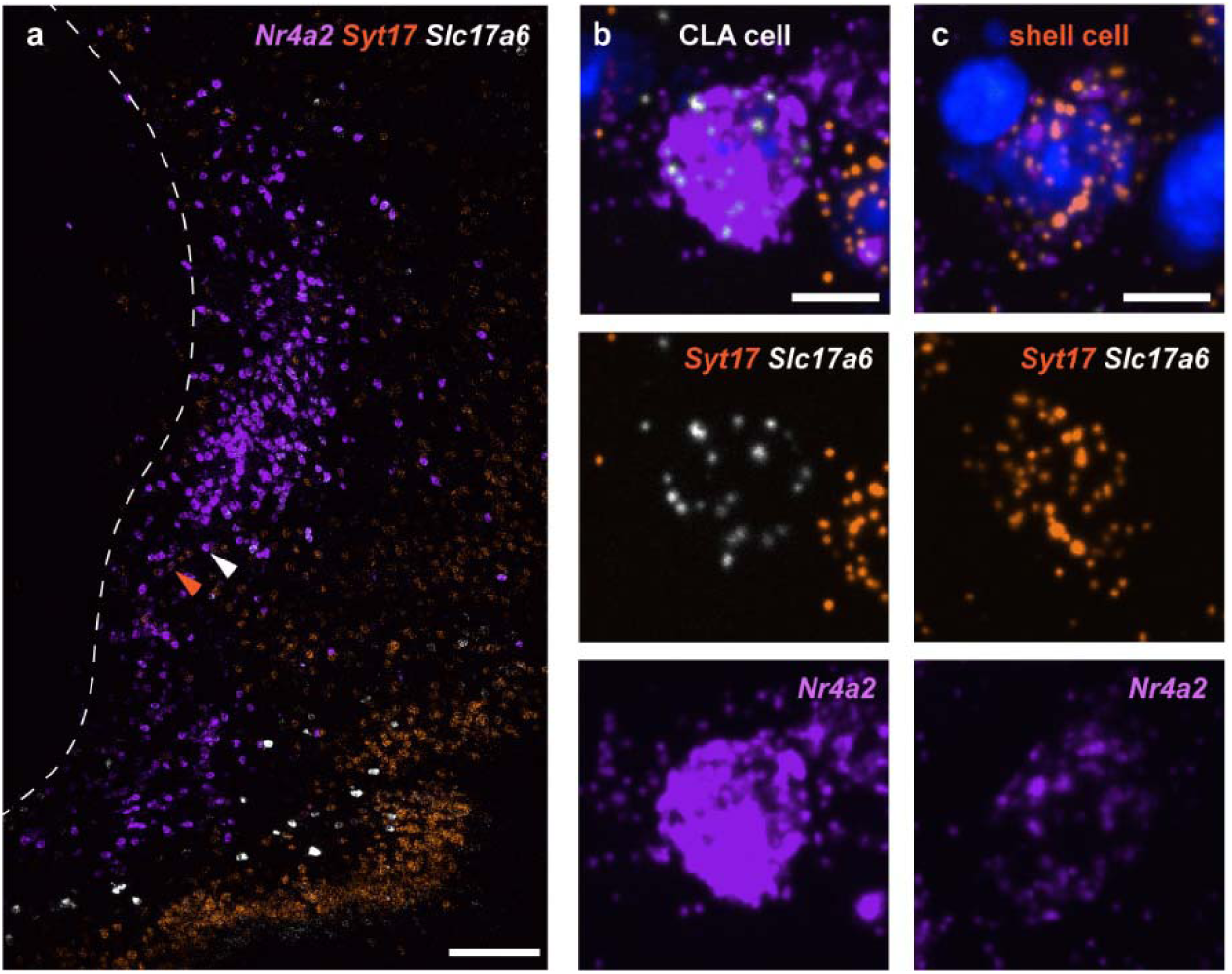
Illustration of CLA and shell cells characterized by high and low expression of *Nr4a2*. **a** Representative image of a coronal section (bregma +1.045) of the claustro-insular region labeled by smFISH with probes for *Nr4a2* (purple), *Syt17* (orange) and *Slc17a6* (white). Arrowheads point towards a CLA cell (white) and a shell cell (orange) shown in (**b,c**). The dotted line indicates the border of the external capsule. Scale bar: 200 μm. **b,c** Higher magnification of cells characterized by high (**b**) and low (**c**) expression of *Nr4a2*. DAPI staining of the nuclei (blue). Scale bar: 10 µm.

**Supplementary Fig. 4.**
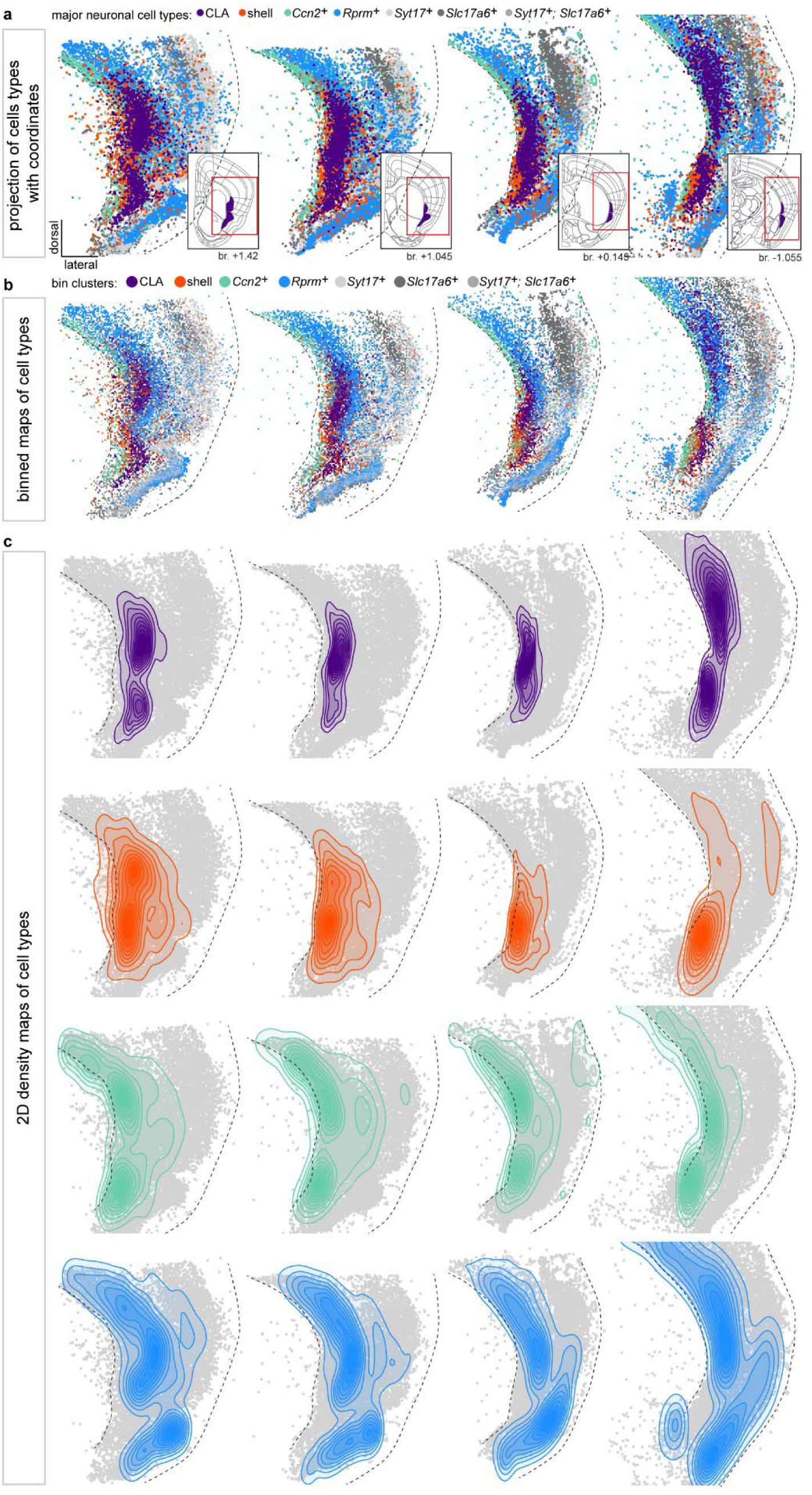
Map of the claustro-insular region. **a** Spatial organization of neuronal cell populations identified in the smFISH dataset, at different anteroposterior levels. Multiple images are aligned and cells are projected on the same framework. Inset shows the representation of the CLA and EP in the Allen Mouse Brain Atlas (mouse.brain-map.org), at the corresponding antero-posterior position. Red squares highlight the region imaged after smFISH labelling for map generation. **b** Proportional visualization of the claustro-insular region, binned into 20 μm bins, at different anteroposterior levels. Bin colors are weighted by the relative proportion of neuronal cell types within that bin. **c** 2D density maps showing the spatial distribution of neuronal cell populations within the claustro-insular region, at different anteroposterior levels. The total area comprises 100% of the 2D kernel density estimation of the distribution of all cells pertaining to the corresponding population.

**Supplementary Fig. 5.**
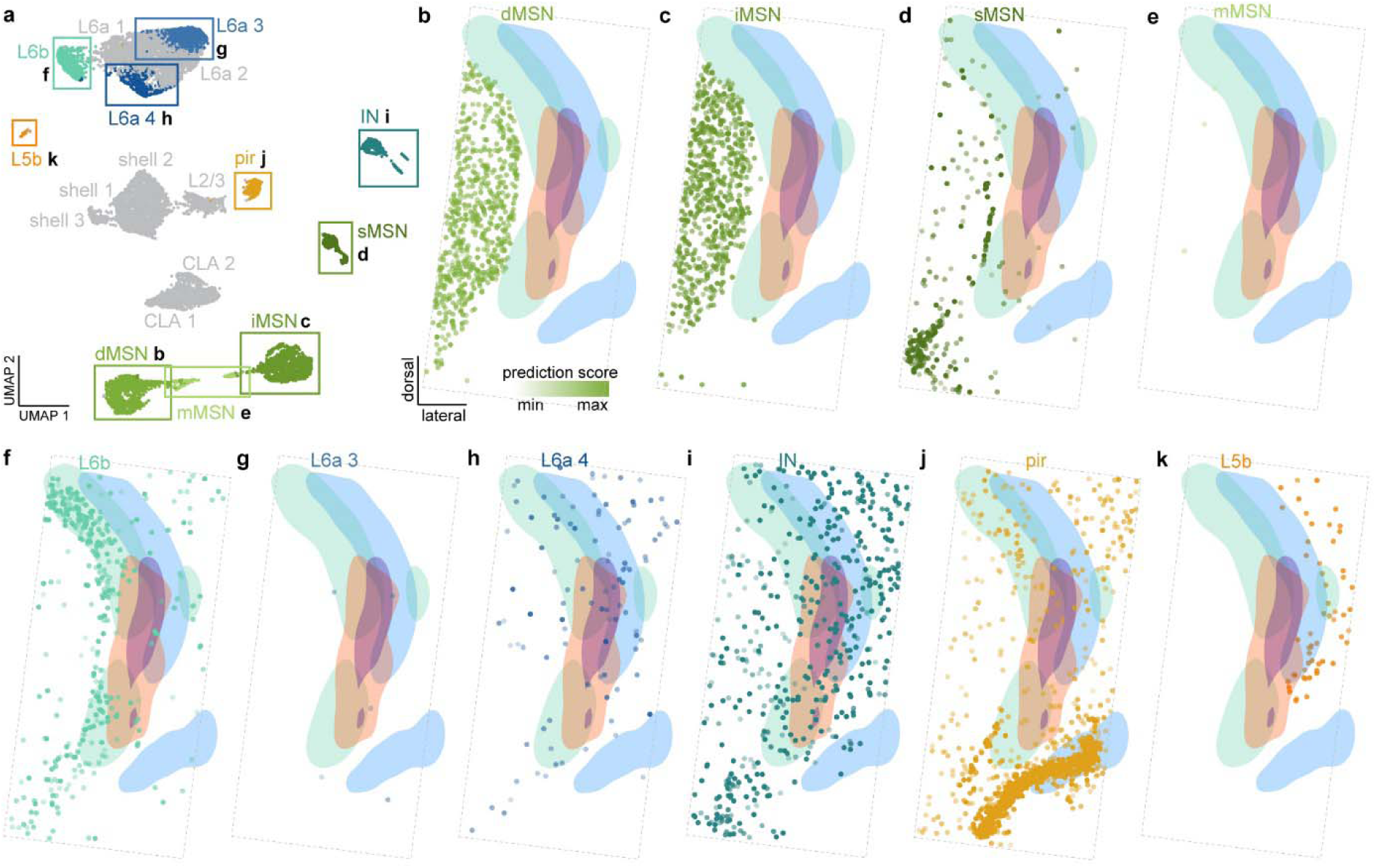
Spatial organization of cell types and subtypes in the claustro-insular region. **a** UMAP plot highlighting the cell types or subtypes whose localization was determined by MERFISH spatial transcriptomics. **b-k** Spatial organization of different neuronal types and subtypes projected on the map of the claustro-insular region at bregma +1.045. Prediction scores highlight the confidence of cell type attribution in the MERFISH data after reference-based mapping onto the scRNAseq dataset. Dotted lines highlight the border of the ROI used for cell selection. Cells from one representative section are shown for each cell type and subtype.

**Supplementary Fig. 6.**
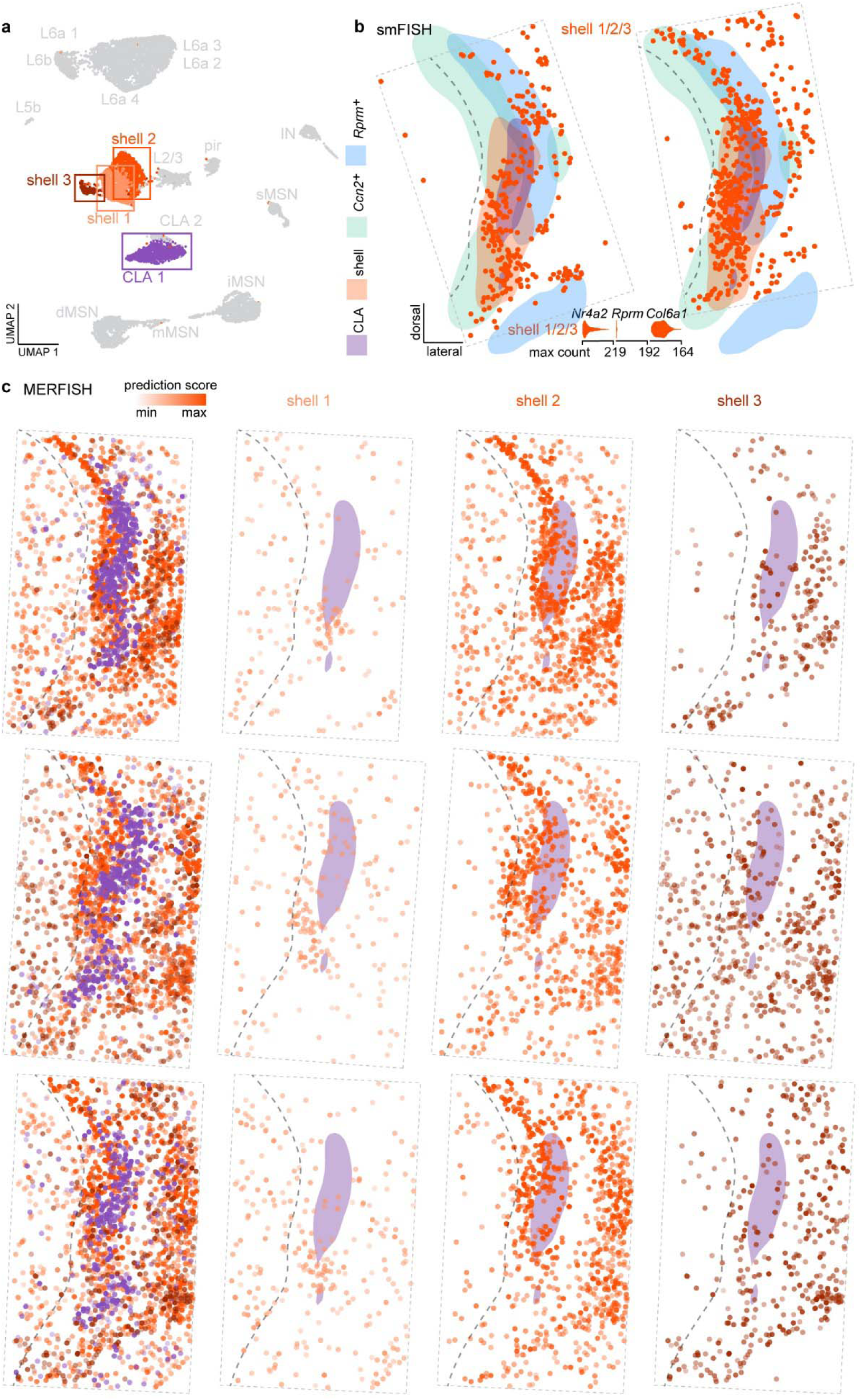
Evaluation of inter-individual variation in shell subtype spatial organization. **a** UMAP plot highlighting the cell subtypes whose localization was determined by smFISH and MERFISH spatial transcriptomics. **b** Spatial organization of shell cells on two different sections labeled by smFISH. Violin plots show the distribution of expression of marker genes used for cell type identification in the smFISH data (raw data are shown in linear scale). Shell cells are characterized by low expression of *Nr4a2*, no expression of *Rprm*, and high expression of *Col6a1*. **c** Spatial organization of shell 1, shell 2, and shell 3 subtypes from three distinct mice from the MERFISH dataset. Cells were projected on the map of the claustro-insular region at bregma +1.045. The purple shaded area corresponds to the CLA reference on the map and is used to highlight the position of shell subtypes relative to the CLA. The dotted line highlights the border of the external capsule on the reference map.

**Supplementary Fig. 7.**
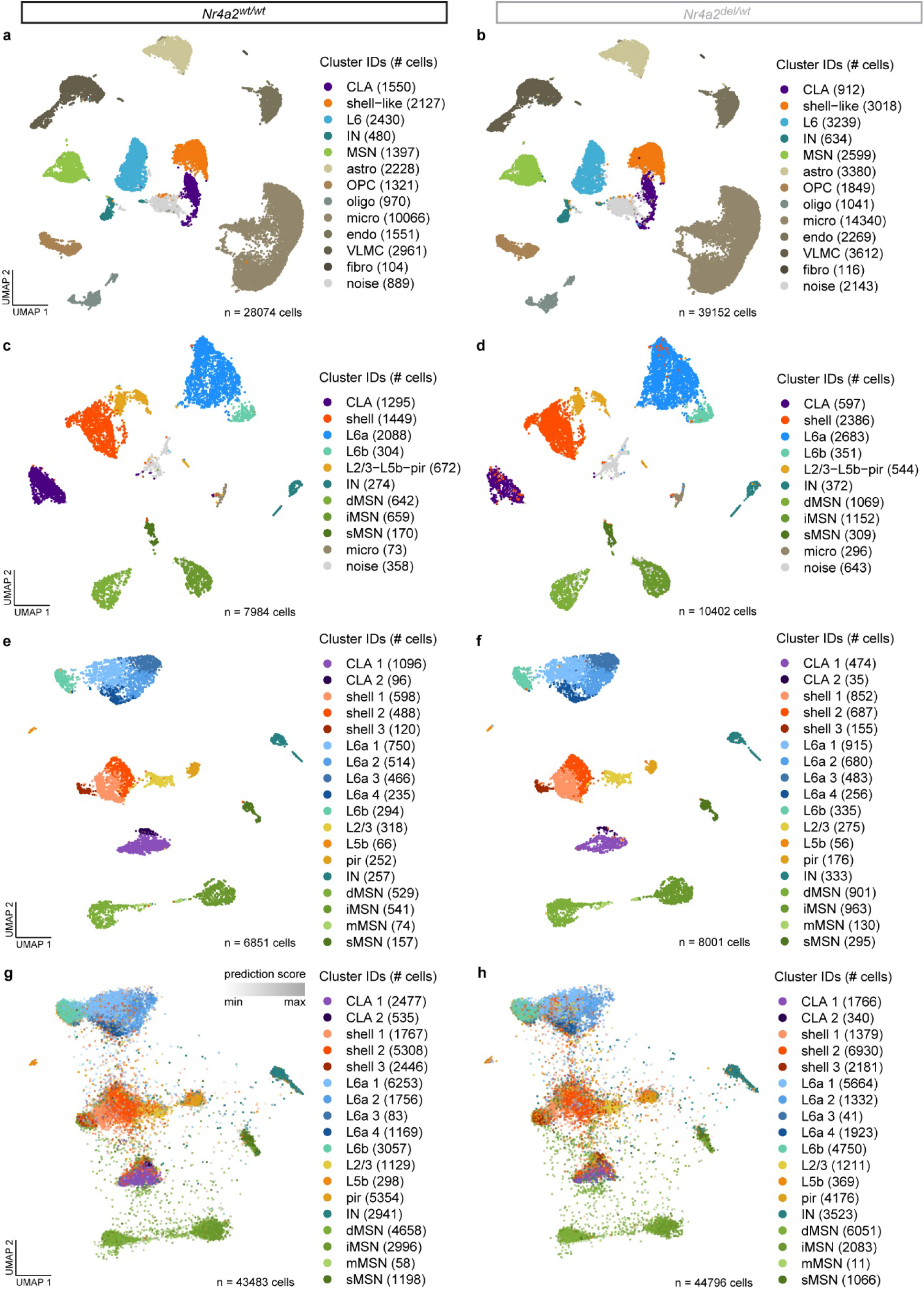
scRNAseq and MERFISH UMAP plots of cells from *Nr4a2* wild-type and haploinsufficient mice. **a,b** Visualization of neuronal and non-neuronal cell types in *Nr4a2^wt/wt^*(**a**) and *Nr4a2^del/wt^*(**b**) mice from the scRNAseq dataset before cell filtering on a UMAP plot. **c,d** Visualization of neuronal cell types in *Nr4a2^wt/wt^* (**c**) and *Nr4a2^del/wt^* (**d**) mice from the scRNAseq dataset before cell filtering on a UMAP plot. **e,f** Visualization of neuronal cell types in *Nr4a2^wt/wt^* (**e**) and *Nr4a2^del/wt^* (**f**) mice from the scRNAseq dataset after cell filtering on a UMAP plot. **g,h** Visualization of neuronal cell types in *Nr4a2^wt/wt^* (**g**) and *Nr4a2^del/wt^* (**h**) mice from the MERFISH dataset after reference-based mapping onto the scRNAseq dataset on a UMAP plot.

**Supplementary Fig. 8.**
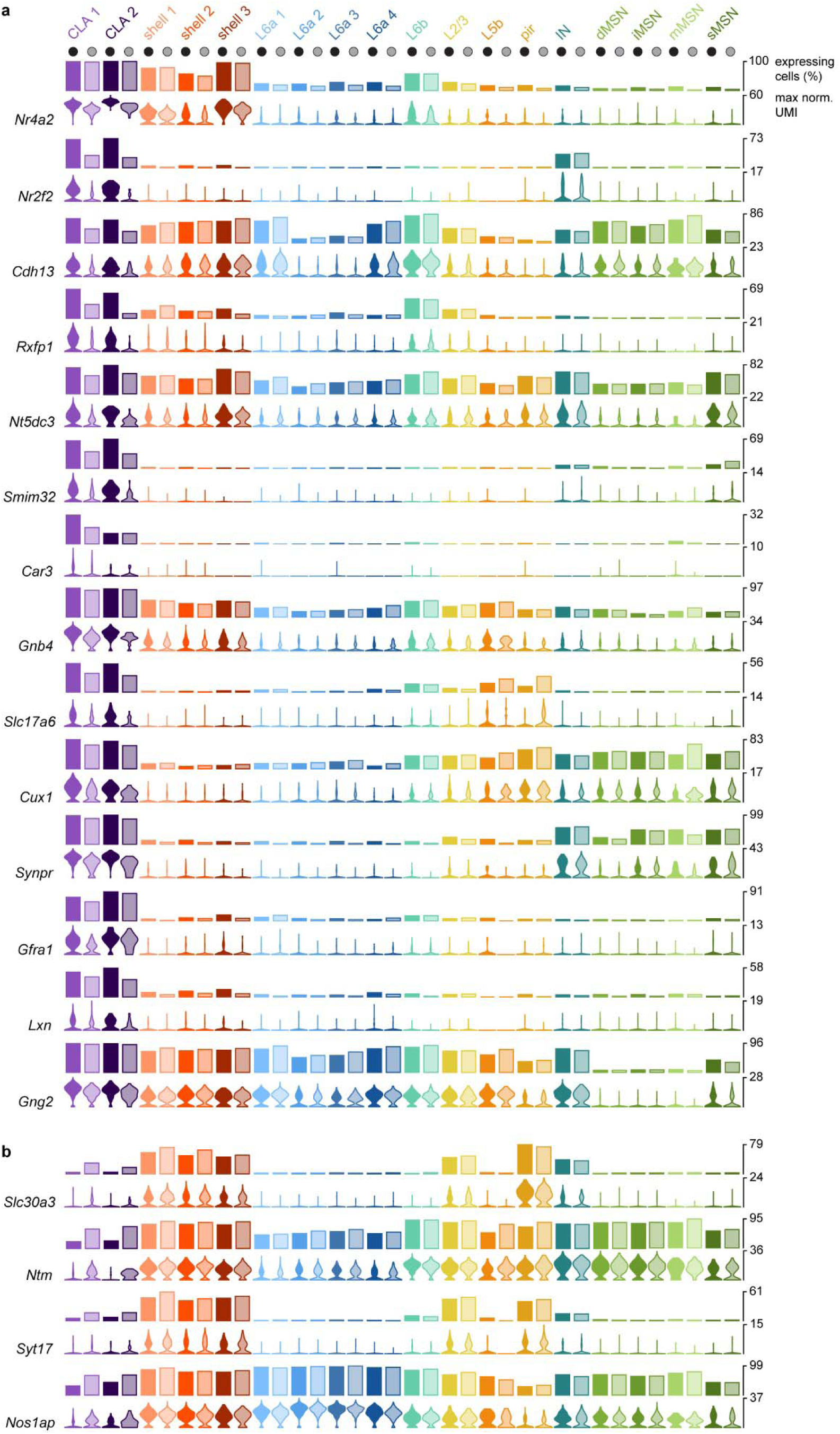
Modulation of claustro-insular marker genes in *Nr4a2* haploinsufficient mice. **a,b** A subset of downregulated (**a**) and upregulated (**b**) genes in *Nr4a2^del/wt^* CLA neurons. Downregulated genes in the plot are markers of CLA neurons, while upregulated genes are markers of shell neurons. *Top*: Bar plots representing the percentage of cells in each cell type and subtype from *Nr4a2^wt/wt^*and *Nr4a2^del/wt^* mice expressing CLA (**a**) and shell (**b**) marker genes at a minimum of 1 raw UMI count in the scRNAseq dataset. *Bottom*: Violin plots showing the distribution of expression of CLA (**a**) and shell (**b**) marker genes across claustro-insular cell types and subtypes from *Nr4a2^wt/wt^* and *Nr4a2^del/wt^* mice in the scRNAseq dataset (normalized data are shown in log_10_ scale).

**Supplementary Fig. 9.**
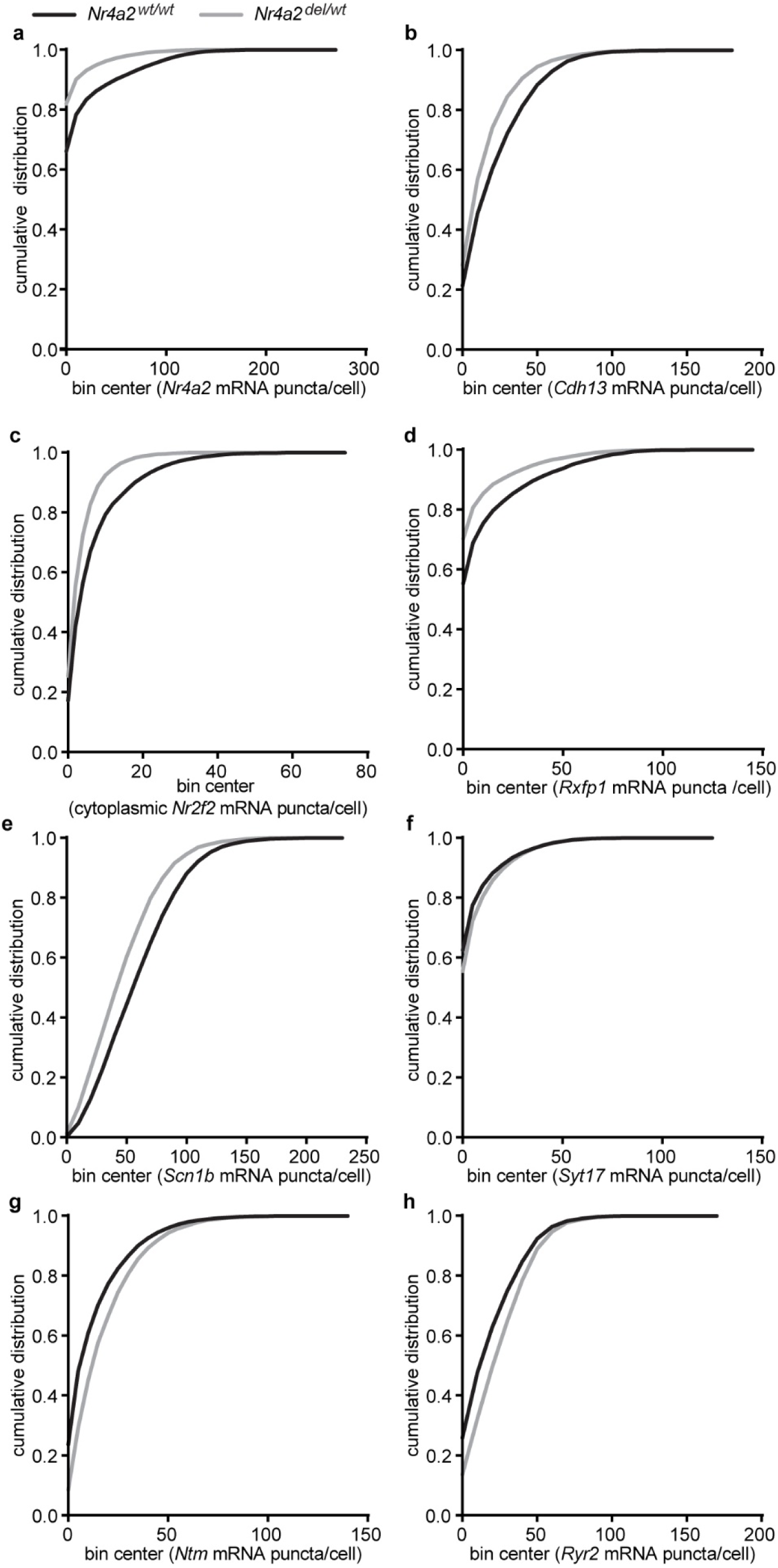
Transcriptomic alterations in *Nr4a2* haploinsufficient mice. **a** Empirical cumulative distribution graphs showing the modulation of *Nr4a2* and other genes (**b-h**) in all cells from the claustro-insular region of *Nr4a2^wt/wt^* (black) and *Nr4a2^del/wt^* (grey) mice. n = 4 *Nr4a2^wt/wt^* mice, 26 sections, 55’697 cells, n = 4 *Nr4a2^del/wt^* mice, 34 sections, 71’389 cells. **b** *Cdh13,* n = 3 *Nr4a2^wt/wt^* mice, 8 sections, 12’850 cells; n = 3 *Nr4a2^del/wt^* mice, 11 sections,17’503 cells, **c** *Nr2f2*, n = 2 *Nr4a2^wt/wt^* mice, 4 sections, 6’746 cells; n = 2 *Nr4a2^del/wt^* mice, 5 sections, 7’717 cells. **d** *Rxfp1*, n = 3 *Nr4a2^wt/wt^* mice, 8 sections, 12’850 cells; n = 3 *Nr4a2^del/wt^* mice, 11 sections, 17’503 cells. **e** *Scn1b*, n = 2 *Nr4a2^wt/wt^* mice, 7 sections, 11’310 cells; n = 2 *Nr4a2^del/wt^* mice, 5 sections, 7’702 cells. **f** *Syt17*, n = 3 *Nr4a2^wt/wt^* mice, 14 sections, 42’912 cells; n = 3 *Nr4a2^del/wt^* mice, 15 sections, 45’572 cells. **g** *Ntm*, n = 2 *Nr4a2^wt/wt^* mice, 6 sections, 18’367 cells; n = 2 *Nr4a2^del/wt^* mice, 8 sections, 24’504 cells. **h** *Ryr2*, n = 2 *Nr4a2^wt/wt^* mice, 8 sections, 24’545 cells; n = 2 *Nr4a2^del/wt^* mice, 7 sections, 21’068 cells.

**Supplementary Fig. 10.**
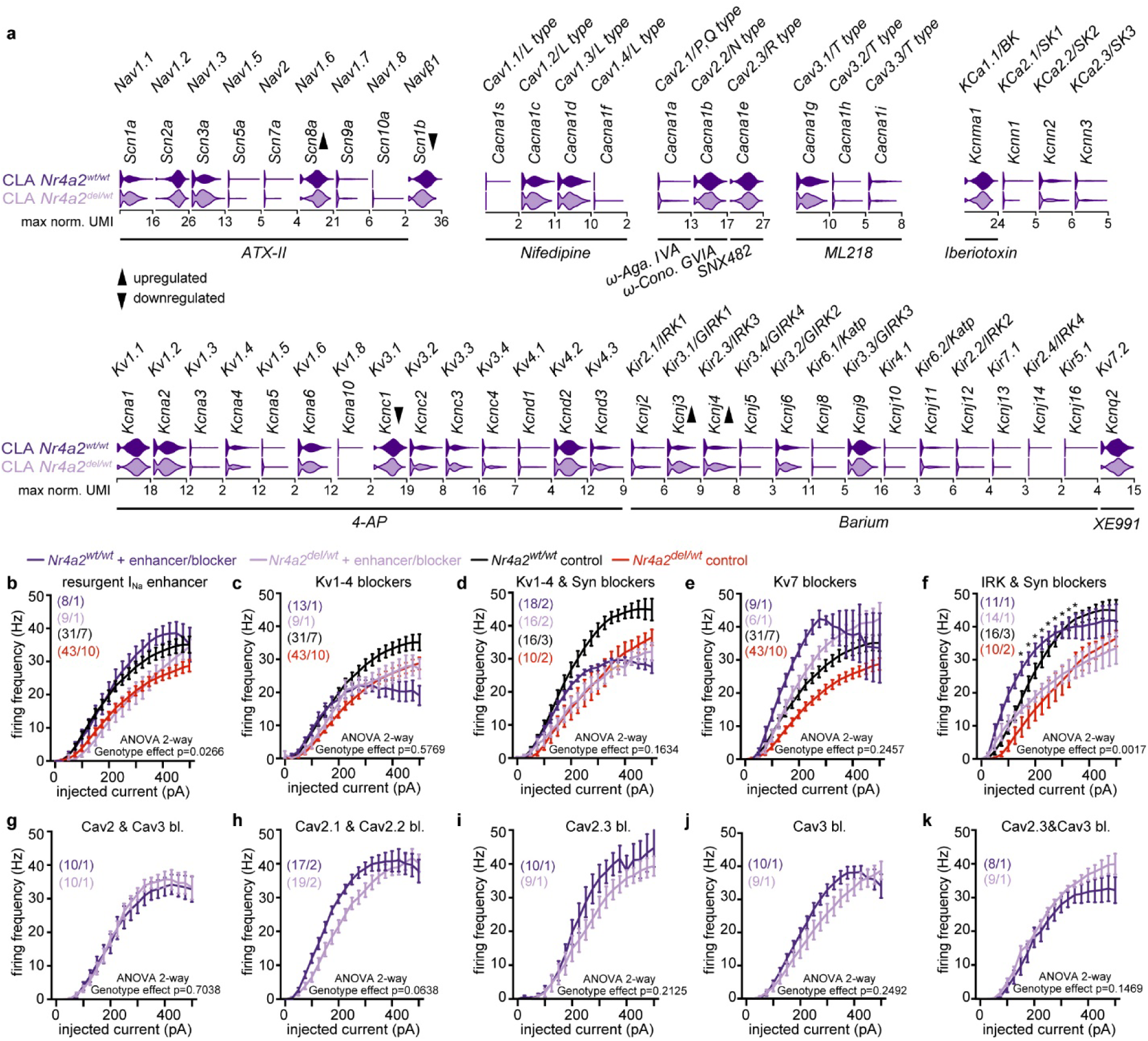
Expression profiles of genes coding for various ion channels and the effects of pharmacological agents on CLA neuron firing activity. **a** Expression level of genes encoding for voltage-gated sodium channels, voltage-gated calcium channels, and various potassium channel subtypes in CLA neurons of *Nr4a2^wt/wt^* (*top*) and *Nr4a2^del/wt^* (*bottom*) mice. The gene names, corresponding channel proteins, and relevant blocking agents are indicated. Genes encoding channels that are significantly downregulated and upregulated in CLA neurons of *Nr4a2^del/wt^* mice, compared to their wild-type littermates, are highlighted with downward-pointing and upward-pointing arrows, respectively. **b-k** Input/Output functions of CLA neurons recorded under various pharmacological conditions: an enhancer of resurgent sodium current ATX-II (**b**); Kv1-4 blocker 4-aminopyridine (1 mM, **c**); 4-aminopyridine (1 mM) and synaptic blockers (**d**); Kv7 blocker XE991 (20 µM, **e**); the inward rectifier blocker Barium and synaptic blockers (**f**); Cav2 and Cav3 blockers (ω-Agatoxin IVA (100 nM), ω-Conotoxin GVIA (3 µM), SNX-482 (400 nM), ML218 (2 µM)) (**g**); Cav 2.1 and Cav2.2 blockers (ω-Agatoxin IVA (100 nM), ω-Conotoxin GVIA (3 µM)) (**h**); Cav2.3 blocker (SNX-482 (400 nM)) (**i**); Cav3 blocker (ML218 (2 µM)) (**j**); Cav2.3 and Cav3 blockers (**k**). To compare the effects of certain blockers with control conditions, input/output functions recorded in ACSF (**b,c,e**) or in the presence of synaptic blockers (**d,f**) for wild-type (black) and haploinsufficient (red) mice were included in some summary graphs. Data are shown as mean ± SEM. 2-way repeated measures ANOVA are followed by post-hoc testing using Fisher’s LSD test with a 5% false discovery rate; *p*-values were adjusted for multiple comparisons using the Benjamini-Hochberg false discovery rate method. See Supplementary Tables, Statistics table for more details.

**Supplementary Fig. 11.**
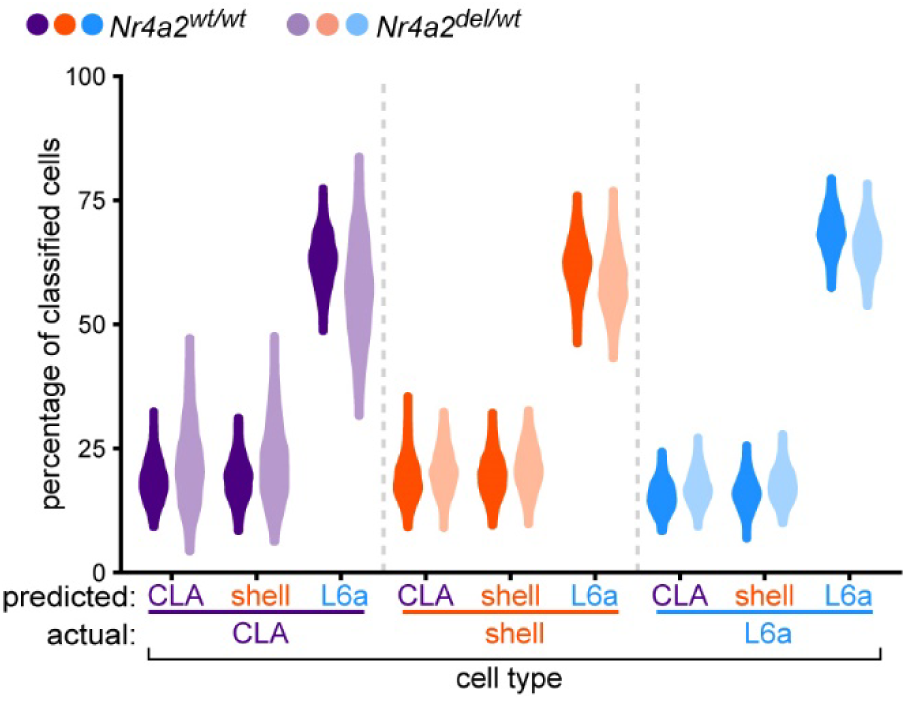
Cell type classification performance after shuffling of identities. Cell type classification performance of CLA, shell and L6a neurons of *Nr4a2^wt/wt^* and *Nr4a2^del/wt^* mice from the scRNAseq dataset using a linear support vector classification algorithm after shuffling of cell type identities. Violin plots show the distribution of classification outcomes across 1’000 iterations.

## Methods

### Animals

All experiments were conducted in accordance with the veterinary guidelines and regulations of the University and of the state of Geneva (Direction de l’expérimentation animale de l’UNIGE). C57BL/6J male mice were purchased at 5–7 weeks of age from Charles River Laboratories. In Nr4a2-SA-IRES-Dre mice, the complete coding sequence of *Nr4a2* was replaced by a splice acceptor–ires-Dre sequence (*Nr4a2^tm^*^1^*^(dreo)Hze^*, MGI: 6790280)^72^. Heterozygous mice were obtained by crossing a heterozygous parent (male or female) with a wild-type parent of the same genetic background (male or female). Mice heterozygous for this mutation are referred to as *Nr4a2^del/wt^*mice, and their wild-type littermates as *Nr4a2^wt/wt^* mice. For electrophysiogical recordings of genetically identified CLA neurons, *Slc17a6^tm2(cre)Lowl^*/J mice^73^ (also known as *Vglut2*-IRES-cre; the Jackson laboratory, strain number 016963) were crossed to *Nr4a2*^tm1(dreo)Hze^ mice. In *Vglut2*-IRES-cre mice, the cre coding sequence does not disrupt the *Slc17a6* gene coding sequence since it was placed after the stop codon. Upon arrival, mice were group-housed in standard type II cages with *ad libitum* access to food and water. They were held on a dark/light cycle of 12/12□h, with controlled temperature (21-22□°C) and humidity (45-55%).

### scRNA-sequencing

#### Tissue collection and dissociation

*Nr4a2^wt/wt^* and *Nr4a2^del/wt^* male (n = 4 8-9 weeks and 3 15 weeks old) and female (n = 3 8-9 weeks and 3 15 weeks old) littermate mice were used for scRNAseq experiments. Tissue collection and dissociation were performed following the methods described by Fodoulian et al. (2020)^58^. Mice were anesthetized with isoflurane and sacrificed by decapitation. Brains were immediately extracted and placed in ice-cold oxygenated artificial cerebro-spinal fluid (ACSF) containing (in mM): 124 NaCl, 3 KCl, 2 CaCl_2_, 1.3 MgSO_4_, 26 NaHCO_3_, 1.25 NaH_2_PO_4_, 10 D-glucose with an osmolarity of 300 mOsm and pH at 7.4 when oxygenated with 95% O_2_, 5% CO_2_. Coronal sections of the brain spanning the antero-posterior localization of the CLA were cut at a thickness of 300 μm using a vibrating-blade microtome (Leica VT1000S; Leica Biosystems). Immediately after sectioning, the CLA and adjacent brain regions were microdissected under a stereomicroscope. Tissue dissociations were performed using the Papain Dissociation System (ref. LK003150; Worthington® Biochemical Corporation) according to the manufacturer’s protocol with slight modifications, using EBSS and medium with serum solutions adapted from Saxena et al., (2020)^74^. After microdissection, extracted tissues were kept in ice-cold oxygenated EBBS#1^74^. Tissues from each mouse were processed individually, with no pooling across different animals. Papain and DNase I vials provided with the kit were reconstituted using the oxygenated EBSS#1 solution. Tissues from each mouse were transferred to a 50 ml Falcon tube containing 5 ml of the papain/DNase I mix, and were incubated at 37°C for 20 minutes in a rotating laboratory oven (ProBlot^TM^ Hybridization Oven; Labnet International) at a speed of 8-10 rpm. Tissues were then gently triturated using a P1000 pipette and left to settle for 1 minute. Cell suspensions were then passed through a 70 μm nylon cell strainer (ref. 352350, Falcon) to remove undigested tissue fragments, transferred to 15 ml Falcon tubes, and centrifuged (Centrifuge 5430R; Eppendorf) at 300 g for 5 min. Supernatants were discarded, and cell pellets were resuspended with 3ml of EBSS#2^74^ to stop digestion. In order to remove myelin debris, two discontinuous density gradients were produced by carefully adding the 3 ml of cell suspensions on top of 5 ml of an ovomucoid-albumin protease inhibitor solution reconstituted following the manufacturer’s instructions. Tubes were centrifuged at 70 g for 6 min at room temperature, the supernatants discarded and the cell pellets re-suspended in 1 ml medium solution with serum. Cell suspensions were then passed through another 70 μm nylon cell strainer.

### Fluorescence-activated cell sorting (FACS)

To ensure the exclusive isolation of live nucleated cells, cell suspensions were incubated with 2 μg/ml of Hoechst 33342 (a UV fluorescent adenine-thymine binding dye; ref. H1399, Life Technologies^TM^) at 37°C for 15 min. In order to exclude dead cells, 1 μM of DRAQ7^TM^ (a far-red fluorescent DNA intercalating dye; ref. DR71000, BioStatus) was added to the cell suspensions before FACS sorting. Hoechst^+^/DRAQ7^-^ cells were then sorted in an empty Eppendorf tube according to their forward scatter (FSC) and side scatter (SSC) properties using a Beckman Coulter MoFlo Astrios cell sorter with a 100 μm nozzle at a pressure of 25 psi. Doublets were excluded after gating on FSC-A/FSC-H, followed by SSC-H/SSC-W. Between 11’000-25’000 Hoechst^+^/DRAQ7^-^ cells were collected from each sample.

### Single-cell capture, cDNA library preparation, and RNA sequencing

Single-cell capture, reverse transcription, cDNA amplification, and library preparation were performed using the 10x Genomics Chromium Controller instrument and the Single Cell 3′ v3.1 reagent kit with dual indexes, following the manufacturer’s protocol (CG000315 Rev B User Guide). Cells were loaded on the 10x Genomics Chromium Controller instrument at a concentration of 400 cells/μl, and the targeted cell recovery was adjusted for each sample based on the number of sorted cells. Eleven rounds of PCR amplification were conducted after reverse transcription, and 13 to 14 rounds during library preparation. After the first round of PCR, cDNA quality and quantity were assessed using the Agilent 2100 Bioanalyzer with the Agilent Bioanalyzer High Sensitivity DNA Assay Kit (ref. 5067-4626; Agilent Technologies). After library preparation, cDNA library molarity and quality were assessed using the Qubit 4.0 with the Qubit™ 1X dsDNA HS Assay Kit (ref. Q33231; Thermo Fisher Scientific) and the TapeStation with the Agilent D1000 ScreenTape (ref. 5067-5582; Agilent Technologies). Libraries from mice of the same age were then pooled and each loaded at 2 nM on 4 lanes for clustering on a paired-end Illumina Flow Cell using the HiSeq 4000 PE Cluster Kit (ref. PE-410-1001; Illumina) and sequenced on an Illumina HiSeq 4000 sequencer using the HiSeq 4000 SBS Kit (ref. FC-410-1001; Illumina) chemistry. Read 1 consisted of 28 bases, while read 2 was either 90 or 100 bases. An average of 5’171 cells were sequenced with 42’199 mean reads, 1’125 median genes, and 1’999 median UMI counts per cell.

### scRNAseq data analysis

#### Demultiplexing

Demultiplexing of Illumina indices was performed on the raw Illumina data using bcl2fastq2 Conversion Software version 2.20.0 (Illumina), which generated a pair of fastq files per sample (i.e. reads 1 and 2).

#### Mapping

Digital gene expression matrices (i.e. matrices with genes in rows, cells in columns, and gene counts as matrix entries) were generated using Cell Ranger version 6.0.0^75^. Fastq files were mapped to the *Mus musculus* genome primary assembly reference 38 (GRCm38) using STAR^76^, implemented in Cell Ranger. A modified version of Ensembl release 102 of the *Mus musculus* GTF annotation was used, with the *Gm45623* annotation replaced by a custom annotation for *Smim32*, as described in Tuberosa et al., 2024^52^. Mapping, UMI counting, and barcode filtering were performed using the count function of Cell Ranger. The expected number of recovered cells (--expect-cells argument) was customized for each sample and set to the targeted cell recovery of the sample.

#### Data filtering

scRNAseq data analyses were performed using R version 4.3.1^77^. The filtered feature-barcode matrices from the Cell Ranger output served as the initial files for downstream analyses. Three rounds of clustering (following the steps detailed below) were applied to the data. The datasets at each round will be named “all cells” (Supplementary Fig. 1a), “neurons unfiltered” (Supplementary Fig. 1d), and “neurons filtered” (Fig. 1b) in the rest of the methods section. Cells were iteratively filtered to retain high-quality cells. After the initial clustering step of the “all cells” dataset, only neuronal clusters were retained for further analysis. These cells (forming the “neurons unfiltered” dataset) were re-clustered to remove clusters formed by low-quality cells or doublets that could not be detected in the full dataset. A final filtering step was applied to the cells, following the criteria detailed in Fodoulian et al. 2020. Briefly, we removed all cells 1) that were characterized by fewer than 1000 expressed genes, 2) for which mitochondrial counts exceeded 10% of their total counts, as high mitochondrial counts indicate suffering or dead cells, 3) for which the total number of detected genes, total number of UMIs, and percentage of mitochondrial counts were three median absolute deviations away from the median after log_10_ transformation, and 4) that showed unusually high or low numbers of genes given their total number of UMIs, after log_10_ transformation, by fitting a loess curve (*loess* function of the stats R package, with span = 0.5 and degree = 2), where the number of genes was taken as the response variable and the number of UMIs as the predictor. Cells for which the model residual was not within three median absolute deviations of the median were filtered out. The final number of cells in each dataset was 67’226 for “all cells”, 18’386 for “neurons unfiltered”, and 14’852 for “neurons filtered”. After cell filtering, we also selected genes expressed in at least three cells separately in each sample, with a minimum expression threshold of 1 UMI count. This filtering resulted in a total of 27’453 genes for “all cells”, 24’549 for “neurons unfiltered”, and 23’916 for “neurons filtered”.

#### Selection of high dropout genes

In order to identify distinct cellular populations, we selected genes for downstream analyses based on their mean expression and dropout-rate relationship using the M3Drop R package version 3.10.6^78^. We fitted a depth-adjusted negative binomial (DANB) model on the raw counts of each sample using the *NBumiFitModel* function of M3Drop. This model accounts for differences in sequencing depth and UMI tagging efficiency between cells. Genes with significantly higher dropout rates than expected by the model were selected for dimensionality reduction and clustering using the *NBumiFeatureSelectionCombinedDrop* function of M3Drop (*p*-value < 0.05, adjusted for multiple comparisons using the Benjamini-Hochberg false discovery rate method^79^. This selection corresponded to a total of 3’860 high dropout genes for “all cells”, 2’316 for “neurons unfiltered”, and 1’780 for “neurons filtered”.

#### Data normalization and confounder regression

To account for technical variability (i.e. tissue dissociation effects and sequencing depth differences between cells) in downstream analyses, a variance stabilizing transformation was applied to the raw counts of each sample using the *SCTransform* function of the Seurat R package version 5.1.0^80–82^. This function implements the *vst* function of the sctransform R package (version 0.4.1 was used)^80–82^. All cells and genes were used for initial negative binomial regression parameter estimation, with the method set to *glmGamPoi_offset* (implemented using the glmGamPoi R package version 1.12.2)^82,83^. The Pearson residuals from the second regularized negative binomial model were computed for all genes and clipped to a range of 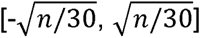. To correct for confounding variables, a final linear model was fitted to the Pearson residuals to regress out the number of UMIs per gene and the percentage of mitochondrial counts for each cell, and the residuals from this model (i.e. corrected Pearson residuals) were centered to the mean. Corrected counts and their log1p transformation were also computed. The process described here was applied either separately to each sample, with the resulting data used for dimensional reduction, clustering, or differential expression analysis - this latter after re-correcting UMI counts using the minimum of the median UMIs across samples via the *PrepSCTFindMarkers* function of Seurat - or applied to all cells together for the analysis of cell type identity shifts, gene regulatory network inference, and reference-based mapping on the “neurons filtered” dataset. For gene expression plotting, raw gene expression values were normalized by the total number of counts per cell and scaled to 10^4^ (i.e. corresponding to norm. UMI). Plotted gene expression values are actual normalized values, but the scales are in log_10_ after adding a pseudocount of 1.

#### Dimensionality reduction, dataset integration, and unsupervised graph-based clustering

Prior to cell clustering, principal component analysis (PCA) was performed on the corrected Pearson residuals of the high dropout genes (see Selection of high dropout genes) using the irlba R package version 2.3.5.1^84^, implemented in the *RunPCA* function of Seurat. Only the first 100 PCs were computed. The elbow in the decrease of the explained standard deviation of the successive PCs was detected using the *KneeLocator* function of the kneed Python package version 0.8.5^85^ with the following parameters: S = 1, curve = “convex”, direction = “decreasing”. The location of the elbow was then used as a threshold to retain the top relevant PCs for downstream processing of the datasets. This resulted in retaining 19 PCs for “all cells”, 20 for “neurons unfiltered”, and 19 for “neurons filtered” (explaining 82%, 78%, and 77% of the variance calculated with the top 100 PCs, respectively). These PC embeddings were subsequently used for dataset integration through the *RunHarmony* function of the harmony R package version 1.2.0^86^. The ridge regression penalty was automatically estimated (with lambda = NULL), the maximum number of Harmony rounds was set to 50, the maximum number of clustering rounds at each Harmony round was set to 200, and the convergence tolerance values for clustering and Harmony rounds were set to 1e-5. Harmony-corrected cell embeddings were then used for clustering and Uniform Manifold Approximation and Projection (UMAP)^87,88^ dimensional reduction. To construct a shared nearest-neighbor (SNN) graph, 15 approximate nearest neighbors were identified for each cell using the RcppAnnoy R package version 0.0.22, implemented in the *FindNeighbors* function of Seurat. A forest of 50 trees was built for the nearest neighbor search, using the Euclidean distance metric. The ranked nearest neighbors were then used to construct the SNN graph, and a Jaccard index cutoff of 1/15 was applied to prune weak connections from the graph. Cells were clustered using the leidenalg Python package version 0.10.2^89^, implemented in the *FindClusters* function of Seurat. The RBConfigurationVertexPartition singleton partition type was used, implementing Reichardt and Bornholdt’s Potts model with a null configuration^90,91^. The Leiden algorithm was run until no improvement in iteration was found (with n.iter = −1L). If singleton clusters were found, these were grouped into the same cluster. Clustering resolution selection varied by dataset. For the “all cells” and “neurons unfiltered” datasets, clustering resolution parameters ranged from 0 to 1 in increments of 0.1, and the lowest resolution that allowed segregation of large “noise” clusters was selected. For the “neurons filtered” dataset, the variation of information (VI)^92^ between successive resolutions (ranging from 0 to 1.5 in increments of 0.1) was computed and normalized by the absolute difference in the number of clusters, and the resolution with the highest normalized VI was chosen. VI was computed using the *compare* function of the igraph R package version 2.0.3. These steps resulted in clustering resolutions of 0.3 for “all cells”, 0.2 for “neurons unfiltered”, and 1.2 for “neurons filtered”. UMAP dimensional reduction was computed using the uwot R package version 0.1.16, implemented in the *RunUMAP* function of Seurat. Since 15 nearest neighbors were identified for each cell for SNN graph construction, 15 neighboring points were used during the local approximation of the manifold structure, and the cosine metric was used to measure distances. Note that UMAP was solely used to visualize the cluster assignment of cells on a 2-dimensional plot.

#### Differential expression analysis

To identify cell type and subtype marker genes, or genes modulated due to *Nr4a2* haploinsufficiency, the Wilcoxon rank sum test was performed on the corrected, log1p-transformed counts, implemented in the *FindMarkers* function of Seurat. When identifying cell type marker genes, differential expression analysis was computed between each cell type and all other cell types combined. For cell subtype marker genes, pairwise comparisons between related cell types were conducted. To identify modulated genes, comparisons within cell types or subtypes were performed, comparing *Nr4a2^wt/wt^* cells to *Nr4a2^del/wt^*cells. Genes expressed in at least 10% of the cells in either group, with a log fold-change of 0.25 and an adjusted *p*-value < 0.05 using the Bonferroni method, were considered significant. To test for robustness in the obtained results, a leave-one-out approach was implemented, where a given sample was removed from the dataset prior to differential expression analysis. To assess false positive results, cell type, subtype, or genotype labels were permuted before conducting the differential expression analysis (n = 1000 permutations).

#### Analysis of cell type identity shifts

To study shifts in cell type identities due to *Nr4a2* haploinsufficiency, classification and prediction of cell type identities based on the corrected Pearson residuals of modulated genes were performed using the *LinearSVC* function of the scikit-learn Python package version 1.3.2^93,94^. Only CLA, shell, and L6a neurons from the “neurons filtered” dataset were used in this analysis. Linear classifier parameters were optimized with *Nr4a2^wt/wt^* cells using cross-validated grid search via the *GridSearchCV* function of scikit-learn. The optimized parameters were “C” for regularization strength, “penalty” for the norm type, and “dual” for the optimization problem formulation. This search was conducted over 10-fold cross-validation iterations, and the performance of the cross-validated model was evaluated by computing the balanced accuracy, resulting in the following optimized values: C = 9e-04, penalty = “l2”, and dual = “auto”. The tolerance value for the stopping criteria was set to 1e-5, and the maximum number of iterations was set to 50’000. Linear classifier models were trained with the optimized parameters described above using 80% of the *Nr4a2^wt/wt^*cells, and cell type labels were predicted for the remaining 20% of *Nr4a2^wt/wt^*cells and all *Nr4a2^del/wt^* cells. To obtain a balanced partition of the data prior to model training, the *partition* function of the groupdata2 R package version 2.0.3 was used. Cell type classification performance was tested over 1’000 iterations. Significant differences between the genotypes in the percentage of classified cells across iterations for each actual-predicted cell type label pair were assessed using a generalized linear model (GLM)^95–97^ with a binomial distribution. Models were fitted via the glmmTMB R package version 1.1.9^98–100^, using Template Model Builder (TMB). No random effect term was specified. Likelihood ratio tests (LRT) with chi-square-based *p*-values were then computed by comparing the resulting models with null models where the fixed effects term (i.e. genotype) was dropped, using the *drop1* function of the stats R package^101^. To correct for multiple comparisons, *p*-values were adjusted using the Holm method^102^. To test for false negative results, permutation of cell identities was performed prior to model training, and accuracy differences between genotypes were compared to the observed results.

#### Gene regulatory network inference

To investigate alterations in transcription factor activity due to *Nr4a2* haploinsufficiency, gene regulatory network inference was conducted on the corrected, log1p-transformed counts of modulated genes using the pySCENIC Python package version 0.12.1^103,104^. Only CLA, shell, and L6a neurons from the “neurons filtered” dataset were used in this analysis. Motif2TF annotations and cisTarget databases generated with the 2022 SCENIC+^105^ motif collection were utilized (mm10 annotation files). The list of transcription factors was filtered to include only those modulated in *Nr4a2^del/wt^* mice. Genes lacking ranking values in the “genes versus motifs” ranking files were also excluded from the analysis. Gene regulatory networks (i.e. transcription factor-target gene co-expression modules) were inferred from the expression of modulated genes using the GENIE3 algorithm^106^ via the *genie3* function of the arboreto Python package version 0.1.6^103^. Regulons (i.e. gene targets of a given transcription factor) were then derived from the co-expression modules using the *modules_from_adjacencies* function of pySCENIC, with the following criteria: gene targets with importance values above the 90^th^ and 95^th^ percentiles were retained; the top 25 and 50 targets of each transcription factor were retained; and the targets for which the transcription factor is within its top 5, 10, and 50 regulators were retained. Activating and repressing regulons were inferred using a Pearson correlation coefficient threshold of 0.03, calculated using all cells. Only regulons containing at least 20 target genes were kept for further analysis. Regulons were pruned, and target genes with a cis-regulatory footprint for the corresponding transcription factor were retained using the “genes versus motifs” ranking files via the *prune2df* function of pySCENIC. Recovery curves were created using the top 500 ranked target genes of each transcription factor, and the area under the curve was calculated using 1% of the ranked genome. Regulons with a normalized enrichment score above 2.75 were retained, and regulon enrichment scores were calculated for each cell using the *aucell* function of pySCENIC. To identify regulons enriched in either *Nr4a2^wt/wt^* or *Nr4a2^del/wt^* cells, regulon specific scores (RSS)^57^ were calculated for each cell type and genotype using the *regulon_specificity_scores* function of pySCENIC, and the contrast in these scores was computed between genotypes.

#### Claustro-insular cellular composition analysis

Differences between genotypes in cell type proportions were tested using the *propeller* function of the speckle Bioconductor package^107^ version 1.0.0. This package implements functions from the limma Bioconductor package (version 3.56.2 was used)^108^. The mouse from which each cell was derived served as a biological replicate. A linear model was fitted to the logit-transformed proportions, and a robust empirical Bayes shrinkage^109^ of the variances was applied. Since there are two genotypes, a *t*-test was used to calculate *p*-values, which were adjusted for multiple comparisons using the Benjamini-Hochberg false discovery rate method^79^.

### MERFISH

#### MERFISH gene panel design

To construct the gene panel for MERFISH^110^ experiments, we prioritized the selection of marker genes that effectively segregate distinct cell populations in the mouse claustro-insular region. Marker genes were selected if they 1) consisted in gene makers of a major claustro-insular cell type, 2) corresponded to differentially expressed genes allowing the segregation of a pair of related cell types, 3) corresponded to previously described markers of the claustro-insular region, or 4) corresponded to variable genes in the single-cell RNA sequencing data. To this list were added generic neuronal and non-neuronal markers, such as *Slc17a7*, *Slc32a1*, *Pvalb*, *Sst*, *Vip*, and *Gfap*. A set of 500 genes was submitted to the Vizgen gene panel portal. Genes with insufficient target regions and genes exceeding the abundance threshold were removed from the panel. In order to avoid optical crowding, additional genes with high abundance were removed from the panel. A final set of 300 genes was used for MERFISH experiments.

#### Sample preparation

Approximately 12-week old mice (n = 1 C57BL6 male mouse, n = 3 *Nr4a2^wt/wt^* male mice and n = 3 *Nr4a2^del/wt^* male mice) were used for MERFISH experiments. Mice were anesthetized with 4% isoflurane (Attane^TM^, Piramal Healthcare) and killed by cervical dislocation. Brains were immediately removed and placed in a mold containing OCT (ref. KMA-0100-00A, CellPath Ltd), which was placed into an isopentane and liquid nitrogen bath. Once frozen, the tissue block was stored at −80°C until sectioning.

#### Cryosections

Brains were cut on a Leica CM3050 cryostat (Leica Biosystems). The cryostat chamber temperature was set at −21°C and the sample temperature at −16°C. Brains were placed in the cryostat for at least 30 minutes before starting sectioning. The OCT block was trimmed until reaching the beginning of the claustro-insular region. Vertical and horizontal alignment was verified using the Allen Mouse Brain Atlas (mouse.brain-map.org and atlas.brain-map.org). 10 μm thick sections were cut approximately at bregma level + 1.045 and mounted onto MERSCOPE slides (ref. PN 20400001, Vizgen). Slides were air dried inside the cryostat for 10-15 minutes. For RNA quality control, a few additional subsequent sections were cut and immediately placed in 350 ul RLT Plus lysis buffer (ref. 74034, RNeasy Plus Micro Kit, Qiagen) and kept at 4°C until analysis.

#### RNA extraction and quality control

RNA was extracted using the RNeasy Plus Micro Kit (ref. 74034, Qiagen) and following to the manufacturer’s protocol. Final elution was done with 14 μl of RNAse-free water at 14000 g. RNA integrity was assessed with the 2100 Bioanalyzer System (RNA Nano Chips, ref. 5067-1511, Agilent technologies,). For all samples analyzed, RIN was 10 and DV200 between 98-100%, indicating good RNA integrity.

#### MERFISH sample preparation and hybridization

MERSCOPE Slides with sections were prepared following the manufacture’s instructions. Briefly, slides were placed in a petri dish and fixed in freshly prepared 4% PFA in 1x PBS for 15 minutes at room temperature. Slides were washed in 1x PBS covered with 70% EtOH. Petri dishes were closed and sealed with parafilm (ref. PM-996, Parafilm) and stored at 4°C until hybridization (maximum 6 weeks). To stain the tissue with MERFISH probes, the manufacturer’s instructions were followed. For each processed brain sample, a sample verification was performed on a separate slide using the Sample Verification Kit (ref. 10400008, Vizgen). Hybridization procedure was the same for verification and experimental samples. Slides were washed and incubated for 30 minutes at 37°C in Formamide Wash Buffer (ref. 20300002, Vizgen) in an incubator. After removing Formamide buffer by aspiration, 50 μl of Mouse Verification Gene Probes (ref. 20300024, Vizgen) or 50 μl of the custom gene panel mix were added onto the sample, a piece of parafilm was placed onto the droplet to equally distribute the liquid. The petri dish was closed, sealed with parafilm, and incubated for 18-24 hours at 37°C in a humidified incubator. Slides were incubated twice in Formamide Wash Buffer for 30 minutes at 47°C. Slides were washed with Sample Prep Wash Buffer for 2 minutes. A Gel Coverslip (ref. 20400003, Vizgen) was coated with 100 ul of Gel Slick Solution. Wash buffer was removed from the sample and it was covered with a thin layer of Gel Embedding Solution (ref. 20300004, Vizgen) containing 10% ammonium persulfate solution and N,N,N’,N’-tetramethylethylenediamine. The Gel Coverslip was placed on top of the sample, which was incubated at room temperature for approximately 90 minutes, until the gel was solidified. For tissue clearing, after removing the Gel Coverslip, slides were incubated in 5 ml warmed Clearing Solution (ref. 20300003, Vizgen) with Proteinase K (ref. P8107S, NEB) in a sealed petri dish for 24 hours at 37°C.

#### MERFISH imaging

Sections were washed twice for 5 minutes in Sample Prep Wash Buffer and incubated for 15 minutes on a rocker with either 3 ml of Verification Staining Reagent (ref. 20300014, Vizgen, for verification samples) or 3 ml of DAPI and PolyT Staining Reagent (ref. 20300021, Vizgen, for experimental samples). Sections were washed with 5 ml of Formamide Wash Buffer for 10 minutes and then placed in Sample Prep Wash Buffer. For verification samples, slides were placed in the MERSCOPE Flow Chamber and Imaging Buffer (ref. 20300016, Vizgen) with Verification Imaging Buffer Activator (ref. 20300015, Vizgen) was injected into the chamber, which were inserted into the MERSCOPE instrument (ref. 10000001, Vizgen) and imaged according to the manufacturer’s protocol. For imaging of samples labeled with the 300 genes panel, the MERSCOPE Imaging Cartridge (ref. 20300018, Vizgen) was prepared by injecting 250 μl of Imaging Buffer activator (ref. 20300022, Vizgen) mixed with 100 μl of RNase inhibitor (ref. M0314L, NEB) into the cartridge. Next, 15 ml of mineral oil (ref. M5904-500ML, Sigma) were pipetted on top of the Imaging Activation Mix. The cartridge was inserted into the MERSCOPE instruments and the fluidics system was primed according to instructions. The sample was placed in the Flow Chamber, inserted into the MERSCOPE instrument and connected to fluidics system. After acquiring a low-resolution image of DAPI labeling, the region of interest was drawn and imaging was initiated. Sections were imaged with a 60x lens and 7 planes on the z-axis.

### MERFISH data analysis

#### MERFISH data preprocessing

Raw images of entire coronal sections were automatically processed by the MERSCOPE Instrument software, generating cell-by-gene and cell-metadata output matrices for each sample. Regions of interest, including cortical deep layers, endopirifom nucleus, piriform cortex, and striatum, were selected around the claustro-insular regions of each section for downstream analyses using R version 4.3.1. The number of expressed genes and mRNA molecules was computed for each cell. Cells with a volume of 500 µm^3^ to 3500 µm^3^ (inclusive) and containing at least 10 but no more than 3000 mRNA molecules, were retained for further analysis. No genes were filtered for downstream analysis. A variance stabilizing transformation was applied to the raw counts using the *SCTransform* function of Seurat^81,82^, as described in *scRNAseq data analysis, Data normalization and confounder regression*, with the following differences: Pearson residuals were clipped to a range of [-10, 10], and the effect of cell size on the number of mRNA molecules was corrected by regressing out cell volume. For visualizing the 2D maps of the brain, the rotation angle for each section was visually estimated, and the rotated coordinates were computed using a rotation matrix.

#### Projection of MERFISH data on the scRNAseq data

To identify cell types in the MERFISH dataset, a reference-based mapping approach was employed, where cell types were predicted using the scRNAseq dataset as a reference. To build this reference, supervised PCA (SPCA)^111^ was performed on the corrected Pearson residuals of the MERFISH panel genes expressed in the “neurons filtered” dataset (296 genes were used) using the *RunSPCA* function of Seurat. The linear matrix factorization during SPCA calculation was supervised using the SNN graph from the “neurons filtered” dataset. Only the first 100 SPCs were computed, and the top relevant SPCs were selected using the *KneeLocator* function of kneed (see *scRNAseq data analysis, Dimensionality reduction, dataset integration, and unsupervised graph-based clustering*). An Azimuth reference was created using the variance-stabilized data, the top SPCs, and the UMAP model from the “neurons filtered” dataset via the *AzimuthReference* function of the Azimuth R package version 0.5.0^53^. The cell type and subtype identities of cells were used for prediction in the MERFISH dataset. The variance-stabilized data of the MERFISH dataset was then projected onto the scRNAseq reference dataset, and cell type and subtype label prediction was performed via the *RunAzimuth* function of Azimuth.

#### Claustro-insular cellular composition analysis

Differences between genotypes in cell type proportions were tested using the *propeller* function of speckle, as described in *scRNAseq data analysis, Claustro-insular cellular composition analysis*, with the following difference: the region of interest from which each cell was derived served as a biological replicate.

### RNAscope smFISH

#### RNAscope sample preparation and hybridization

Mice (see Supplementary Tables for details) were anesthetized with isoflurane and killed by cervical dislocation. Brains were immediately removed, embedded in OCT and frozen in an isopentane bath in liquid nitrogen. Samples were stored at −80°C until sectioning. Brains were cut on a cryostat microtome in 10 μm coronal sections and mounted on Superfrost^TM^ Plus slides (ref. J1800AMNZ, Erpredia). Sections were dried for 1-2 hours at −20°C inside the cryostat, before being transferred for storage at −80°C. Sections were labeled by using the RNAscope™ Multiplex Fluorescent V2 Assay (ref. 323136, Advanced Cell Diagnostics). RNAscope staining was performed following the manufacture’s guidelines for fresh-frozen tissue samples. Before probe hybridization, sections were post-fixed in Formalin 10% for 15 minutes at 4°C, dehydrated, treated with hydrogen peroxide for 10 minutes at room temperature and treated with Protease IV for 15 minutes at room temperature. Sections were labeled with sets of 2-3 probes (see Supplementary Tables, smFISH labeled sections). Probes were hybridized for 2 hours at 40°C in a humidified chamber. Probe signal was developed with TSA Vivid Fluorophores 520 (1:750, ref. 323271), 570 (1:1000-1500, ref. 323272) and 650 (1:1500-2000, ref. 323273), diluted in TSA buffer. Sections were counterstained with DAPI and mounted with ProLongTM Gold antifade (ref. P36935, Invitrogen^TM^). Slides were imaged with a Nikon Ti/CSU-W1 Spinning Disc Confocal microscope using 405, 488, 561 and 640 nm excitation lasers and a 60x 1.49 NA air objective. Image tiles covering the claustro-insular region and of 6 planes on the z-axis were acquired, merging and orthogonal projection was applied within the SlideBook software (Intelligent Imaging Innovations).

#### RNAscope image analysis

Images were analyzed using QuPath (version 0.4.3). Cells were selected by drawing a region of interest (ROI) (see Table 1 for details). Cell segmentation based on DAPI staining was performed using the *Cell detection* function. The cell expansion around the detected nucleus was of 2 μm for analyses shown in Fig. 2 and Fig. 3 (minimizing false positive detection), and of 4 μm for analyses shown in Fig. 4 and Fig. 6 (maximizing detection of all transcripts). In the particular case of *Nr2f2*, only puncta within the cytoplasm were quantified, as only cytoplasmic *Nr2f2* mRNA is specifically enriched in the CLA. Next, puncta corresponding to individual mRNA molecules were automatically detected and quantified using the *Subcellular detection* function. For each fluorophore, a detection threshold was set, and all puncta above this threshold were counted. When multiple puncta were clustered together, an estimation of the number of individual puncta within the cluster was performed automatically by the *Subcellular detection* function. The same parameters were applied on all images from the same experiment. When referring to one section, this corresponds to an image of one claustro-insular region.

### RNAscope data analysis

#### Cluster identification

A custom R script was used for the analysis of RNAscope data on R version 4.3.1. Cells not expressing any of the three genes tested were removed from the data. Two transformation steps were then applied to the count data: 1) to account for outlier cells, the number of puncta per cell was log-transformed, 2) to eliminate gene-specific biases in the clustering of cells, gene-wise standardization of the data was applied by scaling and centering the numbers of puncta of a given gene across cells, using the *scale* function of the base R package. Euclidean distances were then computed between each pair of cells using the *dist* function of the stats R package. The data was finally clustered using Ward’s minimum variance agglomeration method implemented in the *hclust* function of the fastcluster R package version 1.2.6 (with method = “ward.D2”)^112–114^. This package provides a fast implementation of hierarchical agglomerative clustering. The resulting hierarchical clustering tree was iteratively cut into 2 to 12 groups using the *cutree* function of the stats R package. The optimal number of groups was decided through visual inspection of the results (i.e. gene expression across groups, typically using violin plots^115^) and was guided by the single-cell RNA-seq data. Clustering was performed for each probe set separately. The number of mice, images, and cells varied for each experiment and are provided in Supplementary Tables, smFISH data. When the experiment included mice with different genotypes, the data was clustered together and no correction for the genotype effect was performed. When the probe set was designed to identify cell subtypes in the RNAscope data, a second round of clustering was necessary. This involved identifying the group (or groups) containing the desired cells (identified based on their gene expression profiles), and a second round of clustering only on these cells was applied following the steps described above. Right (medial-lateral) and left (lateral-medial) claustro-insular images were used in all experiments and were clustered together per probe set. However, these images did not always respectively correspond to sections from the right and left side of the brain.

#### Rigid image registration

Multiple images were used to build the claustro-insular maps in Fig. 2, Fig. 6 and Supplementary Fig. 4. For each antero-posterior level, a representative image was chosen as reference image, and all other images from the same position were registered to this reference. Horizontal flipping of left images was performed to align all images to the right side. A reference point located between the CLA and the dorsal EP and adjacent to the external capsule was chosen for each image for image centering. Two other landmark points were also chosen for landmark-based image rotation. The relative rotation angle of each image with the reference image was then computed through slope-based angle calculation. 2D coordinate rotation using a rotation matrix was then conducted as the final step of image registration. No scaling of images was performed.

#### Claustro-insular cell type map generation

RNAscope 2D coordinates of analyzed cells were used for the generation of a map of major claustro-insular cell types. As the claustrum extends over several millimeters along the anteroposterior axis of the brain, four representative coronal sections were used for map generation (bregma +1.42, +1.045, +0.145 and −1.055). Two different probe sets were used: probe sets A and B (see Supplementary Tables, smFISH data). The number of mice, images and cells analyzed are described in Supplementary Tables, smFISH data. The data from each probe set was clustered separately as described in the main Methods section (*RNAscope data analysis, Cluster identification.* For details about the identified clusters, see the results section of the manuscript). However, the data from the two probe sets were projected onto the same map following the steps described in *RNAscope image analysis, Rigid image registration*. A reference image was chosen for each antero-posterior level, and they all correspond to probe set A images. The landmark points used for slope-based angle calculation were chosen differently depending on the brain level. For bregma +1.42 to +0.145 images, these points were located at the dorsal and ventral tips of the EP. For bregma −1.055 images, the points were located at the dorsal and ventral tips of the claustrum-like L6a region. Two bregma −1.055 images that were used for clustering were excluded from map generation: one was anterior to the other images, and the second was oblique. Both distorted the final map. The cluster characterized by high expression levels of *Nr4a2* but no expression of *Slc17a6* from probe set B was also excluded from map generation, because both CLA and L6b neurons exhibit high expression levels of *Nr4a2* and as this probe set did not include a probe for *Ccn2* (also known as *Ctgf*), a marker gene for L6b neurons, this cluster constituted a mixed set of neuronal populations and hence distorted the final map. Nevertheless, we were able to identify using this probe set a cluster corresponding to CLA projection neurons characterized by its expression of high level of *Nr4a2* and *Slc17a6*. To highlight the diversity of cell types constituting the mouse claustro-insular region, proportional visualizations of the maps were created (Fig. 2e and Supplementary Fig. 4b). The 2D spatial maps (where each dot corresponds to a cell from a given cluster) were binned into 20 μm bins. The proportion of cells from each cluster was then calculated for each bin, and each bin was assigned a color weighted by the cluster proportions in that bin. To visualize the density distribution of CLA, shell, *Ccn2^+^*, and *Rprm^+^* cells, 2D density maps of these clusters were generated for each brain level (Fig. 2f-j and Supplementary Fig. 4c). The two-dimensional kernel density estimation was computed using the *kde2d* function of the MASS R package^77^ version 7.3.60.0.1, implemented in the *stat_density_2d_filled* function of the ggplot2 R package version 3.5.0. The density estimates were scaled to a maximum of 1 with contour breaks ranging from 0.1 to 1 in increments of 0.1. The number of grid points in the dorso-ventral and medio-lateral directions was set to 100, and the bandwidth was estimated using the normal reference distribution via the *bandwidth.nrd* function of the MASS package. The maps were finally generated by plotting the highest density region of CLA, shell, *Ccn2^+^*, and *Rprm^+^*cells at 50% probability level of the two-dimensional kernel density estimate using the *stat_hdr* function of the ggdensity R package version 1.0.0. All the maps were scaled to the same coordinate space.

#### Spatial alterations in Nr4a2 haploinsufficient mice

To visualize spatial alterations in the claustro-insular region of *Nr4a2* haploinsufficient mice a map of their CLA and shell was produced following the same method as described above. Only coronal sections from bregma level +1.045 were used. Two different probe sets were used (see Supplementary Tables, smFISH data). The number of mice, images and cells analyzed per genotype are described in Supplementary Tables, Statistics table. The steps leading to the spatial projection of cells in 2D (Fig. 6g) and the binning of the map into 20 μm bins are as described above. The normalized CLA/shell contrast (i.e. the relative difference between the two clusters) was computed for each bin and is defined as:

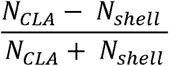

where N is the number of cells from the corresponding cluster.

#### RNAscope statistical analyses

All statistical analyses of RNAscope data were performed using R. Differences between genotypes in the total number of cells (Fig. 6a), number of *Nr4a2^+^* cells (Fig. 6b), number of *Nr4a2* puncta (Fig.4c), and the expression levels of modulated genes (Fig. 4g-j) were tested using generalized linear mixed models (GLMM) with negative binomial distributions, quadratic parameterization^116^, and a log link function (with family = nbinom2(link = “log”)). Models were fitted via the glmmTMB^98–100^ R package version 1.1.9, using Template Model Builder (TMB). Probe sets and/or images were included as random effects depending on the test, as detailed in Supplementary Tables, Statistics table. Likelihood ratio tests (LRT) with chi-square-based *p*-values were then computed by comparing the resulting models with null models where the fixed effects term (i.e. genotype) was dropped, using the *drop1* R function^101^. When testing for differences in the expression levels of modulated genes between genotypes, *p*-values were adjusted for multiple comparisons using the Holm method^102^. Differences between genotypes in cell type proportions were tested using the *propeller* function of speckle, as described in *scRNAseq data analysis, Claustro-insular cellular composition analysis*, with the following difference: the image from which each cell was derived served as a biological replicate.

### Electrophysiological recordings

#### Slice recordings

Slice recordings were conducted on 16- to 23-week-old Vglut2-IRES-cre;*Nr4a2^WT/WT^* and Vglut2-IRES-cre;*Nr4a2^Del/WT^* mice, infected with retrogradely transported adeno-associated viral particles AAV(rg)-hSyn-DIO-EGFP (Addgene #50457) in the mPFC 3-4 weeks prior. In brief, mice were anesthetized with isoflurane (3–5% induction, 1–2% maintenance), and the skin over the skull was removed under local anesthesia using carbostesin (AstraZeneca). The animals were head-fixed in a stereotaxic apparatus (Stoelting) using ear bars and a nose clamp, with their eyes protected by artificial tears. Body temperature was maintained at ∼37°C via a heating pad (FHC). Bilateral craniotomies above the mPFC were performed using an air-pressurized drill. A Glass pipette filled with AAV solution was inserted one hemisphere at a time at the following coordinates relative to bregma: AP: 1.7 mm, ML: ±0.3 mm, and DV: □0.75 & □1.75 mm. A volume of 100–200 nl was injected per site. After the procedure, mice were allowed to recover for 3–4 weeks.

For slice recordings, animals were anesthetized with isoflurane, and the brain was quickly extracted. Coronal slices, 300 μm thick, were cut using a vibratome (Leica VT 928 S1000, Germany). The slices were transferred to continuously oxygenated artificial cerebrospinal fluid (ACSF) at 37°C for 30 minutes, then maintained at room temperature for the remainder of the experiment. The ACSF contained (in mM): 124 NaCl, 3 KCl, 2 CaCl_2_, 1.3 MgSO_4_, 26 NaHCO_3_, 1.25 NaH_2_PO_4_, 10 D-glucose, with an osmolarity of 300 mOsm and a pH of 7.4 when bubbled with 95% O_2_-5% CO_2_. The slices were transferred to the recording chamber and perfused with oxygenated ACSF. All recordings were performed at ∼37°C (using a recording chamber and temperature controller from Luigs & Neumann, Germany). Claustral neurons projecting to the mPFC and expressing the viral construct were identified using a 473 nm LED (Thorlabs, Germany). Neurons were visualized using an IR-DIC microscope (Olympus BX51, Germany) and whole-cell recordings of identified neurons were performed using borosilicate glass pipettes with a resistance of 4–7 MΩ, filled with an intracellular solution containing (in mM): 120 K-gluconate, 10 KCl, 10 HEPES, 4 ATP, 0.3 GTP, 10 phosphocreatine, 1 EGTA, and 0.4% biocytin. The solution had a pH of 7.2-7.3 and an osmolarity of 280-290 mOsm. The signal was amplified using Multiclamp 700A amplifiers (Molecular Devices, USA) with a sampling rate of 50 kHz, high-pass filtered at 4 kHz, and digitized using the PulseQ electrophysiology package running on Igor Pro (Wavemetrics, USA). Neurons were recorded either in ACSF or in ACSF supplemented with various pharmacological agents. Fast GABAergic and glutamatergic transmission was blocked using SR 95531 hydrobromide (GBZ, 10 μM; Tocris), NBQX-disodium salts (10 μM; Tocris), and DL-AP5 (100 μM; Tocris). ATX-II (10 nM, Alomone Labs) is an enhancer of the resurgent current of voltage-gated sodium channels. Kv1-4 Channels were blocked with 4-aminopyridine (1 mM, Tocris), Kv7 channels with XE991 dihydrochloride (20 µM, Alomone Labs), and IRK channels with barium chloride (1 mM, Sigma-Aldrich). Finally, the mechanism underlying calcium-induced calcium release modulation of BK channels was dissected using the following pharmacological channel blockers: Cav1: Nifedipine (10 μM), Cav2.1: ω-Agatoxin IVA (100 nM, Alomone Labs), Cav2.2: ω-Conotoxin GVIA (3 μM, Alomone Labs), Cav2.3: SNX-482 (400 nM, Alomone Labs), Cav3: ML218 (2 μM, Alomone Labs), BKCa channels: Iberiotoxin (100 nM, Alomone Labs), RyR2: Ryanodine (10-20 μM, Alomone Labs).

#### Patch clamp data analysis

To analyze patch clamp data, waveforms acquired in IgorPro6 were converted to MATLAB files. Action potential (AP) peaks were detected with MATLAB’s built-in ‘findpeaks’ function. To identify AP onsets, we computed the rolling average and rolling standard deviation over a window of 30 time points. An onset was defined when the absolute difference between the membrane potential and the rolling average exceeded four times the rolling standard deviation, and the potential was above the rolling average. The half-width was measured at half the AP’s amplitude. Offsets were identified after the peak as the point closest to the onset potential. Rise time was calculated from onset to peak, decay time from peak to offset, and the after-hyperpolarization potential (AHP) was measured from the offset to the lowest potential following the offset.

The area under the curve (AUC) for each cell was calculated as the total surface area under the input/output function curve, using GraphPad Prism 8. To better compare the change in firing frequency between genotypes under different pharmacological conditions, the AUC and AHP amplitude were normalized to the wild-type values. For each pharmacological condition, we calculated the difference between a cell AUC and the mean *Nr4a2^wt/wt^* AUC, then divided this difference by the mean AUC of the control *Nr4a2^wt/wt^* cells for that condition (data presented as a percentage change). For each pharmacological condition, AHP amplitude was normalized by dividing each cell AHP amplitude by the mean *Nr4a2^wt/wt^* AHP amplitude for that condition. Statistical analyses were performed using GraphPad Prism 8.

Several genes encoding different types of sodium and potassium channels were dysregulated in *Nr4a2* haploinsufficient mice (Fig. 5a). The Nav1.6 α subunit and the auxiliary β1 subunit of voltage-gated sodium channels (VGSC, encoded by *Scn8a* and *Scn1b*, respectively) were upregulated and downregulated in haploinsufficient mice, respectively (Fig. 5a and Supplementary Fig. 10a). β subunits play a crucial role in VGSC trafficking and modulate their biophysical properties, such as the resurgent sodium current. A reduction of *Scn1b* expression may lead to a reduced resurgent current and decreased neuronal firing^117,118^. However, neither the kinetics of action potential depolarization (Fig. 5h) nor the pharmacological modulation of the resurgent sodium current differed between genotypes (Supplementary Fig. 10b), suggesting that the alterations in firing properties did not originate from changes in VGSC function.

We investigated whether changes in the expression of genes encoding voltage-gated potassium channels (VGPCs, Kv) could explain the difference in firing properties. Notably Kv3.1 (encoded by *Kcnc1*) belongs to a family of potassium channels that facilitate fast spiking by helping in action potential repolarization^119^. Although reduced *Kcnc1* expression would be expected to prolong the repolarization phase, we observed the opposite effect (Fig. 5h). *Kcnv1* (Kv8.1) and *Kcnf1* (Kv5.1) do not form functional homomeric channels but instead assemble with Kv2 channels to modulate their biophysical properties^120^ and sensitivity to the antagonist 4-aminopyridine^121,122^. SCN1B can also form complexes with various VGPCs, including Kv1, Kv4.2 and Kv7^123,124^. The antagonist of Kv1-4 channels, 4-aminopyridine, caused strong changes in firing properties in wild-type cells but had no effect on haploinsufficient neurons (Supplementary Fig. 10c,d). Additionally, Kv7 antagonist altered the shape of the input/output functions but maintained a similar difference between genotypes (Supplementary Fig. 10e). These findings suggest that the changes in firing properties cannot be easily attributed to alterations in VGPC function.

Finally, GIRK1 and GIRK 3 (encoded by *Kcnj3* and *Kcnj4*, respectively), which were upregulated in haploinsufficient mice, are G-protein-activated inward rectifier potassium channels that can reduce neuronal excitability^125^. Barium, a broad-spectrum blocker of inward rectifier channels, altered the shape of the input/output functions but left a similar difference between genotypes (Supplementary Fig. 10f).

## Data availability

Data will be publicly available upon publication of the manuscript.

## Code availability

All codes used for data analysis will be publicly available upon publication of the manuscript.

## Acknowledgements

We thank all members of I.R and A.C laboratories for helpful discussions, Francisco Resende for expert mouse care, Loris Mannino and Leonardo Marconi for technical assistance, Dr. Jose Manuel Nunes for statistical advice, the Photonic Bioimaging Center of the University of Geneva for assistance with imaging, the NeuroNA Human Cellular Neuroscience Platform of Campus Biotech Geneva for assistance with MERFISH spatial transcriptomics, the Flow Cytometry core facility and the iGE3 Genomics Platform at the Faculty of Medicine of the University of Geneva for expert technical assistance for the scRNAseq experiment. We warmly thank Dr. Hongkui Zeng (the Allen institute) for generously providing the *Nr4a2*^tm1(dreo)Hze^ mice.

## Author contributions

A.C. and I.R. acquired funding for the research. L.F., M.B., M.M., A.C. and I.R. conceived and designed the experiments. L.F., M.B. and M.M. acquired and analyzed data. L.F., M.B., M.M., A.C. and I.R., interpreted the data, wrote and edited the manuscript.

