## Supplementary Tables for "A spatial single-cell atlas of the claustro-insular region uncovers key regulators of neuronal identity and excitability"

| Figure | Parameter | Sample size | Statistical test | Random effect | Statistical data |
|---|---|---|---|---|---|
| Fig. 4c | log2 ratio mean number Nr4a2 mRNA puncta/cell (Nr4a2(del/wt)/Nr4a2(wt/wt)) | Nr4a2(wt/wt) n = 55697 cells, 26 sections, 4 mice; Nr4a2(del/wt) n = 71389 cells, 34 sections, 4 mice. | chi-square-based LRT applied to a negative binomial GLMM with quadratic parametrization | probeset and image | estimate = -1.03, LRT = 85.26, p-value = 2.62e-20 |
| Fig. 4g | log2 ratio mean ratio number mRNA puncta/cell (Nr4a2(del/wt)/Nr4a2(wt/wt)) | Cdh13 & Rxfp: Nr4a2(wt/wt) n = 12850 cells, 8 sections, 3 mice; Nr4a2(del/wt) n = 17503 cells, 11 sections, 3 mice. Scn1b: Nr4a2(wt/wt) n = 11310 cells, 7 sections, 2 mice; Nr4a2(del/wt) n = 7702 cells, 5 sections, 2 mice. Nr2f2: Nr4a2(wt/wt) n = 6746 cells, 4 sections, 2 mice; Nr4a2(del/wt) n = 7717 cells, 5 sections, 2 mice. | chi-square-based LRT applied to a negative binomial GLMM with quadratic parametrization; p-values were adjusted for multiple comparisons using the Holm method | image | Cdh13: estimate = -0.297, LRT = 19.63, p-value = 3.76e-05; Nr2f2: estimate = -0.61, LRT = 21.10, p-value = 2.62e-05; Rxfp1: estimate = -0.572, LRT = 29.64, p-value = 3.63e-07; Scn1b: estimate = -0.233, LRT = 7.76, p-value = 0.0107 |
| Fig. 4h | log2 ratio mean ratio number mRNA puncta/cell (Nr4a2(del/wt)/Nr4a2(wt/wt)) | Syt17: Nr4a2(wt/wt) n = 42912 cells, 14 sections, 3 mice; Nr4a2(del/wt) n = 45572 cells, 15 sections, 3 mice. Ntm: Nr4a2(wt/wt) n = 18367 cells, 6 sections, 2 mice; Nr4a2(del/wt) n = 24504 cells, 8 sections, 2 mice. Ryr2: Nr4a2(wt/wt) n = 24545 cells, 8 sections, 2 mice; Nr4a2(del/wt) n = 21068 cells, 7 sections, 2 mice. | chi-square-based LRT applied to a negative binomial GLMM with quadratic parametrization; p-values were adjusted for multiple comparisons using the Holm method | image | Ntm: estimate = 0.331, LRT = 14.34, p-value = 0.000459; Ryr2: estimate = 0.25, LRT = 20.61, p-value = 2.82e-05; Syt17: estimate = 0.186, LRT = 7.22, p-value = 0.0107 |
| Fig. 4i | mean number of Ntm mRNA puncta/cell | Nr4a2(wt/wt) n = 18367 cells, 6 sections, 2 mice; Nr4a2(del/wt) n = 24504 cells, 8 sections, 2 mice. | chi-square-based LRT applied to a negative binomial GLMM with quadratic parametrization; p-values were adjusted for multiple comparisons using the Holm method | image | CLA: estimate = 0.544, LRT = 33.41, p-value = 2.98e-08; Ntm+: estimate = 0.00993, LRT = 0.13, p-value = 1; Shell: estimate = 0.0627, LRT = 4.69, p-value = 0.0908; Syt17+: estimate = -0.02, LRT = 0.40, p-value = 1 |
| Fig. 4j | mean number of Ryr2 mRNA puncta/cell | Nr4a2(wt/wt) n = 24545 cells, 8 sections, 3 mice; Nr4a2(del/wt) n = 21068 cells, 7 sections, 3 mice. | chi-square-based LRT applied to a negative binomial GLMM with quadratic parametrization; p-values were adjusted for multiple comparisons using the Holm method | image | CLA: estimate = 0.321, LRT = 21.35, p-value = 1.53e-05; Ryr2+: estimate = -0.0148, LRT = 0.17, p-value = 1; Shell: estimate = 0.138, LRT = 7.60, p-value = 0.0175; Syt17+: estimate = -0.0134, LRT = 0.15, p-value = 1 |

| | | | | | |
|---|---|---|---|---|---|
| Fig. 5d | Firing frequency difference ACSF (Nr4a2(del/wt)/Nr4a2(wt/wt)) | Nr4a2(wt/wt) n = 31 cells, 7 mice; Nr4a2(del/wt) n = 43 cells, 10 mice. | two-way repeated measure ANOVA; post-hoc testing using Fisher's LSD test with 5% false discovery rate; p-values were adjusted for multiple comparisons using the Benjamini-Hochberg false discovery rate method | NA | [I(pA)]x[Genotype]: F(20,1440)=3.193, p-value<0.0001, [Genotype]:F(1,72)=13.69,p-value = 0.0004. 0pA: p-value>0.999, 25pA: p-value = 0.9661, 50pA:p-value = 0.7550,75pA:p-value = 0.2690,100pA:p-value = 0.0575, 125pA: p-value = 0.0240, 150pA: p-value = 0.0099, 175pA: p-value = 0.0068, 200pA: p-value = 0.0041, 225pA: p-value = 0.0020, 250pA: p-value = 0.0016, 275pA: p-value = 0.0014, 300pA: p-value = 0.0012, 325pA: p-value = 0.0012, 350pA:p-value = 0.0012, 375pA: p-value = 0.0012, 400pA: p-value = 0.0012, 425pA: p-value = 0.0012, 450pA: p-value = 0.0012, 475pA: p-value = 0.0014, 500pA: p-value = 0.0029 |
| Fig. 5d | Firing frequency difference Synaptic Blockers (Nr4a2(del/wt)/Nr4a2(wt/wt)) | Nr4a2(wt/wt) n = 16 cells, 3 mice; Nr4a2(del/wt) n = 10 cells, 2 mice. | two-way repeated measure ANOVA; post-hoc testing using Fisher's LSD test with 5% false discovery rate; p-values were adjusted for multiple comparisons using the Benjamini-Hochberg false discovery rate method | NA | [I(pA)]x[Genotype]: F(20,480)=3.490, p-value<0.0001, [Genotype]:F(1,24)=14.41, p-value = 0.00088. 0pA: p-value>0.9999, 25pA: p-value = 0.9842, 50pA:p-value = 0.6322,75pA:p-value = 0.2934,100pA:p-value = 0.1448, 125pA: p-value = 0.0742, 150pA: p-value = 0.0204, 175pA: p-value = 0.0067, 200pA: p-value = 0.0023, 225pA: p-value = 0.0010, 250pA: p-value = 0.0005, 275pA: p-value = 0.0003, 300pA: p-value = 0.0003, 325pA: p-value = 0.0003, 350pA:p-value = 0.0004, 375pA: p-value = 0.0010, 400pA: p-value = 0.0017, 425pA: p-value = 0.0022, 450pA: p-value = 0.0045, 475pA: p-value = 0.0098, 500pA: p-value = 0.0246 |
| Fig. 5e | AUC difference (Nr4a2(del/wt)/Nr4a2(wt/wt)) | ACSF : Nr4a2(wt/wt) n = 31 cells, 7 mice; Nr4a2(del/wt) n = 43 cells, 10 mice; Syn Bl: Nr4a2(wt/wt) n = 16 cells, 3 mice; Nr4a2(del/wt) n = 10 cells, 2 mice. | two-way ANOVA; post-hoc testing using Fisher's LSD test with 5% false discovery rate; p-values were adjusted for multiple comparisons using the two-stage linear step-up procedure of Benjamini, Krieger and Yekutieli | NA | [Pharmacology]x[Genotype]: F(1,96)=2.004, p-value = 0.1602, [Genotype]:F(1,96)=26.45, p-value<0.0001. Nr4a2(wt/wt)(ACSF)/Nr4a2(del/wt)(ACSF)=0.0003, Nr4a2(wt/wt)(SynBl)/Nr4a2(del/wt)(SynBl)=0.0003 |
| Fig. 5f | Resting membrane potential difference (Nr4a2(del/wt)/Nr4a2(wt/wt)) | ACSF : Nr4a2(wt/wt) n = 31 cells, 7 mice; Nr4a2(del/wt) n = 43 cells, 10 mice; Syn Bl: Nr4a2(wt/wt) n = 8 cells, 3 mice; Nr4a2(del/wt) n = 4 cells, 2 mice. | two-way ANOVA; post-hoc testing using Fisher's LSD test with 5% false discovery rate; p-values were adjusted for multiple comparisons using the two-stage linear step-up procedure of Benjamini, Krieger and Yekutieli | NA | [Pharmacology]x[Genotype]: F(1,82)=0.3201, p-value = 0.5731, [Genotype]:F(1,82)=0.02292, p-value = 0.8800. Nr4a2(wt/wt)(ACSF)/Nr4a2(del/wt)(ACSF)=0.7126, Nr4a2(wt/wt)(SynBl)/Nr4a2(del/wt)(SynBl)=0.7369 |

| | | | | | |
|---|---|---|---|---|---|
| Fig. 5h | Rise time difference (Nr4a2(del/wt)/Nr4a2(wt/wt)) | ACSF : Nr4a2(wt/wt) n = 31 cells, 7 mice; Nr4a2(del/wt) n = 43 cells, 10 mice; Syn Bl: Nr4a2(wt/wt) n = 16 cells, 3 mice; Nr4a2(del/wt) n = 10 cells, 2 mice. | two-way ANOVA; post-hoc testing using Fisher's LSD test with 5% false discovery rate; p-values were adjusted for multiple comparisons using the two-stage linear step-up procedure of Benjamini, Krieger and Yekutieli | NA | [Pharmacology]x[Genotype]: F(1,96)=0.04584, p-value = 0.8309, [Genotype]:F(1,96)=0.5773, p-value = 0.4492. Nr4a2(wt/wt)(ACSF)/Nr4a2(del/wt)(ACSF)=0.6195, Nr4a2(wt/wt)(SynBl)/Nr4a2(del/wt)(SynBl)=0.5741 |
| Fig. 5h | Decay time difference (Nr4a2(del/wt)/Nr4a2(wt/wt)) | ACSF : Nr4a2(wt/wt) n = 31 cells, 7 mice; Nr4a2(del/wt) n = 43 cells, 10 mice; Syn Bl: Nr4a2(wt/wt) n = 16 cells, 3 mice; Nr4a2(del/wt) n = 10 cells, 2 mice. | two-way ANOVA; post-hoc testing using Fisher's LSD test with 5% false discovery rate; p-values were adjusted for multiple comparisons using the two-stage linear step-up procedure of Benjamini, Krieger and Yekutieli | NA | [Pharmacology]x[Genotype]: F(1,96)=0.6560, p-value = 0.4200, [Genotype]:F(1,96)=11.49, p-value = 0.0010. Nr4a2(wt/wt)(ACSF)/Nr4a2(del/wt)(ACSF)=0.0003, Nr4a2(wt/wt)(SynBl)/Nr4a2(del/wt)(SynBl)=0.1163 |
| Fig. 5h | Amplitude difference (Nr4a2(del/wt)/Nr4a2(wt/wt)) | ACSF : Nr4a2(wt/wt) n = 31 cells, 7 mice; Nr4a2(del/wt) n = 43 cells, 10 mice; Syn Bl: Nr4a2(wt/wt) n = 16 cells, 3 mice; Nr4a2(del/wt) n = 10 cells, 2 mice. | two-way ANOVA; post-hoc testing using Fisher's LSD test with 5% false discovery rate; p-values were adjusted for multiple comparisons using the two-stage linear step-up procedure of Benjamini, Krieger and Yekutieli | NA | [Pharmacology]x[Genotype]: F(1,96)=0.1430, p-value = 0.7062, [Genotype]:F(1,96)=1.562, p-value = 0.2144. Nr4a2(wt/wt)(ACSF)/Nr4a2(del/wt)(ACSF)=0.0462, Nr4a2(wt/wt)(SynBl)/Nr4a2(del/wt)(SynBl)=0.2152 |
| Fig. 5h | Half width difference (Nr4a2(del/wt)/Nr4a2(wt/wt)) | ACSF : Nr4a2(wt/wt) n = 31 cells, 7 mice; Nr4a2(del/wt) n = 43 cells, 10 mice; Syn Bl: Nr4a2(wt/wt) n = 16 cells, 3 mice; Nr4a2(del/wt) n = 10 cells, 2 mice. | two-way ANOVA; post-hoc testing using Fisher's LSD test with 5% false discovery rate; p-values were adjusted for multiple comparisons using the two-stage linear step-up procedure of Benjamini, Krieger and Yekutieli | NA | [Pharmacology]x[Genotype]: F(1,96)=0.1913, p-value = 0.6628, [Genotype]:F(1,96)=8.730, p-value = 0.0039. Nr4a2(wt/wt)(ACSF)/Nr4a2(del/wt)(ACSF)=0.0018, Nr4a2(wt/wt)(SynBl)/Nr4a2(del/wt)(SynBl)=0.0938 |
| Fig. 5h | AHP Amplitude difference (Nr4a2(del/wt)/Nr4a2(wt/wt)) | ACSF : Nr4a2(wt/wt) n = 31 cells, 7 mice; Nr4a2(del/wt) n = 43 cells, 10 mice; Syn Bl: Nr4a2(wt/wt) n = 16 cells, 3 mice; Nr4a2(del/wt) n = 10 cells, 2 mice. | two-way ANOVA; post-hoc testing using Fisher's LSD test with 5% false discovery rate; p-values were adjusted for multiple comparisons using the two-stage linear step-up procedure of Benjamini, Krieger and Yekutieli | NA | [Pharmacology]x[Genotype]: F(1,96)=0.0515 p-value = 0.8209, [Genotype]:F(1,96)=11.13, p-value = 0.0012. Nr4a2(wt/wt)(ACSF)/Nr4a2(del/wt)(ACSF)=0.0057, Nr4a2(wt/wt)(SynBl)/Nr4a2(del/wt)(SynBl)=0.0585 |

| | | | | | |
|---|---|---|---|---|---|
| Fig. 5j | Firing frequency difference VGCC (Nr4a2(del/wt)/Nr4a2(wt/wt)) | Nr4a2(wt/wt) n = 9 cells, 1 mouse; Nr4a2(del/wt) n = 7 cells, 1 mouse. | two-way repeated measure ANOVA; post-hoc testing using Fisher's LSD test with 5% false discovery rate; p-values were adjusted for multiple comparisons using the Benjamini-Hochberg false discovery rate method | NA | [I(pA)]x[Genotype]: F(20,280)=0.1754, p-value>0.9999, [Genotype]:F(1,14)=0.0420, p-value = 0.8404. 0pA: p-value>0.999, 25pA: p-value>0.999, 50pA:p-value>0.999,75pA:p-value>0.999,100pA:p-value>0.999, 125pA: p-value>0.999, 150pA: p-value>0.999, 175pA: p-value>0.999, 200pA: p-value>0.999, 225pA: p-value>0.999, 250pA: p-value>0.999, 275pA: p-value>0.999, 300pA: p-value>0.999, 325pA: p-value>0.999, 350pA:p-value>0.999, 375pA: p-value>0.999, 400pA: p-value>0.999, 425pA: p-value>0.999, 450pA: p-value>0.999, 475pA: p-value>0.999, 500pA: p-value>0.999 |
| Fig. 5j | Firing frequency difference BK channels blocker (Nr4a2(del/wt)/Nr4a2(wt/wt)) | Nr4a2(wt/wt) n = 19 cells, 2 mice; Nr4a2(del/wt) n = 15 cells, 2 mice. | two-way repeated measure ANOVA; post-hoc testing using Fisher's LSD test with 5% false discovery rate; p-values were adjusted for multiple comparisons using the Benjamini-Hochberg false discovery rate method | NA | [I(pA)]x[Genotype]: F(20,640)=0.4497, p-value = 0.9825, [Genotype]:F(1,32)=0.0088, p-value = 0.9258. 0pA: p-value = 0.9950, 25pA: p-value = 0.9950, 50pA:p-value = 0.9950,75pA:p-value = 0.9950,100pA:p-value = 0.9950, 125pA: p-value = 0.9950, 150pA: p-value = 0.9950, 175pA: p-value = 0.9950, 200pA: p-value = 0.9950, 225pA: p-value = 0.9950, 250pA: p-value = 0.9950, 275pA: p-value = 0.9950, 300pA: p-value = 0.9950, 325pA: p-value = 0.9950, 350pA:p-value = 0.9950, 375pA: p-value = 0.9950, 400pA: p-value = 0.9950, 425pA: p-value = 0.9950, 450pA: p-value = 0.9950, 475pA: p-value = 0.9950, 500pA: p-value = 0.9950 |
| Fig. 5j | Firing frequency difference Ryanodine blocker (Nr4a2(del/wt)/Nr4a2(wt/wt)) | Nr4a2(wt/wt) n = 26 cells, 3 mice; Nr4a2(del/wt) n = 24 cells, 3 mice. | two-way repeated measure ANOVA; post-hoc testing using Fisher's LSD test with 5% false discovery rate; p-values were adjusted for multiple comparisons using the Benjamini-Hochberg false discovery rate method | NA | [I(pA)]x[Genotype]: F(20,960)=0.5490, p-value = 0.9457, [Genotype]:F(1,48)=2.912, p-value = 0.0944. 0pA: p-value>0.9999, 25pA: p-value>0.9999, 50pA:p-value>0.9999,75pA:p-value = 0.9171,100pA:p-value = 0.6978, 125pA: p-value = 0.4549, 150pA: p-value = 0.3868, 175pA: p-value = 0.3868, 200pA: p-value = 0.3868, 225pA: p-value = 0.4999, 250pA: p-value = 0.4999, 275pA: p-value = 0.3868, 300pA: p-value = 0.3868, 325pA: p-value = 0.3868, 350pA:p-value = 0.3868, 375pA: p-value = 0.3868, 400pA: p-value = 0.3868, 425pA: p-value = 0.3868, 450pA: p-value = 0.3868, 475pA: p-value = 0.3868, 500pA: p-value = 0.3868 |
| Fig. 5k | % change in AUC - ACSF (Nr4a2(del/wt)/Nr4a2(wt/wt)) | Nr4a2(wt/wt) n = 31 cells, 7 mice; Nr4a2(del/wt) n = 43 cells, 10 mice. | Mann-Whitney U test | NA | Sum of ranks (Nr4a2(wt/wt)) = 1483, Sum of ranks (Nr4a2(del/wt)) = 1292, p-value = 0.0003 |
| Fig. 5k | % change in AUC - Synaptic blockers (Nr4a2(del/wt)/Nr4a2(wt/wt)) | Nr4a2(wt/wt) n = 16 cells, 3 mice; Nr4a2(del/wt) n = 10 cells, 2 mice. | Mann-Whitney U test | NA | Sum of ranks (Nr4a2(wt/wt)) = 276, Sum of ranks (Nr4a2(del/wt)) = 75, p-value = 0.0009 |
| Fig. 5k | % change in AUC - VGCC (Nr4a2(del/wt)/Nr4a2(wt/wt)) | Nr4a2(wt/wt) n = 9 cells, 1 mouse; Nr4a2(del/wt) n = 7 cells, 1 mouse. | Mann-Whitney U test | NA | Sum of ranks (Nr4a2(wt/wt)) = 79, Sum of ranks (Nr4a2(del/wt)) = 57, p-value = 0.8371 |

| | | | | | |
|---|---|---|---|---|---|
| Fig. 5k | % change in AUC - Cav2&Cav3 (Nr4a2(del/wt)/Nr4a2(wt/wt)) | Nr4a2(wt/wt) n = 10 cells, 1 mouse; Nr4a2(del/wt) n = 10 cells, 1 mouse. | Mann-Whitney U test | NA | Sum of ranks (Nr4a2(wt/wt)) = 103, Sum of ranks (Nr4a2(del/wt)) = 107, p-value = 0.9118 |
| Fig. 5k | % change in AUC - Cav2.1&Cav2.2 (Nr4a2(del/wt)/Nr4a2(wt/wt)) | Nr4a2(wt/wt) n = 17 cells, 2 mice; Nr4a2(del/wt) n = 19 cells, 2 mice. | Mann-Whitney U test | NA | Sum of ranks (Nr4a2(wt/wt)) = 385, Sum of ranks (Nr4a2(del/wt)) = 281, p-value = 0.0247 |
| Fig. 5k | % change in AUC - Cav2.3 (Nr4a2(del/wt)/Nr4a2(wt/wt)) | Nr4a2(wt/wt) n = 10 cells, 1 mouse; Nr4a2(del/wt) n = 9 cells, 1 mouse. | Mann-Whitney U test | NA | Sum of ranks (Nr4a2(wt/wt)) = 117, Sum of ranks (Nr4a2(del/wt)) = 73, p-value = 0.1823 |
| Fig. 5k | % change in AUC - Cav3 (Nr4a2(del/wt)/Nr4a2(wt/wt)) | Nr4a2(wt/wt) n = 10 cells, 1 mouse; Nr4a2(del/wt) n = 9 cells, 1 mouse. | Mann-Whitney U test | NA | Sum of ranks (Nr4a2(wt/wt)) = 112, Sum of ranks (Nr4a2(del/wt)) = 78, p-value = 0.3562 |
| Fig. 5k | % change in AUC - Cav2.3&Cav3 (Nr4a2(del/wt)/Nr4a2(wt/wt)) | Nr4a2(wt/wt) n = 8 cells, 1 mouse; Nr4a2(del/wt) n = 9 cells, 1 mouse. | Mann-Whitney U test | NA | Sum of ranks (Nr4a2(wt/wt)) = 68, Sum of ranks (Nr4a2(del/wt)) = 85, p-value = 0.7430 |
| Fig. 5k | % change in AUC - Iberiotoxin (Nr4a2(del/wt)/Nr4a2(wt/wt)) | Nr4a2(wt/wt) n = 19 cells, 2 mice; Nr4a2(del/wt) n = 15 cells, 2 mice. | Mann-Whitney U test | NA | Sum of ranks (Nr4a2(wt/wt)) = 317, Sum of ranks (Nr4a2(del/wt)) = 278, p-value = 0.6074 |
| Fig. 5k | % change in AUC - Ryanodine (Nr4a2(del/wt)/Nr4a2(wt/wt)) | Nr4a2(wt/wt) n = 26 cells, 3 mice; Nr4a2(del/wt) n = 24 cells, 3 mice. | Mann-Whitney U test | NA | Sum of ranks (Nr4a2(wt/wt)) = 739, Sum of ranks (Nr4a2(del/wt)) = 536, p-value = 0.1437 |
| Fig. 5l | Change in AHP Amplitude - ACSF (Nr4a2(del/wt)/Nr4a2(wt/wt)) | Nr4a2(wt/wt) n = 31 cells, 7 mice; Nr4a2(del/wt) n = 43 cells, 10 mice. | Mann-Whitney U test | NA | Sum of ranks (Nr4a2(wt/wt)) = 891, Sum of ranks (Nr4a2(del/wt)) = 1884, p-value = 0.0026 |
| Fig. 5l | Change in AHP Amplitude - Synaptic blockers (Nr4a2(del/wt)/Nr4a2(wt/wt)) | Nr4a2(wt/wt) n = 16 cells, 3 mice; Nr4a2(del/wt) n = 10 cells, 2 mice. | Mann-Whitney U test | NA | Sum of ranks (Nr4a2(wt/wt)) = 179, Sum of ranks (Nr4a2(del/wt)) = 172, p-value = 0.0532 |
| Fig. 5l | Change in AHP Amplitude - VGCC (Nr4a2(del/wt)/Nr4a2(wt/wt)) | Nr4a2(wt/wt) n = 9 cells, 1 mouse; Nr4a2(del/wt) n = 7 cells, 1 mouse. | Mann-Whitney U test | NA | Sum of ranks (Nr4a2(wt/wt)) = 74, Sum of ranks (Nr4a2(del/wt)) = 46, p-value = 0.281 |
| Fig. 5l | Change in AHP Amplitude - Cav2&Cav3 (Nr4a2(del/wt)/Nr4a2(wt/wt)) | Nr4a2(wt/wt) n = 10 cells, 1 mouse; Nr4a2(del/wt) n = 10 cells, 1 mouse. | Mann-Whitney U test | NA | Sum of ranks (Nr4a2(wt/wt)) = 98, Sum of ranks (Nr4a2(del/wt)) = 112, p-value = 0.6305 |
| Fig. 5l | Change in AHP Amplitude - Cav2.1&Cav2.2 (Nr4a2(del/wt)/Nr4a2(wt/wt)) | Nr4a2(wt/wt) n = 17 cells, 2 mice; Nr4a2(del/wt) n = 19 cells, 2 mice. | Mann-Whitney U test | NA | Sum of ranks (Nr4a2(wt/wt)) = 294, Sum of ranks (Nr4a2(del/wt)) = 447, p-value = 0.099 |
| Fig. 5l | Change in AHP Amplitude - Cav2.3 (Nr4a2(del/wt)/Nr4a2(wt/wt)) | Nr4a2(wt/wt) n = 10 cells, 1 mouse; Nr4a2(del/wt) n = 9 cells, 1 mouse. | Mann-Whitney U test | NA | Sum of ranks (Nr4a2(wt/wt)) = 92, Sum of ranks (Nr4a2(del/wt)) = 98, p-value = 0.549 |
| Fig. 5l | Change in AHP Amplitude - Cav3 (Nr4a2(del/wt)/Nr4a2(wt/wt)) | Nr4a2(wt/wt) n = 10 cells, 1 mouse; Nr4a2(del/wt) n = 9 cells, 1 mouse. | Mann-Whitney U test | NA | Sum of ranks (Nr4a2(wt/wt)) = 89, Sum of ranks (Nr4a2(del/wt)) = 101, p-value = 0.4002 |
| Fig. 5l | Change in AHP Amplitude - Cav2.3&Cav3 (Nr4a2(del/wt)/Nr4a2(wt/wt)) | Nr4a2(wt/wt) n = 8 cells, 1 mouse; Nr4a2(del/wt) n = 9 cells, 1 mouse. | Mann-Whitney U test | NA | Sum of ranks (Nr4a2(wt/wt)) = 83, Sum of ranks (Nr4a2(del/wt)) = 88, p-value = 0.8633 |
| Fig. 5l | Change in AHP Amplitude - Iberiotoxin (Nr4a2(del/wt)/Nr4a2(wt/wt)) | Nr4a2(wt/wt) n = 19 cells, 2 mice; Nr4a2(del/wt) n = 15 cells, 2 mice. | Mann-Whitney U test | NA | Sum of ranks (Nr4a2(wt/wt)) = 362, Sum of ranks (Nr4a2(del/wt)) = 379, p-value = 0.4258 |

| | | | | | |
|---|---|---|---|---|---|
| Fig. 5l | Change in AHP Amplitude - Ryanodine (Nr4a2(del/wt)/Nr4a2(wt/wt)) | Nr4a2(wt/wt) n = 26 cells, 3 mice; Nr4a2(del/wt) n = 24 cells, 3 mice. | Mann-Whitney U test | NA | Sum of ranks (Nr4a2(wt/wt)) = 779, Sum of ranks (Nr4a2(del/wt)) = 547, p-value = 0.15 |
| Fig. 6a | number of DAPI+ cells in ROI | Nr4a2(wt/wt) n = 55697 cells, 26 sections, 4 mice; Nr4a2(del/wt) n = 71389 cells, 34 sections, 4 mice. | chi-square-based LRT applied to a negative binomial GLMM with quadratic parametrization | probeset | estimate = -0.02, LRT = 0.78, p-value = 0.376 |
| Fig. 6b | log2 ratio total number Nr4a2+ cells in ROI (Nr4a2(del/wt)/Nr4a2(wt/wt)) | Nr4a2(wt/wt) n = 55697 cells, 26 sections, 4 mice; Nr4a2(del/wt) n = 71389 cells, 34 sections, 4 mice. | chi-square-based LRT applied to a negative binomial GLMM with quadratic parametrization | probeset | estimate = -0.875, LRT = 63.78, p-value = 1.39e-15 |
| Fig. 6c | log2 ratio proportion cell type (Nr4a2(del/wt)/Nr4a2(wt/wt)) | Nr4a2(wt/wt) n = 6851 cells, 5 mice; Nr4a2(del/wt) n = 8001 cells, 8 mice. | moderated t-test with robust empirical Bayes variance shrinkage; p-values were adjusted for multiple comparisons using the Benjamini-Hochberg false discovery rate method | NA | CLA 1: ratio = 0.371, t-value = -3.87, p-value = 0.0244; CLA 2: ratio = 0.333, t-value = -3.37, p-value = 0.0347; pir: ratio = 0.559, t-value = -1.77, p-value = 0.453; mMSN: ratio = 1.33, t-value = 1.74, p-value = 0.453; dMSN: ratio = 1.29, t-value = 1.32, p-value = 0.672; L2/3: ratio = 0.775, t-value = -1.26, p-value = 0.672; iMSN: ratio = 1.34, t-value = 1.13, p-value = 0.708; L5b: ratio = 0.896, t-value = -0.95, p-value = 0.723; sMSN: ratio = 1.58, t-value = 0.93, p-value = 0.723; shell 2: ratio = 1.21, t-value = 0.83, p-value = 0.723; L6a 1: ratio = 1.12, t-value = 0.74, p-value = 0.723; shell 1: ratio = 1.18, t-value = 0.67, p-value = 0.723; L6a 2: ratio = 1.2, t-value = 0.65, p-value = 0.723; L6a 4: ratio = 1.07, t-value = 0.34, p-value = 0.84; L6b: ratio = 0.902, t-value = -0.26, p-value = 0.84; L6a 3: ratio = 0.965, t-value = -0.24, p-value = 0.84; IN: ratio = 0.942, t-value = 0.24, p-value = 0.84; shell 3: ratio = 1.06, t-value = -0.21, p-value = 0.84 |
| Fig. 6f | log2 ratio proportion of cell type (Nr4a2(del/wt)/Nr4a2(wt/wt)) | Nr4a2(wt/wt) n = 42912 cells, 14 sections, 2 mice; Nr4a2(del/wt) n = 45572 cells, 15 sections, 2 mice. | moderated t-test with robust empirical Bayes variance shrinkage; p-values were adjusted for multiple comparisons using the Benjamini-Hochberg false discovery rate method | NA | CLA: ratio = 0.645, t-value = -10.70, p-value = 3.85e-12; Syt17+: ratio = 1.55, t-value = 8.65, p-value = 4.6e-10; Shell: ratio = 0.918, t-value = -0.92, p-value = 0.362 |
| Fig. 7a | percentage of classified cells | Nr4a2(wt/wt) n = 4363 cells, 5 mice; Nr4a2(del/wt) n = 4537 cells, 8 mice. | chi-square-based LRT applied to a logistic regression model; p-values were adjusted for multiple comparisons using the Holm method | NA | actual-predicted cell type pair = CLA-CLA: estimate = -3.66, LRT = 218.89, p-value = 1.27e-48; L6a-CLA: estimate = -21, LRT = 2.29, p-value = 0.912; shell-CLA: estimate = -0.874, LRT = 1.86, p-value = 1; CLA-L6a: estimate = 1, LRT = 0.72, p-value = 1; L6a-L6a: estimate = 0.902, LRT = 1.04, p-value = 1; shell-L6a: estimate = -0.36, LRT = 0.37, p-value = 1; CLA-shell: estimate = 3.82, LRT = 220.44, p-value = 6.51e-49; L6a-shell: estimate = -0.373, LRT = 0.14, p-value = 1; shell-shell: estimate = 0.6, LRT = 1.91, p-value = 1 |

| | | | | | |
|---|---|---|---|---|---|
| Supplementary Fig. 10b | Firing frequency difference resurgent INa enhancer (Nr4a2(del/wt)/Nr4a2(wt/wt)) | Nr4a2(wt/wt) n = 8 cells, 1 mouse; Nr4a2(del/wt) n = 9 cells, 1 mouse. | two-way repeated measure ANOVA; post-hoc testing using Fisher's LSD test with 5% false discovery rate; p-values were adjusted for multiple comparisons using the Benjamini-Hochberg false discovery rate method | NA | [I(pA)]x[Genotype]: F(20,320)=2.553, p-value = 0.0003, [Genotype]:F(1,16)=5.964, p-value = 0.0266. 0pA: p-value>0.9999, 25pA: p-value = 0.9911, 50pA:p-value = 0.7534,75pA:p-value = 0.4538,100pA:p-value = 0.2697, 125pA: p-value = 0.2013, 150pA: p-value = 0.0956, 175pA: p-value = 0.0820, 200pA: p-value = 0.0542, 225pA: p-value = 0.0260, 250pA: p-value = 0.0164, 275pA: p-value = 0.0164, 300pA: p-value = 0.0164, 325pA: p-value = 0.0164, 350pA:p-value = 0.0164, 375pA: p-value = 0.0164, 400pA: p-value = 0.0248, 425pA: p-value = 0.0508, 450pA: p-value = 0.0956, 475pA: p-value = 0.3067, 500pA: p-value = 0.9911 |
| Supplementary Fig. 10c | Firing frequency difference Kv1-4 (Nr4a2(del/wt)/Nr4a2(wt/wt)) | Nr4a2(wt/wt) n = 13 cells, 1 mouse; Nr4a2(del/wt) n = 9 cells, 1 mouse. | two-way repeated measure ANOVA; post-hoc testing using Fisher's LSD test with 5% false discovery rate; p-values were adjusted for multiple comparisons using the Benjamini-Hochberg false discovery rate method | NA | [I(pA)]x[Genotype]: F(20,400)=2.634, p-value = 0.0002, [Genotype]:F(1,20)=0.3217, p-value = 0.5769. 0pA: p-value = 0.9045, 25pA: p-value = 0.9045, 50pA:p-value = 0.9045,75pA:p-value = 0.7932,100pA:p-value = 0.5992, 125pA: p-value = 0.5992, 150pA: p-value = 0.5992, 175pA: p-value = 0.6921, 200pA: p-value = 0.9045, 225pA: p-value = 0.9045, 250pA: p-value = 0.9045, 275pA: p-value = 0.9844, 300pA: p-value = 0.9045, 325pA: p-value = 0.9045, 350pA:p-value = 0.6842, 375pA: p-value = 0.5692, 400pA: p-value = 0.4744, 425pA: p-value = 0.2161, 450pA: p-value = 0.1320, 475pA: p-value = 0.1320, 500pA: p-value = 0.1320 |
| Supplementary Fig. 10d | Firing frequency difference Kv1-4 & Synaptic blocker (Nr4a2(del/wt)/Nr4a2(wt/wt)) | Nr4a2(wt/wt) n = 18 cells, 2 mice; Nr4a2(del/wt) n = 16 cells, 2 mice. | two-way repeated measure ANOVA; post-hoc testing using Fisher's LSD test with 5% false discovery rate; p-values were adjusted for multiple comparisons using the Benjamini-Hochberg false discovery rate method | NA | [I(pA)]x[Genotype]: F(20,640)=3.642, p-value<0.0001, [Genotype]:F(1,32)=2.035, p-value = 0.1634. 0pA: p-value = 0.9193, 25pA: p-value = 0.8804, 50pA:p-value = 0.9193,75pA:p-value = 0.7357,100pA:p-value = 0.3715, 125pA: p-value = 0.1837, 150pA: p-value = 0.0539, 175pA: p-value = 0.0539, 200pA: p-value = 0.0539, 225pA: p-value = 0.0539, 250pA: p-value = 0.0539, 275pA: p-value = 0.1152, 300pA: p-value = 0.1590, 325pA: p-value = 0.1837, 350pA:p-value = 0.4073, 375pA: p-value = 0.5720, 400pA: p-value = 0.4501, 425pA: p-value = 0.7460, 450pA: p-value = 0.4359, 475pA: p-value = 0.2219, 500pA: p-value = 0.1342 |
| Supplementary Fig. 10e | Firing frequency difference Kv7 blocker (Nr4a2(del/wt)/Nr4a2(wt/wt)) | Nr4a2(wt/wt) n = 9 cells, 1 mouse; Nr4a2(del/wt) n = 6 cells, 1 mouse. | two-way repeated measure ANOVA; post-hoc testing using Fisher's LSD test with 5% false discovery rate; p-values were adjusted for multiple comparisons using the Benjamini-Hochberg false discovery rate method | NA | [I(pA)]x[Genotype]: F(20,260)=1.858, p-value = 0.0157, [Genotype]:F(1,13)=1.478, p-value = 0.2457. 0pA: p-value>0.9999, 25pA: p-value>0.9999, 50pA:p-value = 0.9620,75pA:p-value = 0.6023,100pA:p-value = 0.3874, 125pA: p-value = 0.2959, 150pA: p-value = 0.2959, 175pA: p-value = 0.2947, 200pA: p-value = 0.2947, 225pA: p-value = 0.2947, 250pA: p-value = 0.2947, 275pA: p-value = 0.2947, 300pA: p-value = 0.2959, 325pA: p-value = 0.6023, 350pA:p-value = 0.8832, 375pA: |

| | | | | | |
|---|---|---|---|---|---|
| | | | | | p-value = 0.9823, 400pA: p-value = 0.8832, 425pA: p-value = 0.8149, 450pA: p-value = 0.3052, 475pA: p-value = 0.2959, 500pA: p-value = 0.2947 |
| Supplementary Fig. 10f | Firing frequency difference IRK & Synaptic blocker (Nr4a2(del/wt)/Nr4a2(wt/wt)) | Nr4a2(wt/wt) n = 11 cells, 1 mouse; Nr4a2(del/wt) n = 14 cells, 1 mouse. | two-way repeated measure ANOVA; post-hoc testing using Fisher's LSD test with 5% false discovery rate; p-values were adjusted for multiple comparisons using the Benjamini-Hochberg false discovery rate method | NA | [I(pA)]x[Genotype]: F(20,460)=1.796, p-value = 0.0188, [Genotype]:F(1,23)=12.55, p-value = 0.0017. 0pA: p-value<0.9999, 25pA: p-value<0.9999, 50pA:p-value = 0.4390,75pA:p-value = 0.1924,100pA:p-value = 0.0583, 125pA: p-value = 0.0215, 150pA: p-value = 0.0098, 175pA: p-value = 0.0059, 200pA: p-value = 0.0051, 225pA: p-value = 0.0051, 250pA: p-value = 0.0051, 275pA: p-value = 0.0051, 300pA: p-value = 0.0066, 325pA: p-value = 0.0079, 350pA:p-value = 0.0094, 375pA: p-value = 0.0167, 400pA: p-value = 0.0218, 425pA: p-value = 0.0279, 450pA: p-value = 0.0284, 475pA: p-value = 0.0459, 500pA: p-value = 0.0590 |
| Supplementary Fig. 10g | Firing frequency difference Cav2&Cav3 blocker (Nr4a2(del/wt)/Nr4a2(wt/wt)) | Nr4a2(wt/wt) n = 10 cells, 1 mouse; Nr4a2(del/wt) n = 10 cells, 1 mouse. | two-way repeated measure ANOVA; post-hoc testing using Fisher's LSD test with 5% false discovery rate; p-values were adjusted for multiple comparisons using the Benjamini-Hochberg false discovery rate method | NA | [I(pA)]x[Genotype]: F(20,360)=0.2911, p-value = 0.9990, [Genotype]:F(1,18)=0.1492, p-value = 0.7038. 0pA: p-value>0.9999, 25pA: p-value>0.9999, 50pA:p-value>0.9999,75pA:p-value>0.9999,100pA:p-value>0.9999, 125pA: p-value>0.9999, 150pA: p-value>0.9999, 175pA: p-value>0.9999, 200pA: p-value>0.9999, 225pA: p-value>0.9999, 250pA: p-value>0.9999, 275pA: p-value>0.9999, 300pA: p-value>0.9999, 325pA: p-value>0.9999, 350pA:p-value>0.9999, 375pA: p-value>0.9999, 400pA: p-value>0.9999, 425pA: p-value>0.9999, 450pA: p-value>0.9999, 475pA: p-value>0.9999, 500pA: p-value>0.9999 |
| Supplementary Fig. 10h | Firing frequency difference Cav2.1&Cav2.2 blocker (Nr4a2(del/wt)/Nr4a2(wt/wt)) | Nr4a2(wt/wt) n = 17 cells, 2 mice; Nr4a2(del/wt) n = 19 cells, 2 mice. | two-way repeated measure ANOVA; post-hoc testing using Fisher's LSD test with 5% false discovery rate; p-values were adjusted for multiple comparisons using the Benjamini-Hochberg false discovery rate method | NA | [I(pA)]x[Genotype]: F(20,680)=2.727, p-value<0.001, [Genotype]:F(1,34)=0.3671, p-value = 0.0638. 0pA: p-value>0.9999, 25pA: p-value = 0.9599, 50pA:p-value = 0.8194,75pA:p-value = 0.4281,100pA:p-value = 0.1125, 125pA: p-value = 0.0838, 150pA: p-value = 0.0652, 175pA: p-value = 0.0496, 200pA: p-value = 0.0496, 225pA: p-value = 0.0503, 250pA: p-value = 0.0532, 275pA: p-value = 0.0510, 300pA: p-value = 0.0683, 325pA: p-value = 0.1129, 350pA:p-value = 0.2897, 375pA: p-value = 0.4789, 400pA: p-value = 0.5952, 425pA: p-value = 0.5952, 450pA: p-value = 0.9599, 475pA: p-value = 0.5952, 500pA: p-value = 0.5868 |

| | | | | | |
|---|---|---|---|---|---|
| Supplementary Fig. 10i | Firing frequency difference Cav2.3 blocker (Nr4a2(del/wt)/Nr4a2(wt/wt)) | Nr4a2(wt/wt) n = 10 cells, 1 mouse; Nr4a2(del/wt) n = 9 cells, 1 mouse. | two-way repeated measure ANOVA; post-hoc testing using Fisher's LSD test with 5% false discovery rate; p-values were adjusted for multiple comparisons using the Benjamini-Hochberg false discovery rate method | NA | [I(pA)]x[Genotype]: F(20,340)=0.8525, p-value = 0.6483, [Genotype]:F(1,17)=1.678, p-value = 0.2125. 0pA: p-value>0.9999, 25pA: p-value>0.9999, 50pA:p-value>0.9999,75pA:p-value>0.9999,100pA:p-value>0.9999, 125pA: p-value>0.9999, 150pA: p-value>0.9999, 175pA: p-value = 0.9241, 200pA: p-value = 0.6471, 225pA: p-value = 0.4761, 250pA: p-value = 0.3638, 275pA: p-value = 0.3638, 300pA: p-value = 0.3638, 325pA: p-value = 0.3638, 350pA:p-value = 0.3638, 375pA: p-value = 0.3638, 400pA: p-value = 0.9511, 425pA: p-value = 0.6471, 450pA: p-value = 0.8254, 475pA: p-value = 0.6471, 500pA: p-value = 0.6471 |
| Supplementary Fig. 10j | Firing frequency difference Cav3 blocker (Nr4a2(del/wt)/Nr4a2(wt/wt)) | Nr4a2(wt/wt) n = 10 cells, 1 mouse; Nr4a2(del/wt) n = 9 cells, 1 mouse. | two-way repeated measure ANOVA; post-hoc testing using Fisher's LSD test with 5% false discovery rate; p-values were adjusted for multiple comparisons using the Benjamini-Hochberg false discovery rate method | NA | [I(pA)]x[Genotype]: F(20,340)=3.421, p-value<0.0001, [Genotype]:F(1,17)=1.424, p-value = 0.2491. 0pA: p-value>0.9999, 25pA: p-value>0.9999, 50pA:p-value>0.9999,75pA:p-value>0.9999,100pA:p-value = 0.9224, 125pA: p-value = 0.7027, 150pA: p-value = 0.5431, 175pA: p-value = 0.4325, 200pA: p-value = 0.2773, 225pA: p-value = 0.2330, 250pA: p-value = 0.1442, 275pA: p-value = 0.1442, 300pA: p-value = 0.1442, 325pA: p-value = 0.1442, 350pA:p-value = 0.1442, 375pA: p-value = 0.1448, 400pA: p-value = 0.2816, 425pA: p-value = 0.5846, 450pA: p-value>0.9999, 475pA: p-value = 0.8711, 500pA: p-value = 0.2727 |
| Supplementary Fig. 10k | Firing frequency difference Cav2.3&Cav3 blocker (Nr4a2(del/wt)/Nr4a2(wt/wt)) | Nr4a2(wt/wt) n = 8 cells, 1 mouse; Nr4a2(del/wt) n = 9 cells, 1 mouse. | two-way repeated measure ANOVA; post-hoc testing using Fisher's LSD test with 5% false discovery rate; p-values were adjusted for multiple comparisons using the Benjamini-Hochberg false discovery rate method | NA | [I(pA)]x[Genotype]: F(20,300)=1.250, p-value = 0.2118, [Genotype]:F(1,15)=2.340, p-value = 0.1469. 0pA: p-value>0.9999, 25pA: p-value>0.9999, 50pA:p-value>0.9999,75pA:p-value = 0.8179,100pA:p-value = 0.7699, 125pA: p-value = 0.7488, 150pA: p-value = 0.4094, 175pA: p-value = 0.5922, 200pA: p-value = 0.8863, 225pA: p-value = 0.5922, 250pA: p-value = 0.5922, 275pA: p-value = 0.5922, 300pA: p-value = 0.5922, 325pA: p-value = 0.5922, 350pA:p-value = 0.5922, 375pA: p-value = 0.4094, 400pA: p-value = 0.3108, 425pA: p-value = 0.2499, 450pA: p-value = 0.1792, 475pA: p-value = 0.1478, 500pA: p-value = 0.1478 |
| Supplementary Fig. 11 | percentage of classified cells | Nr4a2(wt/wt) n = 4363 cells, 5 mice; Nr4a2(del/wt) n = 4537 cells, 8 mice. | chi-square-based LRT applied to a logistic regression model; p-values were adjusted for multiple comparisons using the Holm method | NA | actual-predicted cell type pair = CLA-CLA: estimate = 0.188, LRT = 2.78, p-value = 0.686; L6a-CLA: estimate = 0.129, LRT = 1.12, p-value = 1; shell-CLA: estimate = 0.126, LRT = 1.25, p-value = 1; CLA-L6a: estimate = -0.253, LRT = 7.65, p-value = 0.0512; L6a-L6a: estimate = -0.143, LRT = 2.25, p-value = 0.686; shell-L6a: estimate = -0.154, LRT = 2.84, p-value = 0.686; CLA-shell: estimate = 0.191, LRT = 2.95, p-value = 0.686; L6a-shell: estimate = |

| | | | | | |
|---|---|---|---|---|---|
| | | | | | 0.102, LRT = 0.72, p-value = 1; shell-shell: estimate = 0.106, LRT = 0.91, p-value = 1 |
